# Proteome-anchored multi-omics of recurrent somatic alterations defines tumour states in cervical cancer and enables state-guided drug repurposing

**DOI:** 10.64898/2026.04.21.719801

**Authors:** Saltiel Hamese, Mutsa Takundwa, Earl Prinsloo, Deepak Balaji Thimiri Govinda Raj

## Abstract

Cervical cancer exhibits substantial molecular heterogeneity, and somatic mutation frequency alone is an incomplete surrogate for functional consequence or therapeutic vulnerability. Here, we present a proteome-anchored bioinformatics framework that integrates genomic, epigenomic, transcriptomic, and proteomic data to infer functionally expressed tumour states and to nominate candidate state-reversing perturbagens. Using matched multi-omics data from the TCGA Cervical Squamous Cell Carcinoma and Endocervical Adenocarcinoma (TCGA-CESC) cohort, recurrently mutated genes were first contextualised within the cervical cancer mutational landscape and prioritised based on regulatory DNA methylation perturbations. For selected programmes, including PIK3CA, TTN, MUC5B, SYNE1, and DST, orthogonal molecular states were independently defined at the genomic, transcriptomic, and epigenomic levels and projected onto the proteome using RPPA-based elastic-net classifiers. These models demonstrated high discriminatory performance across multiple modalities, establishing the feasibility of proteome-level encoding of upstream molecular states. State-specific, directionally signed proteomic signatures were subsequently queried against the L1000 Fireworks Display to identify perturbagens predicted to reverse inferred tumour states. Connectivity mapping revealed convergent vulnerability axes centred on stress-buffering and survival systems rather than single oncogenic drivers, including replication stress tolerance, proteostasis and secretory capacity, checkpoint control, apoptotic buffering, metabolic adaptation, and inflammatory signalling. Among the programmes examined, DST-associated states produced the most statistically robust connectivity, prioritising RTK–mTOR signalling, apoptotic priming, and metabolic–inflammatory modulation as coherent therapeutic hypotheses, while other programmes yielded more moderate signals consistent with hypothesis generation. Collectively, this study establishes a scalable, proteome-anchored multi-omics framework that moves beyond mutation-centric analysis toward functional tumour-state inference and provides a principled route for state-guided therapeutic hypothesis generation in cervical cancer.

## Introduction

### Cervical cancer burden and clinical challenges

Cervical cancer (CC) is the fourth most common cancer in women, making up about 7% of all cases (Cohen et al., 2019). CC is primarily caused by persistent infection with high-risk human papillomavirus (HPV), a common sexually transmitted virus. While most HPV infections are cleared by the immune system, long-term infection can lead to abnormal cell changes that may progress to cancer over 15–20 years, or faster in women with weakened immune systems, such as those with untreated HIV. Factors like the HPV type, immune status, other sexually transmitted infections, smoking, hormonal contraceptive use, and early pregnancies can increase the risk of cancer progression. CC is well-controlled in high-income countries but remains a leading cause of cancer deaths in developing regions (Choi et al., 2023; Cohen et al., 2019). Squamous cell carcinoma (SCC) and adenocarcinoma (ADC) are the primary types, comprising 75–90% and 10–20% of cases, respectively. HPV vaccinations and early screening have reduced CSCC incidence and mortality, but ADC cases are increasing (Cohen et al., 2019; Liu et al., 2024). Current treatment options for CC include surgery, radiation, chemotherapy, and immunotherapy. Radiotherapy is crucial since 20–40% of patients experience recurrence within two years, leading to poor outcomes, with a 10–20% five-year survival rate and increased radioresistance in advanced cases (Xing et al., 2023). Significant disparities in cancer incidence, prevalence, and treatment response exist among racial, ethnic and economic groups. Cancer prevalence and incidence patterns have been observed to be increasing across ethnic and racial groups. Tumour heterogeneity further complicates treatment outcomes **(Figure 1)**, with multi-scale variation across patients (intertumour heterogeneity) and within individual tumours (intratumour heterogeneity) contributing to differences in disease progression, therapeutic response, and survival. Specific genetic subtypes are often linked to racial and ethnic groups, further contributing to disparities in outcomes (Thomas & Peters, 2024). Moreover, studies have shown that Black women with cancer, particularly CC, experience worse survival outcomes than White women, influenced by education, lifestyle factors, diabetes, postmenopausal hormone use, and tumour heterogeneity (Harris et al., 2022). To address the increasing burden of cancer in Africa, especially in low- and middle-income countries in sub-Saharan Africa, well-designed therapeutic interventions are essential (Ntekim and Olopade, 2022; Proietto et al., 2023). Current management is strongly anchored to FIGO stage and histology, with surgery commonly used in early-stage disease and concurrent chemoradiotherapy forming a backbone for locally advanced cancers, yet relapse and treatment-limiting toxicity remain common challenges. Recent phase III trials illustrate that meaningful survival gains can be achieved by modifying standard regimens—such as through short-course induction chemotherapy before chemoradiation in locally advanced cervical cancer and the addition of immune checkpoint blockade to definitive chemoradiotherapy—highlighting both progress and the need for rational patient selection strategies.

**Figure 1.**
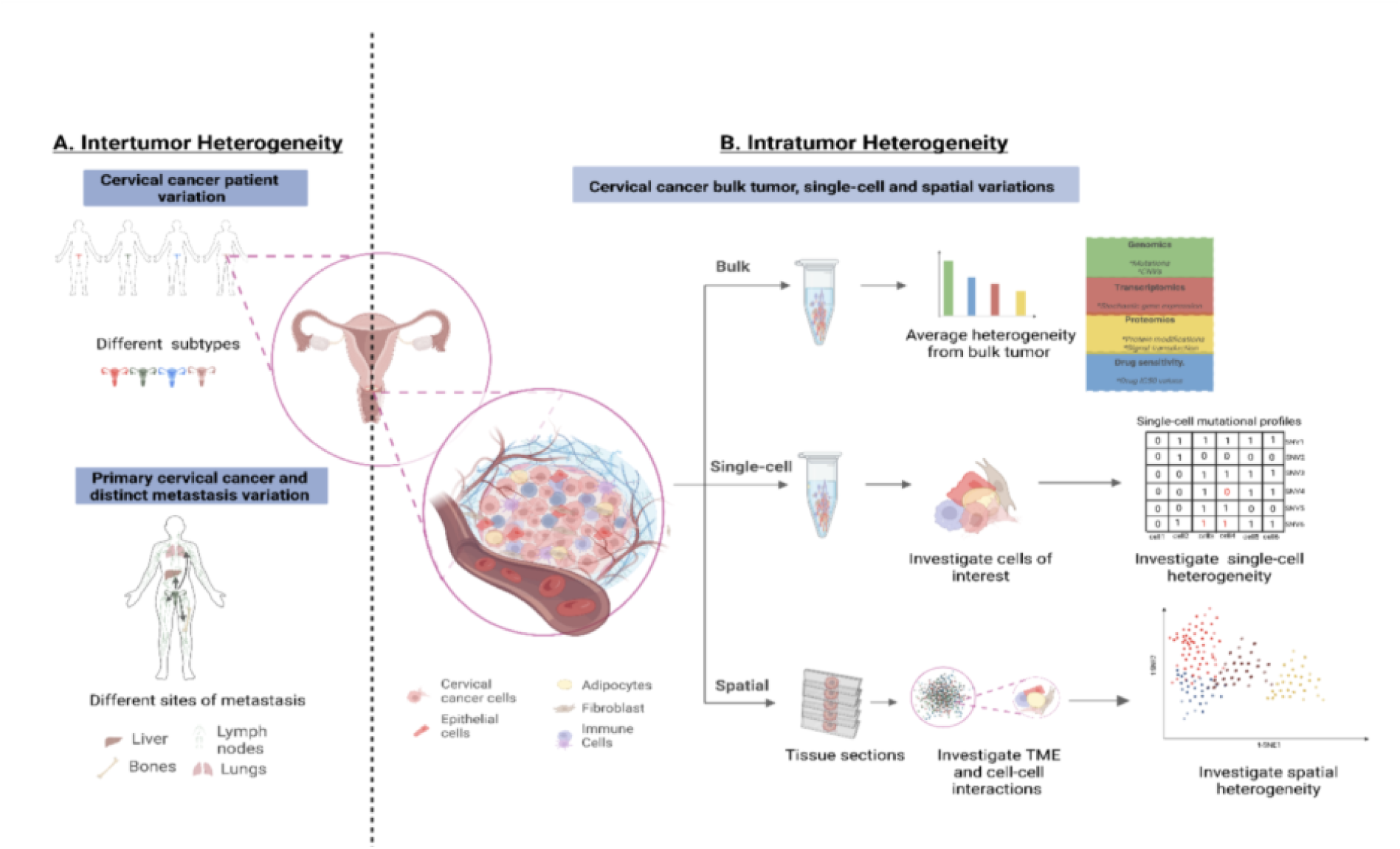
Multi-scale tumour heterogeneity in cervical cancer: from interpatient diversity to intratumour complexity. (A) Intertumour heterogeneity arises from patient-specific variation in cervical cancer subtypes, HPV status, genetic and epigenetic landscapes, and metastatic dissemination patterns, contributing to divergent clinical trajectories, therapeutic responses, and survival outcomes. (B) Intratumour heterogeneity (ITH) reflects dynamic molecular and cellular diversity within individual tumours, driven by clonal evolution, somatic mutation, epigenetic reprogramming, and tumour microenvironment interactions. This complexity can be interrogated across multiple resolutions: bulk profiling captures averaged omics signals, single-cell approaches resolve subclonal architecture and cell-state diversity, and spatial profiling delineates tumour–microenvironment interactions and tissue organisation.

### Somatic mutation–driven tumour heterogeneity and its impact on cancer signalling pathways

Tumour heterogeneity represents a central biological challenge underlying variable disease progression, treatment response, and clinical outcome in cervical cancer. At the population level, intertumour heterogeneity reflects differences between patients in histology, HPV status, molecular subtype, immune contexture, and therapeutic sensitivity. Within individual patients, intratumour heterogeneity (ITH) arises through the accumulation of genetic and epigenetic alterations, clonal evolution, and dynamic interactions between malignant cells and the tumour microenvironment (TME), enabling tumour adaptation, therapeutic resistance, and disease recurrence (Kim et al., 2018). These layered sources of heterogeneity complicate clinical management by driving divergent clinical trajectories even among patients with similar clinicopathological features. Although ITH has been extensively characterised in other solid malignancies such as breast, colorectal, and non-small cell lung cancers, its functional consequences in cervical cancer remain comparatively underexplored, particularly beyond mutation-centric descriptions (Yin et al., 2019; Pellegrina & Vandin, 2022). At the genomic level, this heterogeneity is reflected in the somatic mutation landscape of cervical cancer, which is characterised by both high mutational prevalence and substantial inter-tumour diversity. Comprehensive profiling of 289 primary tumours from the TCGA-CESC cohort revealed that 85% of cases harbour at least one somatic alteration in recurrently mutated genes, underscoring the widespread contribution of genomic instability to tumour heterogeneity. The mutation spectrum is dominated by a limited set of frequently altered genes, including TTN (29%), PIK3CA (28%), KMT2C (19%), MUC16 (17%), KMT2D (13%), FLG (13%), EP300 (12%), DMD (12%), SYNE1 (12%), and FBXW7 (12%). While mutations in large genes such as TTN and MUC16 likely reflect passenger effects associated with gene size, recurrent alterations in PIK3CA, KMT2C/KMT2D, EP300, and FBXW7 are consistent with established oncogenic signalling and chromatin-regulatory drivers. Importantly, most tumours exhibit heterogeneous and largely non-overlapping mutation patterns, with limited co-occurrence among driver genes, highlighting pronounced intertumour genetic diversity. Tumour mutational burden further varies markedly across samples, with a median of 73 variants per tumour and a right-skewed distribution driven by a subset of hypermutated cases. At the variant level, single-nucleotide polymorphisms predominate, with missense mutations representing the most common functional class, followed by nonsense and frameshift alterations. The mutational signature is characterised by a predominance of C>T transitions, consistent with APOBEC-mediated mutagenesis and other context-specific processes active in cervical carcinogenesis.

A critical dimension of this heterogeneity lies in its manifestation through tumour-intrinsic signalling states rather than isolated genomic events. Among recurrently altered genes, SYNE1 was selected purely as an illustrative example to demonstrate the analytical framework, given its intermediate mutation frequency, established roles in nuclear architecture and mechanotransduction, and its ability to exhibit strong proteomic separability despite minimal transcriptomic signal. As illustrated in **Figure 2**, SYNE1-associated multi-omics features are mapped onto canonical oncogenic signalling networks, including ErbB/EGFR-driven receptor tyrosine kinase activation, PI3K–AKT–mTOR signalling, RAF–MEK–ERK/MAPK cascades, and p53-mediated DNA damage and checkpoint responses, alongside immune modulation and metabolic reprogramming modules. Within tumours, distinct subclonal populations may differentially engage these pathways, generating functional heterogeneity in proliferation, survival signalling, checkpoint control, immune evasion, and metabolic adaptation. Importantly, diverse upstream genomic and epigenomic alterations can converge on shared downstream signalling outputs, reflecting functional convergence that obscures genotype–phenotype relationships when considered in isolation. This systems-level view emphasises that tumour heterogeneity is encoded not only in molecular diversity but in the dynamic integration and activation of signalling networks that ultimately govern tumour behaviour. Furthermore, tumour-intrinsic signalling is tightly coupled to microenvironmental context, reinforcing heterogeneity through reciprocal interactions with stromal, immune, and metabolic components of the TME. Growth factor–mediated activation of receptor tyrosine kinases propagates proliferative and survival signals, while replication stress and DNA damage responses converge on checkpoint regulators and apoptotic machinery. Concurrently, metabolic rewiring and immune regulatory pathways enable tumour adaptation, immune escape, and sustained survival under stress conditions. Overlay of elastic-net–derived multi-omics annotations highlights the direction and modality of SYNE1-associated molecular effects across these interconnected pathways (Completed in this study), where node colouring denotes feature-specific coefficients: green, positive expression; red, negative expression; green “Me”, positive methylation; red “Me”, negative methylation; yellow, negative mutation (loss-of-function); and blue, positive mutation (gain-of-function), with “+” and “−” indicating positive and negative coefficients, respectively. Together, these annotations reveal coordinated perturbations across growth-factor signalling, checkpoint control, metabolic adaptation, and immune modulation, providing a systems-level framework for interpreting how SYNE1-associated alterations are functionally encoded within tumour-intrinsic signalling networks and their interaction with the TME. Resolving such multi-layered heterogeneity therefore requires integrative analytical frameworks capable of linking genomic variation to downstream regulatory, transcriptional, and signalling states that ultimately dictate tumour behaviour and therapeutic vulnerability (Grausenburger et al., 2024).

**Figure 2.**
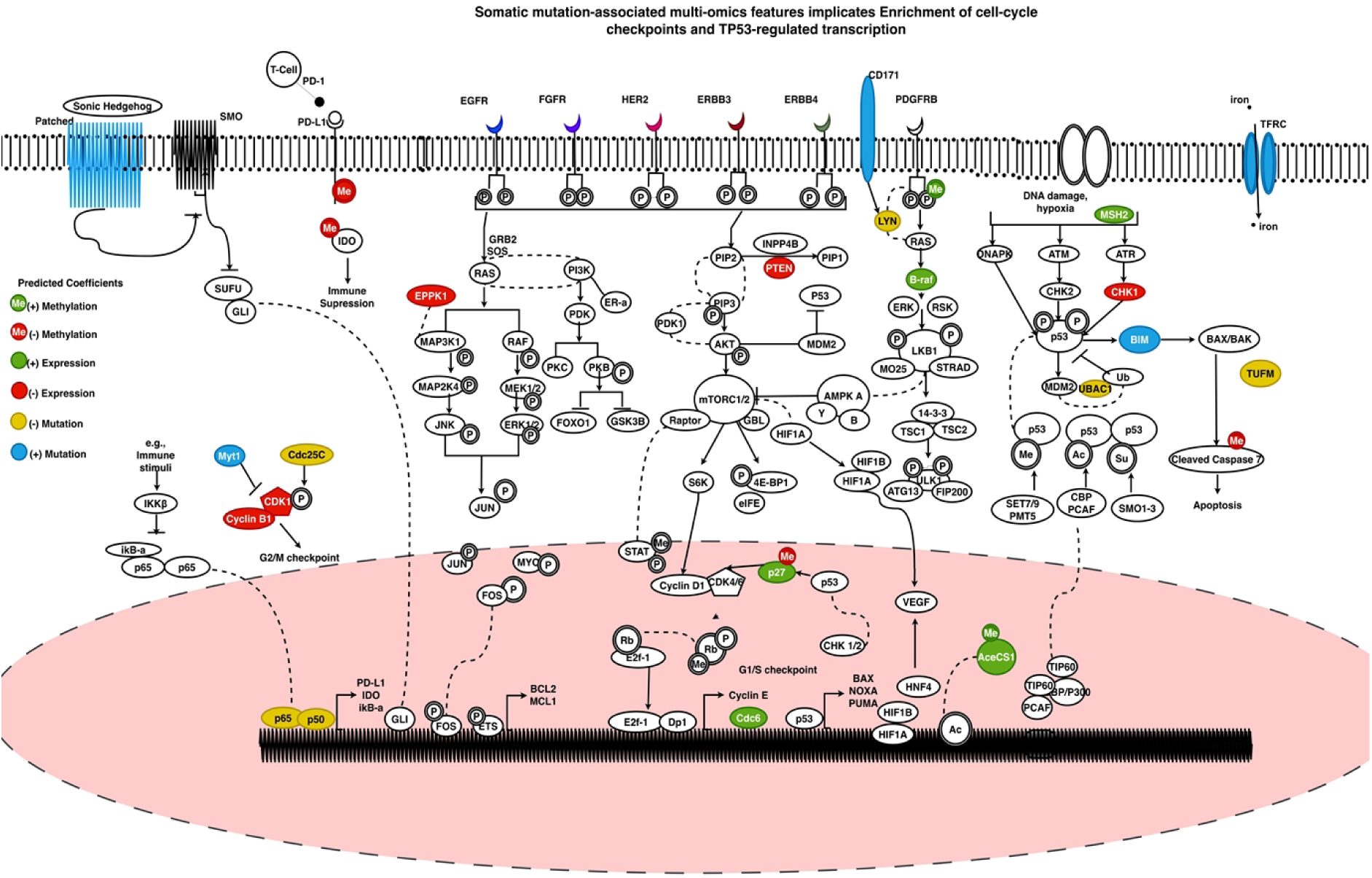
SYNE1 multi-omics features mapped onto canonical cancer signalling pathways within the tumour microenvironment. Comprehensive pathway map illustrating the integration of SYNE1-associated multi-omics features into established oncogenic signalling networks governing tumour cell behaviour and tumour microenvironment (TME) interactions. The schematic captures key signalling axes, including ErbB/EGFR-driven receptor tyrosine kinase activation, PI3K–AKT–mTOR signalling, RAF–MEK–ERK/MAPK cascades, and p53-mediated DNA damage and checkpoint responses, alongside immune modulation and metabolic reprogramming modules. Consistent with current models of tumour biology, these pathways operate within a highly interactive TME, where cancer cells integrate intrinsic oncogenic signals with extrinsic cues derived from stromal, immune, and metabolic components. Growth factor–mediated activation of RTKs propagates proliferative and survival signalling, while replication stress and DNA damage responses converge on checkpoint regulators and apoptotic machinery. Concurrently, metabolic rewiring and immune regulatory pathways contribute to tumour adaptation, immune evasion, and sustained survival under stress conditions. Collectively, this integrated signalling landscape reflects the reciprocal interplay between tumour-intrinsic signalling programmes and microenvironmental influences that drive cancer progression. Overlay of elastic-net–derived multi-omics annotations highlights the direction and modality of SYNE1-associated molecular effects across the network (Completed in this study). Node colouring denotes feature-specific coefficients: green indicates positive expression, red indicates negative expression, green “Me” denotes positive methylation, red “Me” denotes negative methylation, yellow indicates negative mutation (loss-of-function), and blue indicates positive mutation (gain-of-function). The symbols “+” and “−” correspond to positive and negative model coefficients, respectively. These annotations reveal coordinated perturbations across growth-factor signalling, checkpoint control, metabolic adaptation, and immune modulation, providing a systems-level view of how SYNE1-associated alterations are functionally encoded within canonical cancer signalling pathways and the tumour microenvironment.

#### Cervical cancer molecular heterogeneity to proteome-defined functional states

Large-scale integrative profiling has established cervical cancer as a biologically heterogeneous disease shaped by HPV-associated transformation, APOBEC mutagenesis, and diverse recurrent somatic alterations that converge on pathways such as PI3K/RTK signalling and chromatin regulation (Burk et al., 2017). In the TCGA-CESC cohort, multi-omic integration resolved reproducible molecular subtypes spanning squamous and adenocarcinoma-enriched tumours, including an “endometrial-like” subset that is predominantly HPV-negative and enriched for alterations in KRAS, ARID1A and PTEN (Burk et al., 2017). Importantly, mutation recurrence alone is an incomplete proxy for functional relevance: large, frequently mutated genes such as TTN can reflect background mutational processes and tumour mutational burden rather than acting as direct drivers, and TTN mutation counts correlate strongly with TMB across cancers, this motivates analytic frameworks that separate driver-like events from passenger-heavy landscapes and avoid interpreting mutation frequency as a direct proxy for functional mechanism or biological importance (Oh et al., 2020). This motivates analytic frameworks that separate driver-like events from passenger-heavy landscapes and explicitly connect genomic perturbations to propagated pathway effects and clinically actionable cellular states (Boström & Larsson, 2022). A key limitation of mutation-only stratification is that many phenotypes arise through regulatory and signalling rewiring that is not captured by variant calls. Aberrant DNA methylation is a pervasive feature of cancer that remodels regulatory circuitry, constrains transcriptional competence, and can establish stable tumour states independently of coding mutations (Nishiyama & Nakanishi, 2021). In addition, transcript abundance does not reliably predict protein abundance or pathway activity because translation, protein turnover, subcellular localisation, and post-translational modifications can decouple transcriptomic and proteomic layers; large-scale analyses and mechanistic syntheses highlight that mRNA–protein concordance is variable and often modest in complex systems (Liu et al., 2016). These issues are particularly salient in virally driven cancers, where APOBEC activity and host–virus interactions shape mutational signatures and downstream pathway activation in ways that may be more coherently encoded at epigenetic and signalling-protein levels than at bulk mRNA levels (Henderson et al., 2014). Collectively, this supports a network propagation model, in which the functional impact of recurrent alterations is transmitted across interconnected molecular layers—spanning epigenetic regulation, transcriptional output, and proteomic signalling—rather than acting as isolated genomic events. Reverse-phase protein arrays (RPPAs) provide a practical, proteome-centric readout of signalling state by quantifying total and post-translationally modified proteins at scale from limited material, thereby enabling pathway-level inference at the level of functional effectors (Coarfa et al., 2021). RPPA has been deployed across TCGA projects and has proven useful for integrating proteomic features with genomic and transcriptomic data to reveal emergent pathway biology and clinically relevant subtypes across tumour lineages (Akbani et al., 2014). In this context, RPPA is particularly valuable for capturing kinase-network activity, stress-response circuitry, and other mechanistically interpretable programmes that may be weak, fragmented, or non-canonical at the mutation level, thereby nominating therapeutic vulnerabilities more directly than mutation frequency alone. This study is motivated by three convergent needs in cervical cancer research: first, to move beyond mutation frequency as a surrogate for functional importance; second, to model how recurrent somatic alterations propagate across epigenetic, transcriptomic, and proteomic layers; and third, to translate these propagated tumour states into actionable and testable therapeutic hypotheses. Building on TCGA’s comprehensive molecular characterization of cervical cancer and the availability of matched multi-omics data, we develop a bioinformatics framework that (i) contextualizes recurrent somatic alterations within the cervical cancer mutational landscape, (ii) evaluates mutation-associated regulatory rewiring using regulator-focused DNA methylation analysis, (iii) infers functionally expressed tumour states via RPPA-based elastic-net classifiers across orthogonal molecular modalities, and (iv) applies L1000FWD connectivity mapping to nominate repurposing candidates predicted to reverse each classifier-defined state (Burk et al., 2017). By explicitly aligning proteome-informed state inference with perturbational drug-signature reversal, this framework is designed to support downstream state-stratified therapeutic discovery, including the rational prioritization of drug combinations that target complementary stress-buffer systems—such as replication stress tolerance, proteostasis capacity, checkpoint control, and inflammatory survival signalling—grounded in functional proteomic readouts and established connectivity mapping methodology.

## Materials

### Multi-Omics Dataset

Multi-omics data for cervical cancer were obtained from The Cancer Genome Atlas (TCGA), specifically the Cervical Squamous Cell Carcinoma and Endocervical Adenocarcinoma cohort (TCGA-CESC) and processed using a custom integrative pipeline **(Table 1)**. All molecular and clinical datasets were accessed through the Genomic Data Commons (GDC) portal using the Bioconductor package TCGAbiolinks, which provides a reproducible interface for TCGA data querying, downloading, and harmonisation (Colaprico et al., 2016; Grossman et al., 2016). Data retrieval and preparation were performed using the functions GDCquery(), GDCdownload(), and GDCprepare(), enabling explicit control over data categories, types, workflows, and sample filtering. Four molecular layers were retrieved and integrated: **(i)** RNA-Seq transcriptomics, **(ii)** DNA methylation profiling using the Illumina HumanMethylation450 BeadChip, **(iii)** Reverse-Phase Protein Array (RPPA) proteomics, and **(iv)** somatic mutation data in Mutation Annotation Format (MAF), together with matched clinical metadata **(Table 1)**. RNA-Seq data were generated using STAR for alignment and quantification (Dobin et al., 2013) and normalised using the voom framework to support linear modelling of count-based expression data (Law et al., 2014). DNA methylation profiles were generated using the Illumina 450K BeadChip platform (Bibikova et al., 2011). Proteomic measurements were obtained from the MD Anderson Cancer Center RPPA platform (Akbani et al., 2014; Li et al., 2017), and somatic mutation calls were derived from ensemble variant-calling pipelines represented in standardised MAF format (Ellrott et al., 2018; McLaren et al., 2016). Clinical annotations were obtained from TCGA clinical XML files and harmonised GDC clinical tables (Burk et al., 2017). Only primary tumour samples were retained to ensure biological consistency and to minimise confounding effects from recurrent or metastatic disease. A detailed summary of all data sources, experimental platforms, processing workflows, and final analytical formats used in this study is provided in **Table 1**.

**Table 1.**
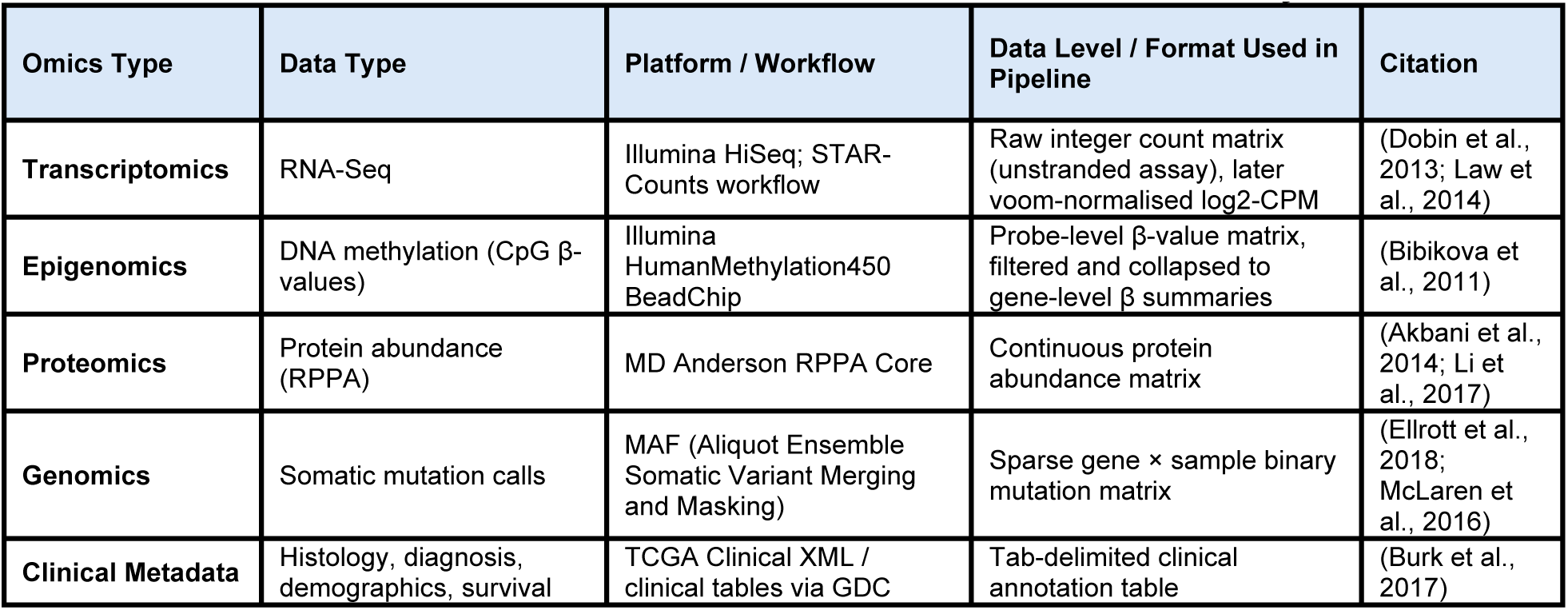
TCGA-CESC data sources and formats as used in this study.

### Molecular Data Pre-processing and Quality Control

Raw molecular datasets were subjected to modality-specific preprocessing and quality control. RNA-Seq preprocessing followed established Bioconductor workflows for count-based modelling using edgeR and limma (Robinson et al., 2010; Law et al., 2014; Ritchie et al., 2015). Lowly expressed genes were removed using filterByExpr(), library sizes were normalised using the Trimmed Mean of M-values (TMM) method, and expression values were transformed to log₂ counts-per-million using the voom method. Quality control included inspection of expression distributions, evaluation of mean–variance relationships, and principal component analysis (PCA) to assess global sample structure. DNA methylation preprocessing incorporated probe-level filtering using annotation resources from minfi (Aryee et al., 2014; Pidsley et al., 2016), removing cross-reactive probes, probes affected by extension base polymorphisms, and probes with >10% missing values. CpG-level measurements were collapsed to gene-level methylation summaries by averaging β-values across annotated probes. Somatic mutation data were processed using maftools (Mayakonda et al., 2018), with high-impact mutations defined as nonsense, frameshift, splice-site, nonstop, and translation start-site variants. Mutation profiles were converted into sparse binary gene × sample matrices. RPPA proteomic measurements were retained as continuous protein abundance values, and antibody identifiers were mapped to canonical gene symbols by resolving RPPA aliases and removing phosphosite suffixes (Akbani et al., 2014; Li et al., 2017). Data manipulation and organisation were performed using the tidyverse ecosystem (Wickham et al., 2019), and graphical outputs were generated using ggplot2 (Wickham, 2016).

### Omics Integration, Modelling, and Statistical Analysis

Processed RNA-Seq expression data, DNA methylation summaries, RPPA proteomic measurements, and somatic mutation matrices were integrated within a unified gene-centric framework. RNA-Seq gene symbols served as the reference backbone onto which gene-level methylation values, mutation indicators, and proteomic measurements were mapped, ensuring alignment of molecular features across all samples. This integration generated a harmonised multi-omics dataset capturing transcriptional, epigenetic, genomic, and proteomic dimensions for each gene, enabling direct cross-layer comparisons and downstream modelling. Recurrently mutated genes (PIK3CA, TTN, MUC5B, SYNE1, and DST) were selected for functional characterisation. For each gene, molecular activation states were defined independently across mutation status, transcriptomic expression, and DNA methylation layers. Differential protein abundance associated with each state was evaluated using linear models with empirical Bayes moderation implemented in limma (Ritchie et al., 2015), generating effect sizes, nominal p-values, and Benjamini–Hochberg false discovery rate (FDR)-adjusted q-values. Statistical significance was primarily assessed at q < 0.05, with interpretation guided by concordance between statistical support, effect size, and biological plausibility. Exploratory principal component analysis (PCA) was used to assess global data structure and identify potential outliers. Predictive modelling was performed using elastic-net regularised regression implemented in glmnet (Friedman et al., 2010), which combines L1 and L2 penalties to enable feature selection in high-dimensional, correlated proteomic datasets. RPPA features with more than 20% missing values were excluded, remaining values were imputed using per-feature medians, and all features were standardised using z-score scaling. Model hyperparameters were optimised via cross-validation, and performance was evaluated using receiver operating characteristic (ROC) curves and area under the curve (AUC) metrics computed with pROC (Robin et al., 2011). Sparse model coefficients were extracted to define proteomic signatures associated with each tumour state.

### Function and Drug-Repurposing Databases

Top proteomic features derived from elastic-net models were subjected to pathway and network-based interpretation. Pathway enrichment analysis was performed using Reactome (Jassal et al., 2020), providing mechanistic annotation of protein modules within each tumour programme. Transcription factor regulatory relationships were inferred using Enrichr-KG (TRRUST v2 library) knowledge graph resources (Evangelista et al., 2023; Han et al., 2018), and protein–protein interaction networks were constructed using STRING to contextualise functional modules and signalling hubs (Szklarczyk et al., 2025). Directionally signed proteomic signatures (derived from model coefficients) were subsequently used for connectivity mapping against the L1000 Fireworks Display (L1000FWD) database to support state-informed therapeutic hypothesis generation in cervical cancer. The L1000CMap paradigm (Subramanian et al., 2017) systematically links genes, drugs, and disease states based on shared or opposing expression signatures. L1000FWD allows interactive visualization and signature similarity searches using up- and down-regulated gene sets, facilitating identification of perturbagens that either mimic or reverse tumour-associated molecular states (Wang et al., 2018). Connectivity scores, Z-scores, nominal p-values, and FDR-adjusted q-values were extracted, and compounds with negative connectivity were prioritised as candidate reversing agents. Results were interpreted using a rank-based framework emphasising consistency and biological coherence rather than strict statistical thresholds, linking inferred tumour states to candidate therapeutic interventions.

## Method

### Multi-omics analysis workflow towards proteomics signatures and candidate therapeutics in cancers

To systematically identify mutation-associated proteomic programmes and potential therapeutic perturbagens in cervical cancer, we implemented a multi-omics analytical framework integrating genomic, epigenomic, transcriptomic, and proteomic layers. The workflow **(Figure 3)** combines data acquisition, rigorous preprocessing, gene-centric multi-omics integration, mutation-driven functional projection, predictive proteomic modelling, and network-based biological interpretation, followed by connectivity mapping for candidate therapeutic discovery. This integrative strategy enables the identification of molecular signatures linking recurrent genomic alterations to downstream protein-level phenotypes and pharmacologically actionable targets.

**Figure 3.**
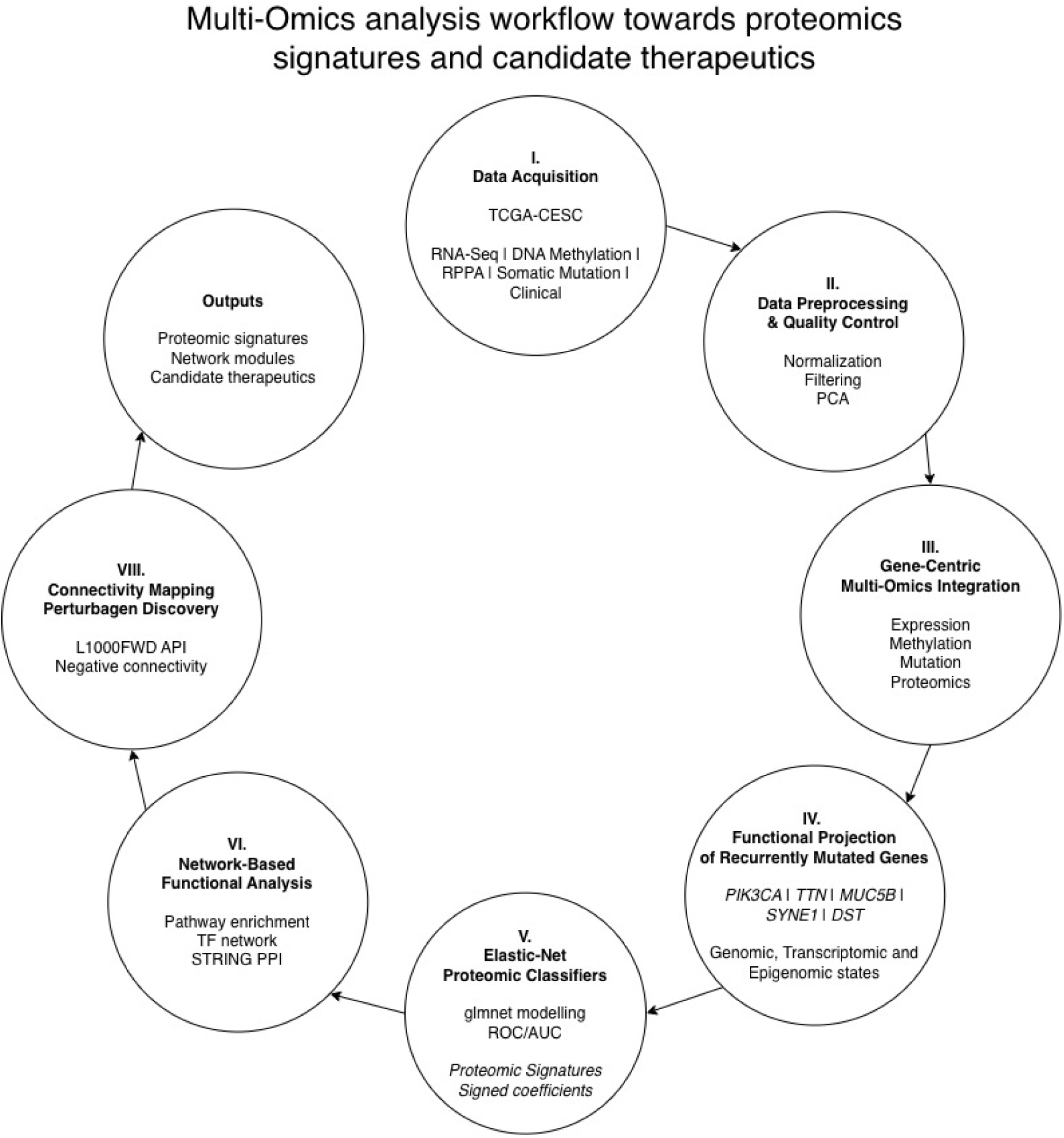
Multi-omics workflow for identifying mutation-associated proteomic signatures and candidate therapeutic perturbagens in cervical cancer. Multi-omic datasets from the TCGA-CESC cohort—including RNA-Seq transcriptomes, DNA methylation profiles, RPPA proteomics, somatic mutation data, and clinical annotations—were integrated to construct a gene-centric analytical framework. (I) Data acquisition and cohort assembly ensured fully matched primary tumour samples across all molecular layers. (II) Molecular datasets underwent modality-specific preprocessing and quality control, including normalization, filtering, and principal component analysis. (III) Gene-centric integration aligned expression, methylation, mutation, and proteomic measurements per gene across the cohort. (IV) Recurrently mutated genes (PIK3CA, TTN, MUC5B, SYNE1, DST) were projected onto transcriptomic, methylomic, and proteomic layers to define mutation-associated protein abundance changes. (V) Elastic-net modelling generated sparse proteomic classifiers predictive of mutation-defined molecular states, with model performance evaluated using ROC/AUC metrics. (VI) Top predictive proteins were functionally characterized via pathway enrichment, transcription factor regulatory analysis, and protein–protein interaction networks. (VII) Directionally signed proteomic signatures were queried against the L1000 Fireworks Display database to identify candidate perturbagens capable of reversing mutation-associated tumour states. This workflow produces interpretable proteomic signatures, network-level functional maps, and prioritized therapeutic compounds.

#### I. Data Acquisition and Cohort Assembly

Multi-omic data for cervical squamous cell carcinoma and endocervical adenocarcinoma from The Cancer Genome Atlas cervical cohort (TCGA-CESC) were obtained through the Genomic Data Commons. Data modalities included RNA-Seq raw integer read counts generated using the STAR-Counts workflow (Transcriptome Profiling), DNA methylation β-values derived from the Illumina HumanMethylation450 BeadChip platform, Reverse Phase Protein Array proteomic measurements produced by the MD Anderson RPPA Core, and somatic mutation data distributed as masked Mutation Annotation Format (MAF) files. Clinical metadata—including histological subtype, demographic variables, HPV annotations, and survival information—were retrieved programmatically using the GDCquery_clinic() function. Sample identifiers were harmonised across all molecular modalities using the first 12 characters of the TCGA barcode to ensure patient-level matching. Samples were subsequently intersected across RNA-Seq, DNA methylation, RPPA proteomics, mutation, and clinical datasets to generate a fully matched multi-omics cohort containing only primary tumour specimens with complete molecular measurements across all layers.

#### II. Molecular Data Preprocessing and Quality Control

Raw molecular datasets consisting of RNA-Seq count matrices, DNA methylation β-value matrices, RPPA protein abundance measurements, and somatic mutation MAF files were subjected to modality-specific preprocessing and quality control. RNA-Seq preprocessing followed established Bioconductor workflows for count-based modelling. Lowly expressed genes were removed using filterByExpr() from edgeR, library sizes were normalised using the Trimmed Mean of M-values (TMM) method, and expression values were transformed to log₂ counts-per-million using the voom method implemented in limma. Quality control included inspection of expression distributions, evaluation of mean–variance relationships, and principal component analysis to assess sample structure. DNA methylation probes were filtered to remove cross-reactive probes, probes influenced by extension base polymorphisms, and probes with more than 10% missing values. CpG-level measurements were subsequently collapsed to gene-level methylation summaries by averaging β-values across annotated probes. Somatic mutation data were processed using maftools, with high-impact mutations defined as nonsense, frameshift, splice-site, nonstop, and translation start-site variants; mutation profiles were converted into binary gene × sample matrices. RPPA proteomic measurements were retained as continuous protein abundance values, and antibody identifiers were mapped to canonical gene symbols by removing phosphosite suffixes and resolving RPPA aliases. These procedures produced quality-controlled molecular datasets consisting of normalised RNA-Seq expression matrices, gene-level methylation summaries, binary mutation matrices, and curated RPPA proteomic profiles.

#### III. Gene-Centric Multi-Omics Integration

Processed RNA-Seq expression data, DNA methylation summaries, RPPA proteomic measurements, and somatic mutation matrices were integrated within a unified gene-centric annotation framework. RNA-Seq gene symbols served as the reference backbone onto which gene-level methylation values, mutation indicators, and RPPA protein measurements were mapped. Alignment across modalities ensured that molecular measurements corresponding to the same gene were synchronised across the matched cohort. This procedure generated an integrated multi-omics gene annotation matrix capturing transcriptional expression, epigenetic methylation status, genomic mutation status, and proteomic representation for each gene across the cohort.

#### IV. Functional Projection of Recurrently Mutated Genes onto the Methylome, Transcriptome, and Proteome

The integrated multi-omics dataset and RPPA proteomic profiles were used to functionally characterise recurrently mutated genes within the TCGA-CESC cohort. Five frequently mutated genes—PIK3CA, TTN, MUC5B, SYNE1, and DST—were selected for modelling. For each gene, molecular activation states were defined independently across three data layers: genomic mutation status, transcriptomic expression state, and gene-level DNA methylation state. Differential protein abundance associated with each molecular state was evaluated using linear models with empirical Bayes moderation implemented in limma. The resulting protein-level signatures associated with mutation-defined molecular states were visualised using volcano plots and used as inputs for downstream modelling analyses.

#### V. Elastic-Net Classifiers

Proteomic matrices derived from RPPA measurements and gene-specific molecular state labels defined by mutation, expression, and methylation layers were used to train elastic-net classifiers. Proteins with more than 20% missing values were removed, remaining missing values were imputed using the per-protein median, and features were standardised using z-score scaling. Predictive models were trained using the glmnet framework, with model hyperparameters (α and λ) optimised through cross-validation. Model performance was evaluated using receiver operating characteristic curves and the area under the curve metric. This procedure yielded sparse proteomic signatures composed of non-zero elastic-net coefficients that represent the proteins most predictive of each mutation-defined molecular state **(Supplementary Figure S0)**.

#### VI. Network-Based Functional Characterisation of Multi-Omics Signatures

Top proteomic features derived from elastic-net models for each mutation-associated programme were subjected to network-based functional interpretation. Pathway enrichment analysis was performed using Enricher-KG with the KEGG 2021 Human, Reactome 2022, Gene Ontology Biological Process, and Human Phenotype Ontology libraries. Transcription factor regulatory networks were inferred using Enrichr knowledge graph resources. Protein–protein interaction networks were subsequently constructed using STRING to visualise functional interaction modules among the multi-omics features. These analyses produced integrated pathway, transcription factor, and protein interaction networks describing the biological organisation of mutation-associated proteomic programmes.

#### VII. Proteome-Guided Connectivity Mapping and Perturbagen Discovery

Directionally signed proteomic signatures derived from elastic-net models were used to identify candidate therapeutic perturbagens through connectivity mapping. Up- and down-regulated gene sets derived from proteomic coefficients were queried against the L1000 Fireworks Display connectivity database. Connectivity scores, Z-scores, p-values, and FDR-adjusted q-values were extracted, and compounds exhibiting negative connectivity were prioritised as candidate reversing perturbagens. This analysis generated ranked lists of therapeutic compounds predicted to counteract mutation-associated tumour states, thereby linking inferred proteomic signatures to potential pharmacological interventions.

## Results

### Data Quality

Across all modalities, quality control indicated high technical consistency, no dominant outliers, and preservation of biologically meaningful structure following preprocessing **(Figure 4)**. RNA-seq normalisation produced aligned global expression distributions and stable mean–variance behaviour consistent with successful TMM scaling and voom precision modelling **(Figure 4A)**. DNA methylation data retained the expected bimodal β-value structure after probe masking and missingness filtering, with no evidence of array-level failure or global bias across samples **(Figure 4B)**. Somatic mutation profiles exhibited canonical TCGA-like characteristics, including predominance of missense variants, enrichment of single-nucleotide substitutions, a right-skewed tumour mutation burden distribution, and recurrent mutation of known large genes and established drivers **(Figure 4C)**. RPPA proteomic data displayed coherent low-dimensional structure without extreme sample outliers, supporting its suitability as the functional response layer for downstream modelling **(Figure 4D)**. Collectively, these diagnostics confirm that subsequent integrative and predictive analyses reflect biological heterogeneity rather than technical artefacts.

**Figure 4.**
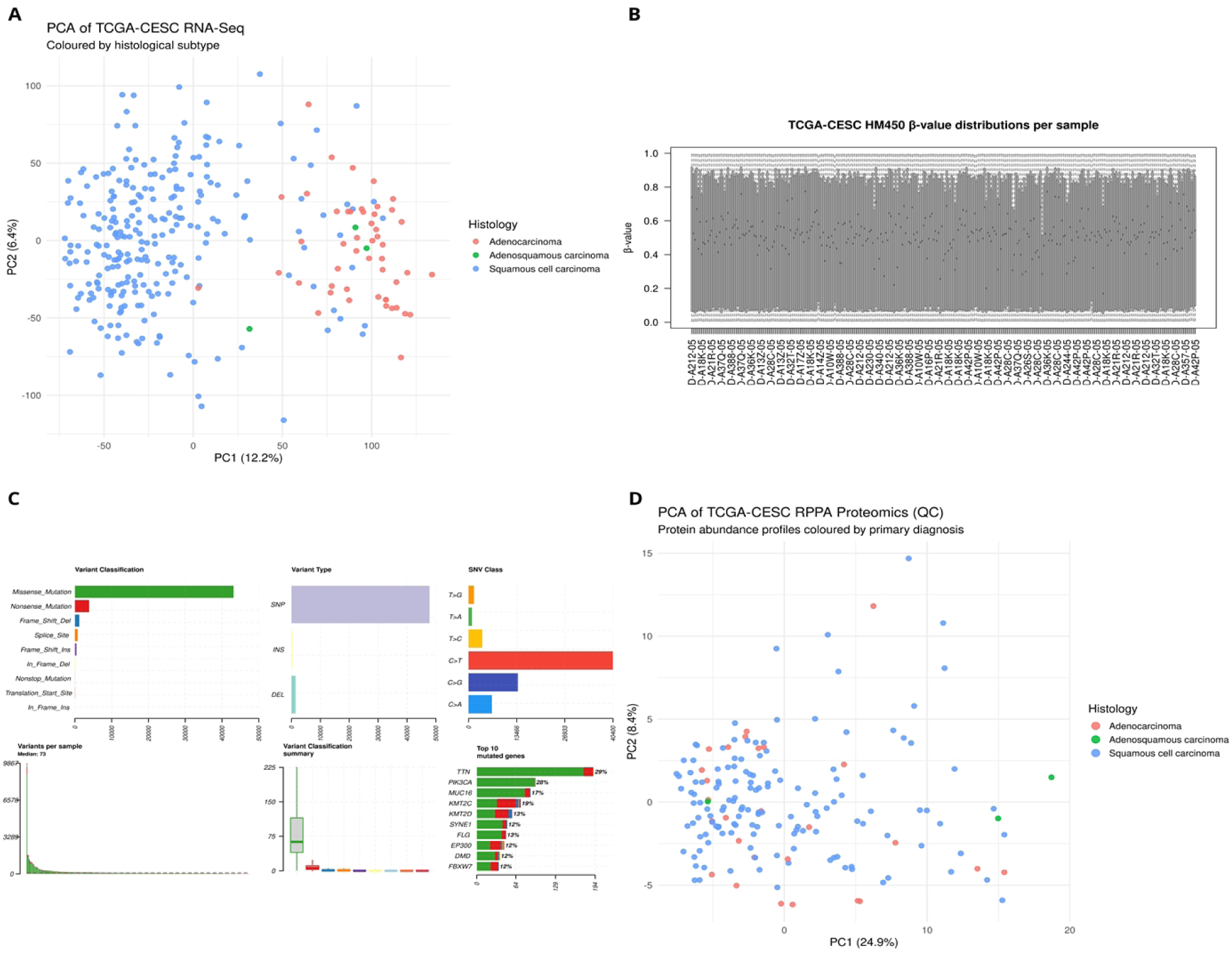
Multi-omics quality control and data integrity in the TCGA-CESC cohort. Quality control diagnostics across transcriptomic, epigenomic, genomic, and proteomic layers demonstrate high technical consistency and preservation of biologically meaningful structure. **(A)** Principal component analysis (PCA) of voom-normalised RNA-seq expression profiles shows coherent low-dimensional structure with partial separation by histological subtype, indicating that biological signal dominates transcriptomic variation following TMM normalisation and variance modelling. **(B)** Distribution of Illumina HM450 DNA methylation β-values per sample after probe masking and missingness filtering, displaying the expected bimodal structure and comparable global profiles across tumours, with no evidence of array-level artefacts. **(C)** Summary of somatic mutation characteristics derived from TCGA-CESC MAF files, including variant classification, variant type, single-nucleotide substitution spectrum, tumour mutation burden per sample, and the most frequently mutated genes, consistent with canonical TCGA mutation patterns. **(D)** PCA of RPPA protein abundance profiles coloured by histological subtype, revealing structured proteomic variation without extreme outliers and supporting RPPA as a robust functional readout for downstream modelling. Collectively, these diagnostics confirm that downstream integrative analyses are driven by biological heterogeneity rather than technical noise.

### Multi-Omics Integration

Following modality-specific normalisation and quality control, all molecular layers were integrated into a unified gene-centric framework that served as the structural foundation for all downstream analyses. This integration produced a master annotation dictionary in which each gene constitutes a single analytical unit annotated across transcriptomic, epigenomic, proteomic, and somatic mutation layers (a detailed description of the unified gene-centric annotation dictionary is provided in **Supplementary Table S1**). For each gene, the dictionary encodes stable genomic identifiers, gene biotype, chromosomal coordinates, and explicit indicators of modality availability, alongside quantitative summaries such as CpG probe coverage, RPPA target representation, and mutation burden metrics. This design enables downstream analyses to distinguish true biological absence from technical non-measurement. DNA methylation provided near-complete gene coverage, with the majority of somatic mutation events mapping to genes represented by CpG probes, whereas RPPA coverage was intentionally sparse and enriched for signalling regulators and pathway hubs, reflecting the targeted nature of the RPPA platform. This asymmetric but complementary modality structure is explicitly encoded in the gene-centric dictionary, with RNA-Seq serving as the annotation backbone for cross-modal alignment. As a result, any gene selected during modelling can be immediately contextualised in terms of its transcriptional measurement, epigenomic coverage, proteomic representation, and mutational status. This unified representation ensures deterministic multi-omics alignment and enables biologically interpretable projection of upstream genomic, transcriptomic, and epigenomic perturbations onto downstream proteomic tumour states.

### Somatic mutation landscape of cervical cancer

Comprehensive somatic mutation profiling was performed on 289 primary cervical cancer tumours from the TCGA-CESC cohort using curated MAF files aligned to GRCh38. Across the cohort, 246 tumours (85%) harboured at least one somatic alteration in the top recurrently mutated genes, underscoring the high mutational prevalence characteristic of cervical cancer. Oncoplot analysis revealed a mutation landscape dominated by a limited set of highly recurrent genes (**Figure 5**). TTN was the most frequently altered gene (29% of samples), followed by PIK3CA (28%), KMT2C (19%), MUC16 (17%), KMT2D (13%), FLG (13%), EP300 (12%), DMD (12%), SYNE1 (12%), and FBXW7 (12%). While alterations in large genes such as TTN and MUC16 are likely enriched due to gene size–associated passenger effects, recurrent mutations in PIK3CA, KMT2C/KMT2D, EP300, and FBXW7 are consistent with established oncogenic signalling and chromatin-regulatory drivers in cervical cancer. Most tumours exhibited heterogeneous mutation patterns with limited co-occurrence among driver genes, highlighting substantial inter-tumour genetic diversity. Tumour mutation burden varied markedly across samples, with a median of 73 variants per tumour and a pronounced right-skewed distribution driven by a subset of hypermutated cases (**Figure 4C**). Single-nucleotide polymorphisms constituted the dominant variant type, with insertions and deletions occurring at substantially lower frequencies. Missense mutations represented the predominant variant classification, followed by nonsense and frameshift events. The single-nucleotide substitution spectrum was dominated by C>T transitions, consistent with known mutational processes active in cervical cancer, including APOBEC-associated mutagenesis. Collectively, these findings define a heterogeneous yet biologically coherent somatic mutation landscape characterised by recurrent alterations in PI3K pathway components and chromatin modifiers, alongside substantial variability in tumour mutation burden.

**Figure 5.**
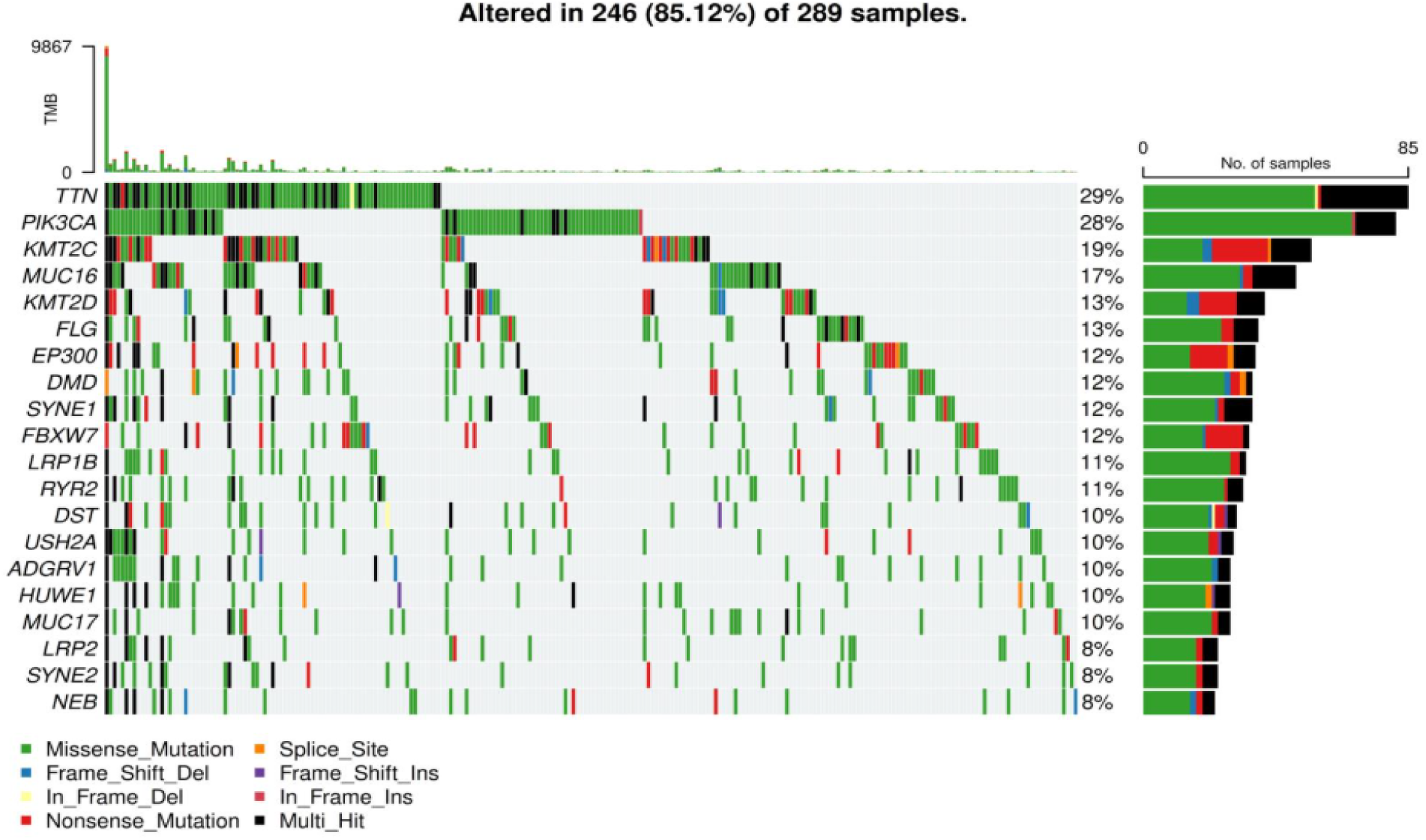
Somatic mutation landscape of TCGA cervical cancer (TCGA-CESC). Oncoplot depicting somatic mutation patterns across **289 primary TCGA-CESC tumours**, restricted to **the top 20 most frequently mutated gene**s. Each column represents an individual tumour sample and each row represents a gene, with mutation types colour-coded by variant classification. The upper barplot shows tumour mutation burden (TMB) per sample, while the right-hand barplot summarises the number and proportion of tumours harbouring mutations in each gene. Overall, **246 of 289 tumours (85.1%)** exhibit at least one alteration in these recurrently mutated genes, highlighting the pervasive somatic mutational burden in cervical cancer.

### Epigenomic consequence of recurrent somatic mutations in cervical cancer

To characterise the epigenetic propagation of recurrent somatic mutations prior to proteomic modelling, we performed regulator-restricted differential DNA methylation analyses, focusing on CpGs mapping to curated, gene-specific regulatory networks. This targeted approach was designed to capture pathway-centred epigenetic remodelling rather than diffuse genome-wide methylation shifts. Across all programmes, significant mutation-associated CpGs were sparse and highly structured, underscoring gene-specific modes of epigenetic coupling **(Supplementary Table S2)**. For PIK3CA, regulator-restricted analysis revealed the most robust and coherent epigenetic signal. Among 129 CpGs tested across 307 tumours (80 mutant, 227 wild type), 17 CpGs reached False Discovery Rate (FDR) significance, with effects strongly concentrated in PIK3R1 and PIK3CD. Multiple probes (e.g. cg25008444, cg06445944, cg01239651, cg09101894, cg25091228, cg16512854) showed consistent negative logFC values, indicating relative hypomethylation in mutant tumours. This directional consistency supports selective epigenetic activation of upstream PI3K regulatory components, rather than global methylome disruption. In contrast, TTN mutation status showed minimal epigenetic perturbation. Of 743 CpGs examined, only a single CpG (cg18127003 in GATA4) reached FDR significance, supporting the interpretation of TTN as largely epigenetically inert despite its high mutation frequency. For SYNE1, a moderate but structured epigenetic signature was observed. Among 1,347 CpGs tested, eight CpGs reached FDR significance, mapping to genes involved in cytoskeletal dynamics and chromatin regulation (RHOA, ITGA6, ACTA1, ACTB, ARID1B). All significant CpGs were hypomethylated in mutant tumours, consistent with coordinated modulation of mechanotransduction and nuclear–cytoskeletal coupling pathways. DST mutation status exhibited a highly focal epigenetic effect. Despite testing 1,309 CpGs, only three CpGs were significant, all mapping to TUBB and showing positive logFC values, indicating hypermethylation in mutant tumours. This suggests selective microtubule-associated regulatory perturbation rather than broad epigenetic reprogramming. Finally, MUC5B-mutant tumours displayed a biologically coherent but focal signature. Five CpGs reached FDR significance among 1,049 tested, mapping to IL1B, IL6, EGFR, GATA6, and ARID1B, all showing hypomethylation. These loci implicate inflammatory signalling, epithelial lineage regulation, and chromatin accessibility, aligning with mucin-associated tumour plasticity. Collectively, these analyses reveal marked heterogeneity in mutation-associated epigenetic propagation. PIK3CA and MUC5B show pathway-coherent remodelling, SYNE1 displays structured mechanotransduction-linked changes, whereas TTN and DST exhibit minimal or highly focal effects **(Supplementary Table S2)**. These findings provide a mechanistic basis for prioritising downstream proteomic modelling and tumour-state inference across programmes. A summary of regulator-focused differential methylation stratified by mutation status is provided in **Table 2**, including CpG counts, sample composition, significant hits (FDR < 0.05), directionality, and biological interpretation. Analyses were restricted to tumours with matched methylation data (n = 244), ensuring consistency with downstream integrative modelling. Within this subset, the mutational landscape remained broadly representative of TCGA-CESC, with TTN (35%) and PIK3CA (33%) as the most frequently altered genes, alongside recurrent mutations in chromatin regulators (KMT2C, KMT2D, EP300) and structural genes (MUC16, FLG, DMD). Notably, key genes including TTN, PIK3CA, SYNE1, DST, and MUC5B harboured at least one significant CpG and are highlighted in the oncoplot **(Supplementary Figure S3)**, marking loci of convergent genetic and epigenetic alteration.

**Table 2.** Epigenetic alterations associated with recurrent somatic mutations in the cervical cancer cohort.

| Mutation | Regulators tested (CpGs) | Samples (mut / wt) | Significant CpGs (FDR < 0.05) | Dominant genes implicated | Direction of effect (mut vs wt) | Interpretation |
| --- | --- | --- | --- | --- | --- | --- |
| **PIK3CA** | 129 | 80 / 227 | 17 | *PIK3R1*,<br>*PIK3CD* | Hypomethylation | Strong, pathway-centered epigenetic rewiring of PI3K regulatory network |
| **TTN** | 743 | 84 / 223 | 1 | *GATA4* | Hypomethylation | Minimal signal; consistent with passenger behaviour |
| **SYNE1** | 1,347 | 35 / 272 | 8 | *RHOA*, *ITGA6*,<br>*ACTA1*,<br>*ARID1B* | Hypomethylation | Coordinated modulation of mechanotransduction and chromatin architecture |
| **DST** | 1,309 | 29 / 278 | 3 | *TUBB* | Hypermethylation | Highly focal microtubule-associated regulatory effect |
| **MUC5B** | 1,049 | 21 / 286 | 5 | *IL1B*, *IL6*,<br>*EGFR*,<br>*GATA6*,<br>*ARID1B* | Hypomethylation | Inflammation- and epithelial-state-linked epigenetic perturbation |

\* Across all programmes, significant mutation-associated CpGs were sparse and highly structured, underscoring gene-specific modes of epigenetic coupling (Supplementary Table S2).

### Transcriptomic consequences of recurrent somatic mutations in cervical cancer

#### Transcriptomic consequences of MUC5B mutation are limited and non-canonical

To assess whether MUC5B mutation is associated with broad transcriptional reprogramming, we performed genome-wide differential expression analysis comparing MUC5B-mutant tumours (n = 21) with wild-type cases (n = 283) using RNA-seq data from the TCGA-CESC cohort. Across 20,644 genes tested, only 22 genes met the significance threshold (FDR < 0.05), indicating a relatively modest and fragmented transcriptional footprint associated with MUC5B mutation (**Supplementary Table S4**). The majority of significant transcripts were upregulated in MUC5B-mutant tumours and were dominated by long non-coding RNAs and poorly characterised loci (including LINC02432, AC112206.3, AC011632.1, and related intergenic transcripts), alongside a limited set of protein-coding genes such as MYO18B, SPDYC, SLC5A12, POU6F2, PON1, and MAGEA4. Notably, only a single gene (ITPKA) exhibited significant downregulation in the mutant group, underscoring the strong directional bias toward transcript upregulation but the absence of a balanced, pathway-coherent transcriptional response. Importantly, the differentially expressed genes identified do not converge on a single signalling pathway or regulatory programme and include multiple loci with limited functional annotation, suggesting that the observed transcriptional changes likely reflect indirect or context-dependent effects rather than direct execution of a MUC5B-centred regulatory network. To further test whether MUC5B mutation preferentially perturbs a predefined regulatory circuitry, we repeated differential expression analysis using a restricted gene set comprising 44 curated MUC5B-associated regulators. In contrast to the genome-wide analysis, no genes within this targeted set were significantly differentially expressed (FDR < 0.05), despite identical sample composition and statistical thresholds. Together, these analyses indicate that MUC5B mutation is not associated with widespread, pathway-coherent transcriptional deregulation at the bulk RNA-seq level. The limited number of significant genes, their heterogeneous functional annotations, and the absence of signal within the curated regulatory set support the interpretation that MUC5B-associated tumour biology is not primarily encoded at the level of steady-state mRNA abundance. This finding provides a clear rationale for downstream proteome-anchored modelling, motivating the use of RPPA-based classifiers and connectivity mapping to capture post-transcriptional, signalling, and stress-adaptive tumour states that are not apparent from transcriptomic profiling alone.

#### PIK3CA mutation is associated with broad transcriptional reprogramming and selective rewiring of PI3K pathway components

To characterise the transcriptomic consequences of PIK3CA mutation prior to proteomic modelling, we performed genome-wide differential expression analysis comparing PIK3CA-mutant tumours (n = 79) with wild-type cases (n = 225) in the TCGA-CESC cohort. Across 20,644 genes tested, 387 genes were significantly differentially expressed at FDR < 0.05, indicating a substantial transcriptional footprint associated with PIK3CA mutation. The most strongly altered transcripts are summarised in **Table 3**. Both up- and down-regulated genes were observed, reflecting broad regulatory reprogramming rather than activation of a single linear pathway. Among the most significantly upregulated genes in PIK3CA-mutant tumours were MISP3, SORT1, KDM4B, AK9, RMND5B, RAB11FIP4, and MAML3, implicating altered cytoskeletal organisation, chromatin modification, vesicular trafficking, and signalling modulation. Conversely, prominent downregulated genes included MPP3, BEX3, C1R, EMC4, CAVIN1, TCEAL9, and PWP1, highlighting suppression of membrane scaffolding, immune complement components, and transcriptional regulatory factors **(Table 3)**. The coexistence of these opposing transcriptional shifts suggests that PIK3CA mutation is associated with coordinated redistribution of cellular resources across signalling, structural, and regulatory programmes rather than uniform pathway upregulation. To determine whether PIK3CA mutation selectively perturbs its canonical signalling circuitry, we next performed differential expression analysis restricted to a curated set of six PI3K pathway regulators. Within this focused analysis, three genes were significantly differentially expressed at FDR < 0.05 **(Supplementary Table S5)**. Notably, PIK3R1 was significantly upregulated in PIK3CA-mutant tumours, whereas PIK3CD and PIK3C3 were downregulated. This pattern indicates compensatory rebalancing of PI3K complex composition and signalling output rather than simple amplification of catalytic activity. Specifically, increased expression of the regulatory subunit PIK3R1, coupled with suppression of alternative catalytic or trafficking-associated components, supports a model in which oncogenic PIK3CA mutation induces adaptive transcriptional rewiring within the PI3K network itself. These findings are consistent with structured pathway-level modulation and provide a mechanistic bridge between widespread transcriptional reprogramming and the distinct proteomic states observed downstream. Together, these transcriptomic analyses demonstrate that PIK3CA mutation is associated with widespread and structured transcriptional reprogramming, in contrast to the minimal and non-canonical RNA-level effects observed for MUC5B mutation. Importantly, the combination of global transcriptional shifts and targeted regulatory rebalancing within the PI3K pathway provides a biologically grounded rationale for downstream RPPA-based multi-state classification. These results support the hypothesis that proteomic tumour states associated with PIK3CA mutation reflect both broad transcriptional context and post-transcriptional integration, motivating subsequent functional projection to the proteome and connectivity-based therapeutic inference.

**Table 3.**
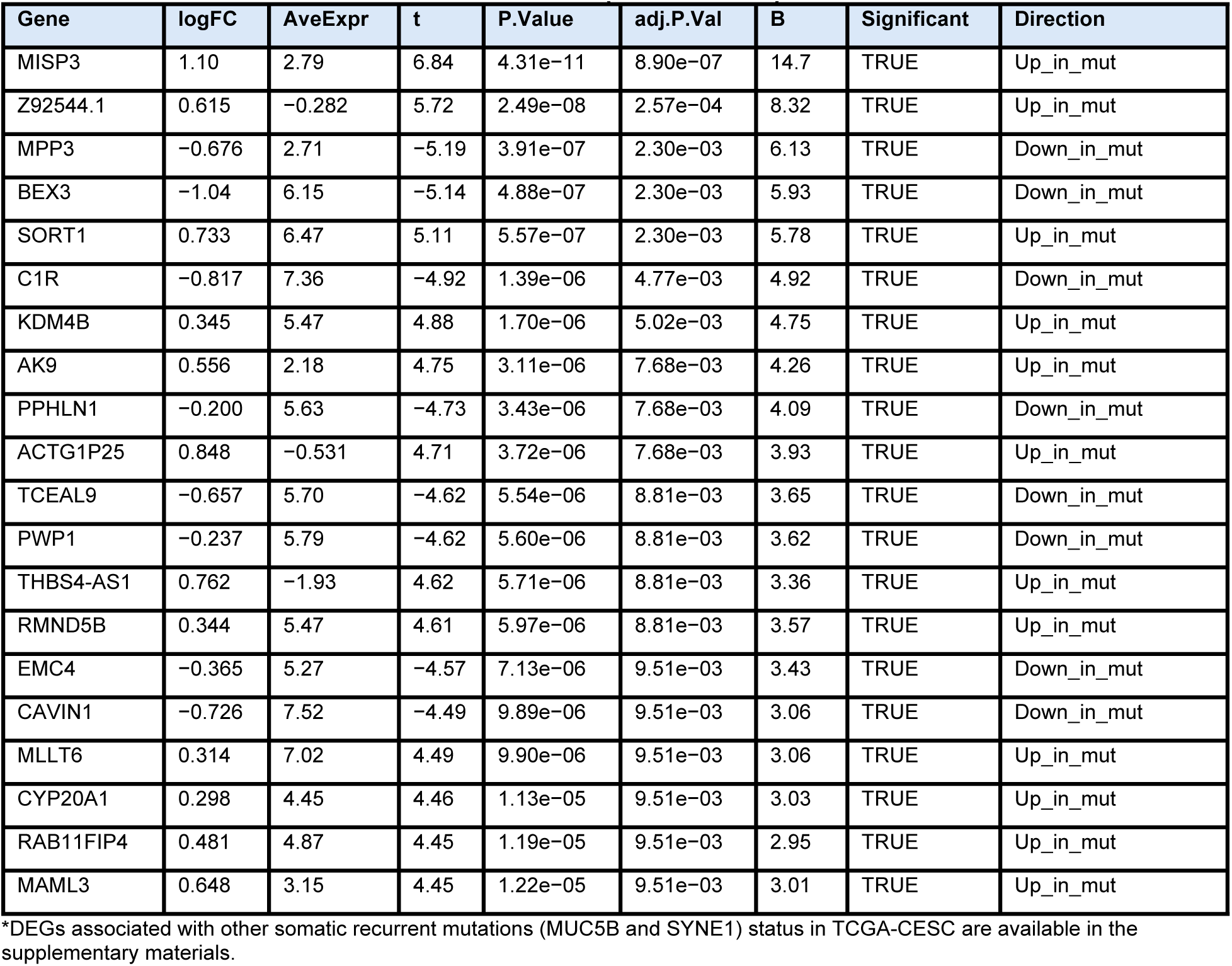
Genome-wide differentially expressed genes (DEGs) associated with PIK3CA mutation status in TCGA-CESC (FDR < 0.05)

#### SYNE1 mutation is associated with minimal transcriptional disruption and absence of canonical regulatory rewiring

To evaluate the transcriptomic impact of SYNE1 mutation prior to proteomic modelling, we performed genome-wide differential expression analysis comparing SYNE1-mutant tumours (n = 35) with wild-type cases (n = 269). Across 20,644 genes tested, only a single gene met the significance threshold (FDR < 0.05). This gene, EPOP, was modestly but significantly upregulated in SYNE1-mutant tumours (logFC = 0.96, adj. P = 0.0085). No additional transcripts exhibited statistically significant differential expression, indicating an exceptionally limited RNA-level footprint associated with SYNE1 mutation. To assess whether SYNE1 mutation selectively perturbs its putative regulatory network, we repeated differential expression analysis using a restricted gene set comprising 41 curated SYNE1-associated regulators. In contrast to the genome-wide analysis, no genes within this targeted set were differentially expressed at FDR < 0.05, despite adequate sample size and identical statistical thresholds. This absence of signal suggests that SYNE1 mutation does not induce coherent transcriptional rewiring of its canonical regulatory circuitry. Collectively, these findings demonstrate that SYNE1 mutation is largely transcriptionally silent in bulk RNA-seq data, despite its strong and highly separable proteomic signature observed in downstream RPPA classifiers. This dissociation between RNA- and protein-level effects supports a model in which SYNE1 alteration primarily influences tumour biology through post-transcriptional, structural, or signalling-mediated mechanisms, rather than through direct modulation of steady-state gene expression. This provides a strong biological rationale for proteome-centric modelling and connectivity mapping to capture SYNE1-associated tumour phenotypes that are invisible at the transcriptomic level.

#### TTN and DST mutations are transcriptionally silent at the bulk RNA-seq level

To determine whether mutations in large cytoskeletal and structural genes are associated with detectable transcriptional consequences, we performed genome-wide and regulator-restricted differential expression analyses for TTN and DST using bulk RNA-seq data. For TTN, comparison of mutant (n = 82) and wild-type tumours (n = 222) across 20,644 genes tested identified no significantly differentially expressed genes at FDR < 0.05. Similarly, analysis restricted to a curated set of 22 TTN-associated regulators yielded no significant transcriptional changes, indicating an absence of coherent RNA-level perturbation attributable to TTN mutation. An analogous pattern was observed for DST, another large cytolinker gene. Genome-wide differential expression analysis comparing DST-mutant (n = 29) and wild-type tumours (n = 275) again identified no significant differentially expressed genes following multiple-testing correction. Restricting the analysis to 43 curated DST-associated regulators likewise revealed no transcriptional signal, despite adequate cohort size and identical statistical thresholds. Together, these results demonstrate that TTN and DST mutations are transcriptionally silent in bulk RNA-seq data, both globally and within their putative regulatory networks. This lack of RNA-level signal contrasts sharply with the strong and highly structured proteomic signatures associated with these genes observed in downstream RPPA classifiers. The dissociation between transcriptional and proteomic effects supports a model in which alterations in TTN and DST primarily influence tumour biology through post-transcriptional, structural, or signalling-mediated mechanisms, rather than through direct modulation of steady-state mRNA abundance. These findings further justify the use of proteome-centric modelling to capture tumour phenotypes driven by large structural gene alterations that are invisible to transcriptome-based analyses.

### Proteomic consequences of recurrent somatic mutations in cervical cancer

In MUC5B-mutant tumours, a distinct and biologically interpretable proteomic profile was observed **(Figure 6A)**. Several proteins exceeded stringent significance thresholds, including ACSS1, ERα_pS118, and CHD1L, implicating coordinated metabolic reprogramming, oestrogen receptor signalling, and chromatin remodelling. These features are consistent with the biology of mucin-associated loci as enhancer-rich and inflammation-responsive, suggesting that MUC5B mutation status reflects broader transcriptional and epigenetic state shifts rather than isolated mucin dysregulation. In contrast, DST-mutant tumours exhibited a strong and structured proteomic signature **(Figure 6B)**. Multiple proteins surpassed both statistical and effect-size thresholds, including EGFR_pY1173, PD-L1, FASN, and PAI1. The enrichment of EGFR phosphorylation and immune checkpoint signalling indicates activation of growth-factor and immune evasion pathways, while increased FASN abundance implicates altered lipid metabolism. Given DST’s role in cytoskeletal organisation and epithelial integrity, these findings support a model of adhesion disruption coupled with compensatory activation of proliferative, metabolic, and immune-modulatory programmes, representing one of the most coherent mutation-associated proteomic states. SYNE1-mutant tumours displayed an intermediate phenotype **(Figure 6C)**. While most proteins did not reach significance, a consistent subset showed directional changes, most notably increased NF-κB_pS536, indicating activation of inflammatory signalling pathways. This pattern aligns with SYNE1’s role in nuclear architecture and mechanotransduction, where structural perturbation preferentially impacts stress-responsive signalling rather than core proliferative or metabolic processes. By contrast, TTN-mutant tumours showed a largely stable proteomic landscape **(Figure 6D)**, with most proteins clustering near zero log₂ fold change and failing to meet statistical or effect-size thresholds. Only a small number of proteins, including ZAP70 and HMHA1, showed modest increases. This sparse and immunologically skewed profile supports the interpretation of TTN as a passenger mutation, reflecting mutation burden or immune context rather than driving a coherent proteomic programme. Collectively, these analyses demonstrate that not all recurrently mutated genes exert equivalent proteomic effects **(Figure 6)**. While TTN mutations are largely proteomically silent, mutations in genes such as DST and MUC5B are associated with coordinated and biologically interpretable programmes spanning growth-factor signalling, immune regulation, metabolism, and chromatin remodelling. These findings support the integration of regulatory methylation prioritisation with proteomic profiling as a strategy to distinguish functional tumour-state alterations from mutation-burden artefacts and to prioritise pathways for mechanistic and therapeutic investigation.

**Figure 6.**
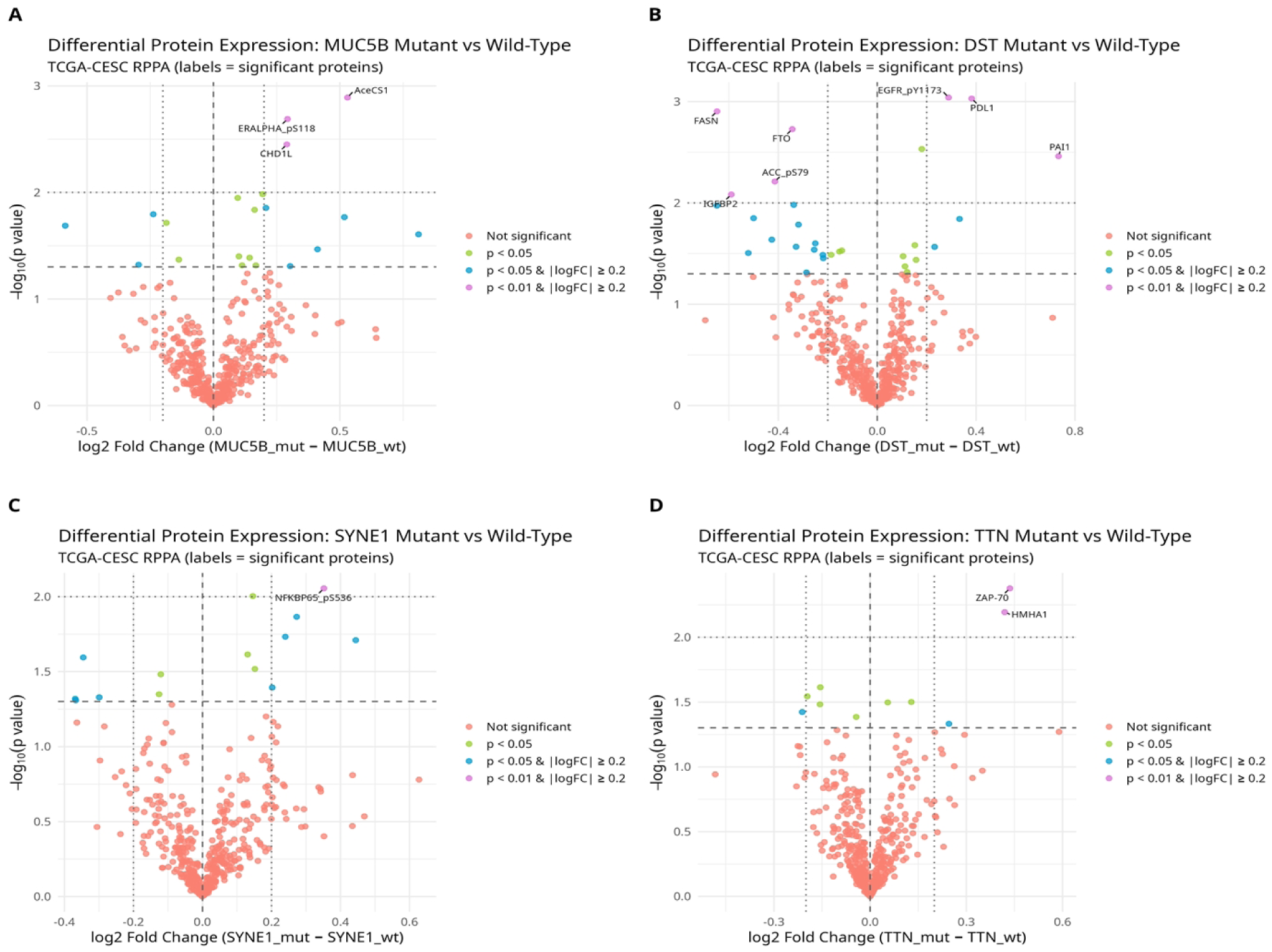
Proteomic consequences of recurrent somatic mutations in cervical cancer. Volcano plots showing differential protein abundance measured by RPPA between mutant and wild-type tumours in the TCGA-CESC cohort for four recurrently mutated genes. **(A)** MUC5B, **(B)** DST, **(C)** SYNE1, and **(D)** TTN. For each comparison, the x-axis denotes the log₂ fold change in protein abundance (mutant minus wild type), and the y-axis denotes −log₁₀(p value) from limma empirical Bayes–moderated linear models. Horizontal dashed lines indicate the nominal significance threshold (p = 0.05), and vertical dashed lines indicate the effect-size threshold (|log₂ fold change| ≥ 0.2). Proteins are coloured by statistical significance and effect size, and selected significantly altered proteins are labelled. These analyses reveal gene-specific and heterogeneous proteomic signatures associated with mutation status, ranging from coordinated signalling and metabolic alterations (e.g. MUC5B and SYNE1) to more focal and sparse effects consistent with passenger-like behaviour (e.g. TTN), highlighting distinct modes by which recurrent somatic mutations are functionally propagated to the proteome in cervical cancer.

#### Proteomic propagation of orthogonal *PI3K* states and epigenetic remodelling of *PI3K* regulator

To illustrate how distinct molecular layers encode PI3K pathway activity, we examined the proteomic and epigenomic consequences of three orthogonal PIK3CA states—genomic mutation, transcriptional activation, and DNA methylation—using TCGA-CESC data (**Figure 7A–D**). Differential protein abundance analysis using RPPA revealed that these states generate markedly different downstream proteomic footprints, indicating hierarchical propagation of PI3K activity across molecular layers. As shown in **Figure7A**, comparison of PIK3CA-mutant versus wild-type tumours (32 mutant vs 140 wild-type) revealed a largely sparse proteomic response. The majority of proteins clustered tightly around log₂ fold change ≈ 0 and did not exceed nominal significance thresholds, indicating that PIK3CA mutation alone does not globally reprogram protein abundance. Nevertheless, a small subset of proteins showed reproducible directional shifts. Fibronectin and RB were among the strongest positive-effect features in mutant tumours, whereas Cyclin E1 and Granzyme B exhibited negative shifts. These selective changes implicate extracellular matrix remodelling, cell-cycle regulation, and immune effector activity, suggesting that genomic PI3K activation produces focused rather than global proteomic perturbations. In contrast, stratification by PIK3CA transcriptional state produced a substantially stronger and more coherent RPPA signature (**Figure 7B**). A visibly larger fraction of proteins exceeded both nominal significance and effect-size thresholds, demonstrating that transcriptional activation of PIK3CA is more efficiently propagated to the proteomic layer than mutation status alone. Among the most prominent features, EEF2 showed the strongest positive shift in PIK3CA-overexpressed tumours, consistent with enhanced translational capacity downstream of growth and metabolic signalling. This pattern supports the view that transcriptional activation represents a functionally executed PI3K state at the protein level. Analysis of PIK3CA methylation state revealed an intermediate phenotype (**Figure 7C**). While most proteins again clustered near log₂ fold change ≈ 0, a limited subset crossed nominal significance thresholds with moderate effect sizes. Proteins such as IDO, IGFBP2, p16^INK4A, and cleaved Caspase-7 were among the most prominent features, implicating immune modulation, growth-factor signalling, cell-cycle control, and apoptotic execution. Thus, epigenetic stratification generates biologically structured but comparatively diffuse proteomic changes—stronger than those driven by mutation status yet weaker than those associated with transcriptional activation. To determine whether the limited proteomic footprint of PIK3CA mutation reflects upstream regulatory adaptation, we examined CpG-level methylation changes within PI3K pathway regulators. As shown in **Figure 7D**, the strongest and most significant differentially methylated probes were concentrated in PIK3R1 and PIK3CD. Leading CpGs (e.g., PIK3R1: cg25008444, cg06445944, cg01239651, cg09101894; PIK3CD: cg16512854) exhibited negative Δβ values, consistent with relative hypomethylation in PIK3CA-mutant tumours. This directional bias indicates targeted epigenetic remodelling of key PI3K regulatory elements accompanying oncogenic PIK3CA mutation. Together, **Figure 7A–D** establish a stratified hierarchy of PI3K state encoding across molecular layers. PIK3CA mutation produces selective and sparse proteomic changes but is accompanied by focused epigenetic adaptation of PI3K regulators; PIK3CA methylation state generates moderate, biologically structured proteomic shifts; and PIK3CA transcriptional activation yields the strongest and most coherent RPPA signature. These results demonstrate that transcriptional state is the dominant executor of PI3K activity at the proteomic level, while genomic and epigenomic states contribute complementary and mechanistically informative layers of regulation. This multi-omics dissociation provides a strong rationale for modelling PI3K pathway activity using orthogonal molecular states rather than mutation status alone.

**Figure 7.**
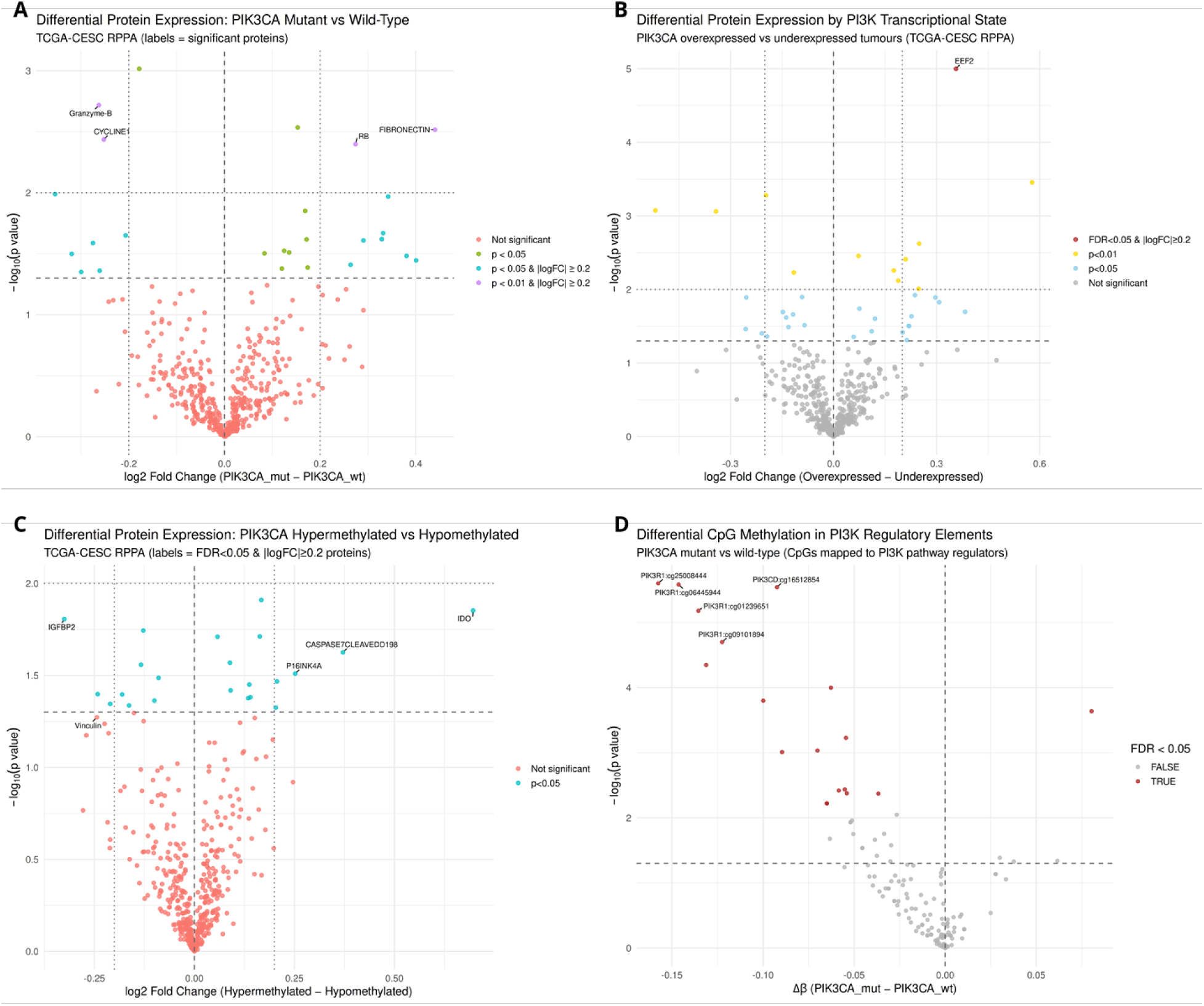
Proteomic consequences of orthogonal PI3K states and CpG-level remodelling of PI3K regulators in TCGA-CESC. **A:** RPPA differential protein abundance for PIK3CA-mutant vs wild-type tumours; labelled points denote the most significant proteins, with dashed/dotted reference lines indicating nominal p-value and effect-size thresholds. **B:** RPPA differential protein abundance for *PIK3CA* overexpressed vs underexpressed tumours; *EEF2* is highlighted among the strongest positive-shift proteins. **C:** RPPA differential protein abundance for *PIK3CA* hypermethylated vs hypomethylated tumours; a limited subset of proteins (e.g., *IDO, IGFBP2, P16INK4A, cleaved Caspase-7*) shows nominally significant shifts. **D:** CpG-level differential methylation within PI3K regulatory genes for PIK3CA-mutant vs wild-type tumours; leading differentially methylated probes map primarily to *PIK3R1* and *PIK3CD* and show negative Δβ values (relative hypomethylation in mutants), indicating targeted epigenetic adaptation of PI3K pathway regulators.

### Elastic-net RPPA classifiers for orthogonal states in cervical cancer with functional validation

#### PIK3CA multi-omic state inference reveals modality-specific proteomic encoding with strongest signal for methylation-defined states

Elastic-net classifiers trained on RPPA features demonstrated clear, modality-dependent separability of PIK3CA molecular states **(Figure 8)**. Four patients were excluded from methylation modelling due to undefined gene-level methylation (all-NA β values), yielding n = 170 for methylation and expression analyses and n = 172 for mutation status. Discrimination was complete for the PIK3CA methylation-defined state (AUC = 1.000, n = 170), whereas performance was moderate-to-high for PIK3CA mutation status (AUC = 0.880, n = 172) and expression-defined state (AUC = 0.825, n = 170), indicating that epigenetically anchored PIK3CA states are most consistently propagated to the proteome in this cohort. Reactome pathway enrichment of model-derived RPPA features further clarified the signalling architecture underlying PIK3CA-associated tumour states **(Figure 9E)**. Enriched pathways prominently included PI3K–AKT signalling, receptor tyrosine kinase activation, cell-cycle regulation, DNA damage response, and metabolic control pathways, consistent with the central role of PIK3CA in integrating growth-factor signalling with downstream metabolic and proliferative programmes. Additional enrichment of pathways linked to inflammatory signalling and transcriptional regulation further indicates that PIK3CA-associated states coordinate oncogenic signalling with broader regulatory circuits governing tumour adaptation. Feature-level interrogation revealed distinct and modality-specific proteomic programmes **(Supplementary Table S7)**. The expression-anchored model was characterised by positive coefficients for SRC and INPP4B, alongside coordinated negative loadings for translational and metabolic regulators (EEF2, HK2), apoptosis-related proteins (BAK, MCL1), and stress-response mediators (ATR, PPIF / Cyclophilin-F), consistent with altered survival signalling, translational control, and metabolic flux accompanying transcriptionally defined PIK3CA states. In contrast, the methylation-anchored model was dominated by large-magnitude coefficients implicating tumour suppressor and transcriptional regulators (VHL, RB1, NF2), oncogenic and inflammatory signalling nodes (BRAF, STAT3), DNA damage and chromatin-associated machinery (DDB1, SETD2), and metabolic and translational control components (ACSS1, SLC16A3 / MCT4, EIF4EBP1). Together, these features form a coherent multi-pathway signature spanning growth-factor signalling, chromatin regulation, metabolic rewiring, and replication stress tolerance, plausibly underlying the near-perfect classifier performance observed for methylation-defined PIK3CA states **(Figure 8)**. Finally, the mutation-anchored model highlighted a more heterogeneous but still discriminable proteomic imprint, with contributions from extracellular matrix and proliferation-associated proteins (Fibronectin, CCNE1, CCND3), stress and signalling intermediates (MAPK14 / p38, AKT2), autophagy regulation (ULK1), and DNA repair machinery (LIG4). These results indicate that mutation-defined PIK3CA states are associated with broader rewiring of stress signalling, proliferative control, and extracellular matrix interaction within the tumour proteome. Consistent with these signalling patterns, transcription factor regulatory network analysis revealed that the PIK3CA-associated regulatory programme is dominated by ESR1-centred transcriptional control coupled to RB1–E2F cell-cycle regulatory interactions, linking hormone-responsive transcriptional circuitry with proliferative signalling networks **(Figure 10)**. Collectively, these results demonstrate that PIK3CA status is encoded by distinct RPPA feature sets across molecular layers, with the strongest and most coherent proteomic representation arising from epigenetically defined PIK3CA states **(Supplementary Table S7)**. This hierarchy suggests that epigenetic regulation may stabilise oncogenic PI3K signalling outputs, enabling consistent downstream propagation to the tumour proteome. Consistent with these pathway and regulatory patterns, protein–protein interaction network analysis further revealed interconnected signalling modules linking replication stress tolerance, inflammatory signalling, developmental pathways, and metabolic adaptation within the PIK3CA-associated tumour state **(Figure 12)**. The STRING-derived network highlighted interactions among proteins including ATR, LIG4, STAT3, AKT2, BRAF, HK2, ACSS1, and SLC16A3, integrating DNA repair machinery, oncogenic signalling, metabolic control, and transcriptional regulation into a unified functional framework. These network relationships illustrate how multiple signalling modules converge to define the PIK3CA methylation-associated proteomic state in cervical cancer.

**Figure 8.**
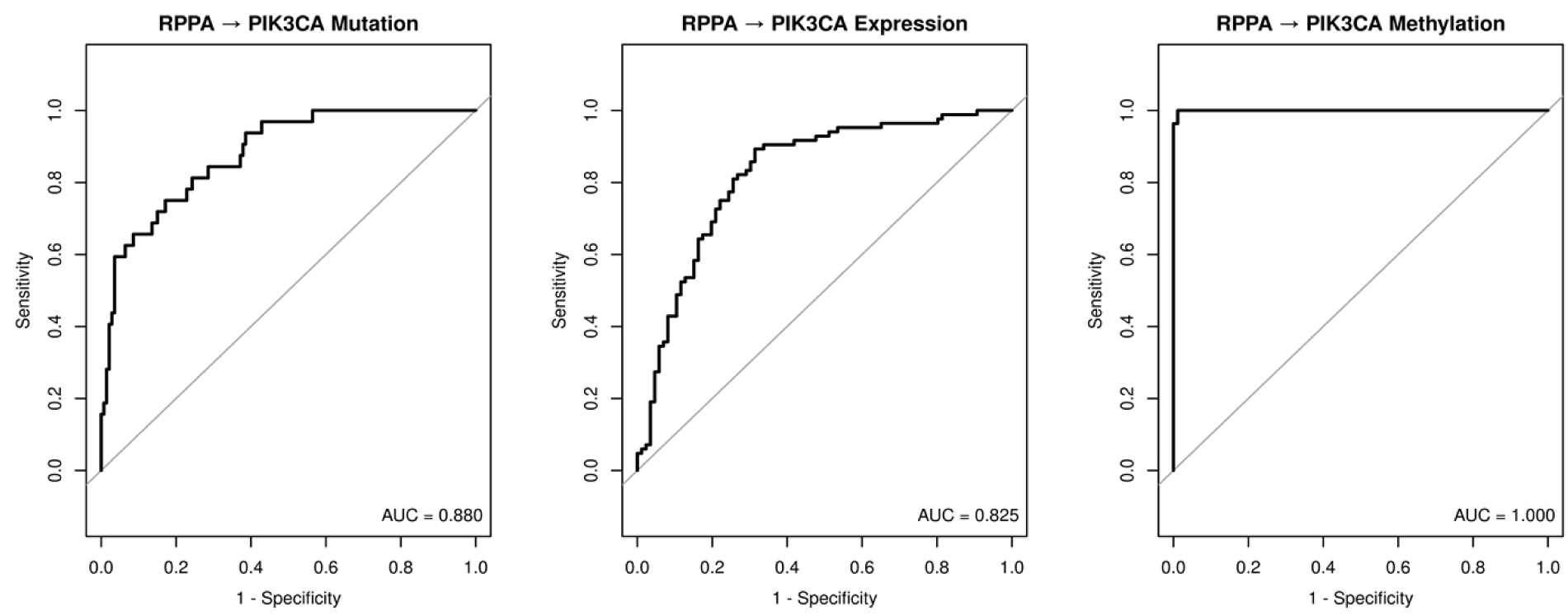
Proteome-based inference of PIK3CA molecular states across genomic, transcriptomic, and epigenomic modalities. Receiver operating characteristic (ROC) curves summarising elastic-net classifier performance for predicting *PIK3CA* molecular states using RPPA proteomic features. **Left**: *PIK3CA* mutation status (mutant vs wild type; n = 172) shows moderate-to-high discrimination (AUC = 0.880). **Middle:** *PIK3CA* expression state (high vs low; n = 170) demonstrates robust but weaker separability (AUC = 0.825). **Right:** PIK3CA methylation state (hypermethylated vs hypomethylated; n = 170) is predicted with near-perfect accuracy (AUC = 1.000). Grey diagonals indicate no-discrimination baselines. Differences in classifier performance highlight modality-specific proteomic encoding of *PIK3CA* status, with epigenetically defined states exhibiting the most coherent and discriminative proteomic signature in the TCGA-CESC cohort. **\*** Equivalent RPPA-based classifier performance plots for other recurrently mutated genes analysed in this study are provided in the Supplementary Figure S6.

**Figure 9.**
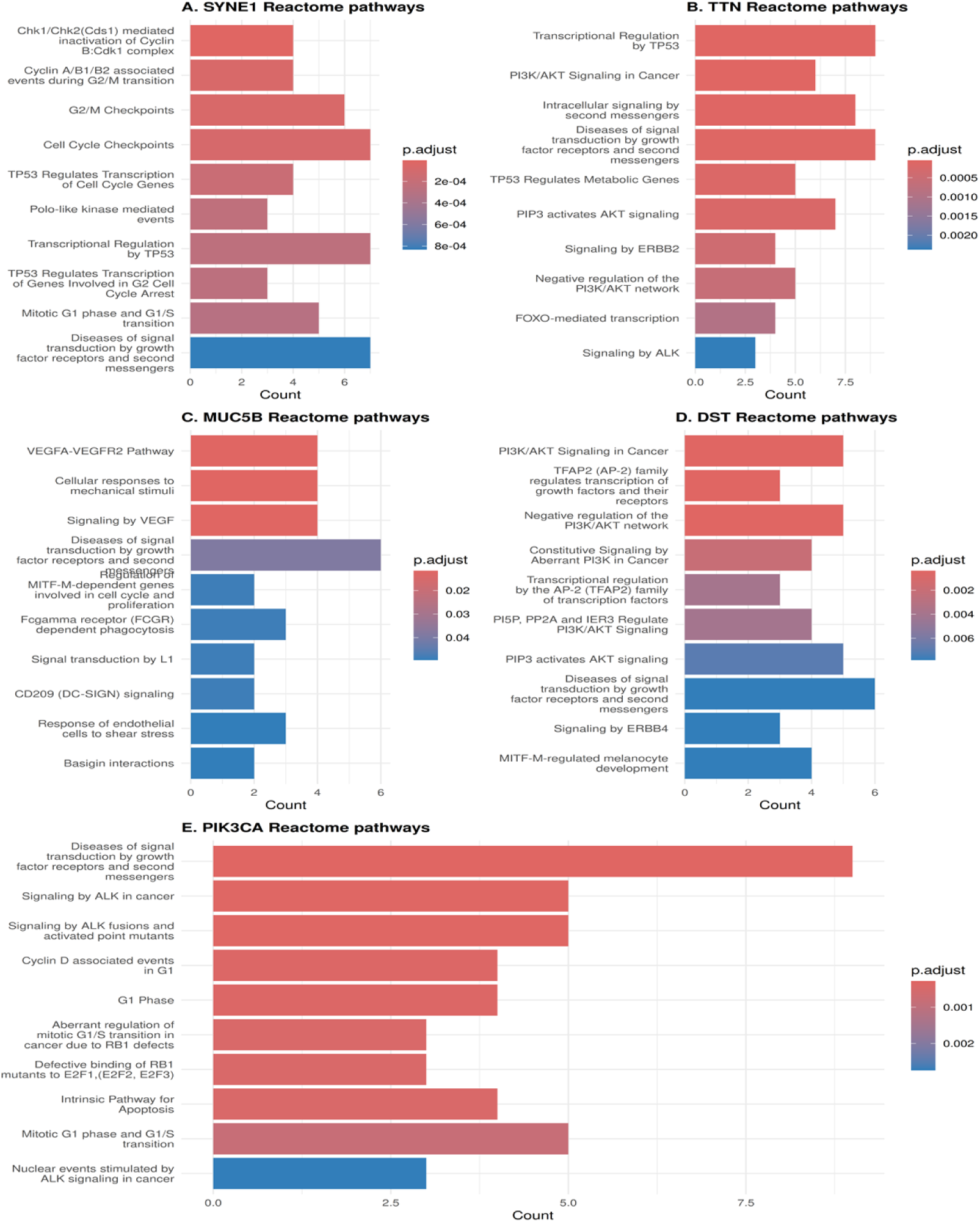
*Somatic mutation–associated Reactome pathway enrichment across proteome-defined tumour states in TCGA-CESC*. Reactome enrichment was performed on RPPA features underlying elastic-net tumour-state classifiers. Bar length indicates feature count and colour represents adjusted p-value. (A) SYNE1: Enrichment of cell-cycle checkpoints and TP53-regulated transcription. (B) TTN: Signalling-rewired state dominated by TP53, PI3K–AKT, and receptor tyrosine kinase pathways (ERBB2, ALK), with concurrent positive and negative regulation of AKT signalling, indicating dynamic signalling flux and stress adaptation. (C) MUC5B: VEGF signalling, mechanotransduction, and immune-related pathways, consistent with tumour–microenvironment interaction. (D) DST: PI3K–AKT regulation, AP-2 transcriptional control, and ERBB signalling. (E) PIK3CA: Canonical growth factor signalling, AKT activation, cell-cycle progression, and apoptosis pathways. Overall, mutation-associated states resolve into distinct proteomic programmes spanning checkpoint control, signalling rewiring, and microenvironmental interaction.

**Figure 10.**
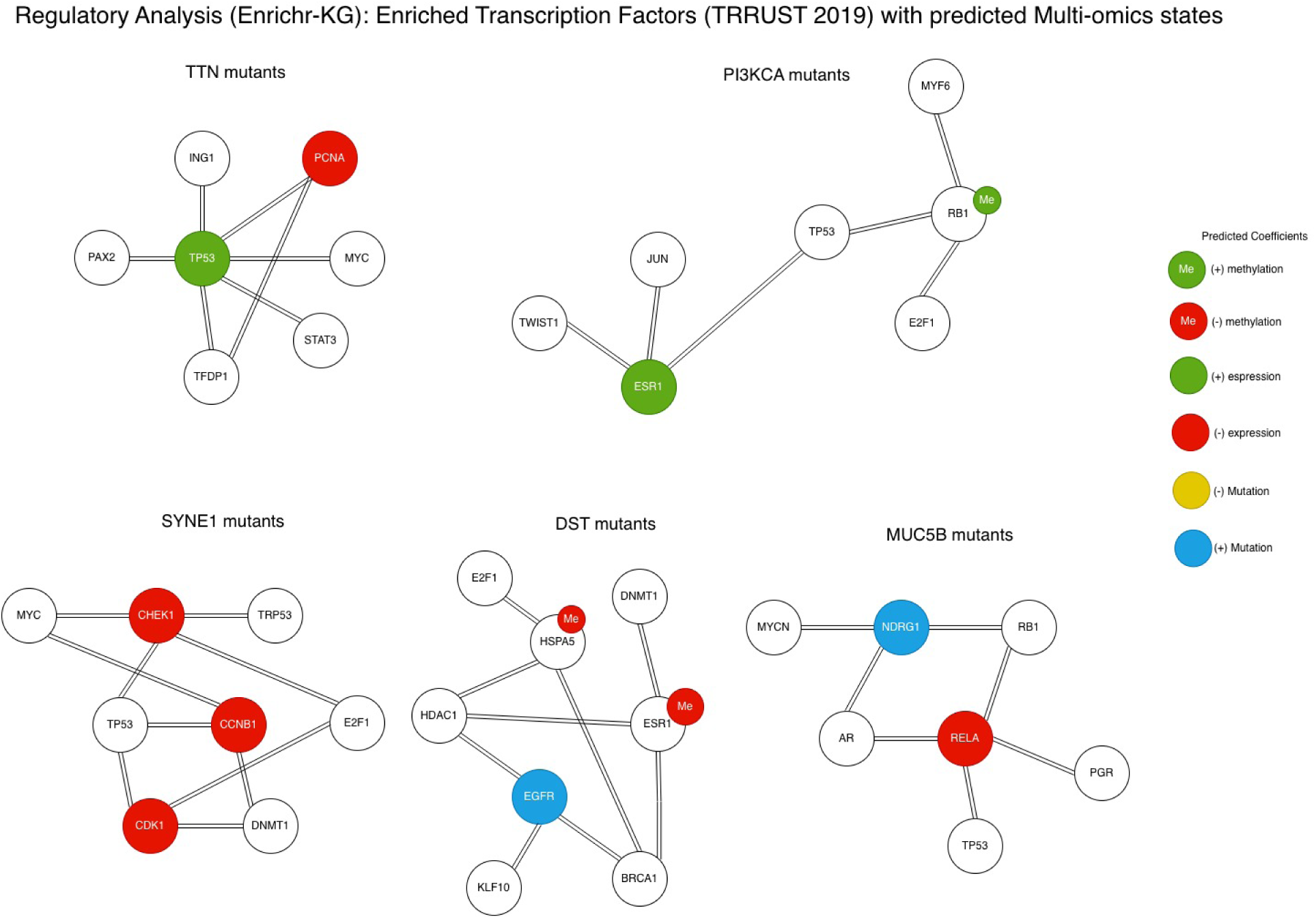
Multi-omics regulatory analysis of mutation-associated transcription factor networks in cervical cancer. Transcription factor enrichment networks derived from Enrichr-KG using the top multi-omics features associated with recurrently mutated genes in the cohort. Networks are shown for TTN, PIK3CA, SYNE1, DST, and MUC5B mutant programmes, illustrating the transcriptional regulators predicted to control each mutation-defined state. The TTN-associated module is centred on TP53 and reflects replication stress signalling, while the PIK3CA programme is dominated by ESR1 with RB1–E2F cell-cycle regulatory interactions. The SYNE1 programme highlights checkpoint and mitotic regulators including CHEK1, CDK1, and CCNB1, consistent with replication stress and checkpoint fragility. In contrast, the DST programme integrates receptor signalling and chromatin regulation, including EGFR gain-of-function and epigenetic modulation of ESR1 and HSPA5. The MUC5B module is dominated by RELA-mediated inflammatory transcriptional activity with NDRG1 gain-of-function, consistent with stress-responsive signalling and tumour–microenvironment interactions. Node colours represent multi-omics annotations: green, positive expression; red, negative expression; green “Me”, positive methylation; red “Me”, negative methylation; yellow, negative mutation; and blue, positive mutation. “+” indicates positive coefficients and “−” indicates negative coefficients.

#### SYNE1 multi-omic state inference identifies a highly separable mutation/expression proteomic programme with weaker methylation encoding

Elastic-net classifiers trained on RPPA features demonstrated strong proteome-level separability of SYNE1 mutation and expression states, with near-perfect discrimination for mutation status (AUC = 1.000, n = 172) and expression-defined state (AUC = 0.999, n = 170) **(Supplementary Figure S6)**. In contrast, performance for the SYNE1 methylation-defined state was substantially lower (AUC = 0.736, n = 172), indicating that epigenetically defined SYNE1 variation is less consistently propagated to protein-level signalling in this cohort. Reactome pathway enrichment of model-derived RPPA features further clarified the signalling architecture underlying these SYNE1-associated states **(Figure 9A)**. Enriched pathways highlighted cell-cycle regulation, checkpoint signalling, DNA damage response pathways, and receptor-mediated signalling modules, consistent with coordinated activation of replication stress tolerance and adaptive survival signalling. Additional enrichment of pathways related to cytoskeletal organisation and nuclear structural regulation further aligns with the established role of SYNE1 in nuclear envelope architecture and mechanotransduction. Together, these pathway programmes suggest that SYNE1-associated tumour states integrate structural perturbation with checkpoint rewiring and stress-responsive signalling. Feature-level interrogation revealed a coherent survival- and checkpoint-linked proteomic programme dominated by expression- and mutation-driven effects **(Table 4)**. The expression-based model was characterised by strong negative loading of the tumour suppressor PTEN (coef = −0.650), accompanied by positive coefficients for DNA damage and replication-associated regulators (MSH2, CDC6), oncogenic signalling (BRAF), and metabolic adaptation (ACSS1), alongside increased CDKN1B (p27). Conversely, key G2/M machinery and checkpoint components—including CDK1, CCNB1 (Cyclin B1), and CHEK1—as well as the structural cytoskeletal protein EPPK1, carried negative coefficients. Together, this pattern indicates altered checkpoint enforcement and replication stress adaptation rather than wholesale proliferative activation, consistent with SYNE1’s role in nuclear architecture and mechanotransduction. The mutation-defined model exhibited large-magnitude coefficients implicating iron metabolism and proliferative demand (TFRC, coef = 0.898), inflammatory and survival signalling (RELA / NF-κB p65), apoptosis regulation (BCL2L11 / BIM), and invasion or adhesion (L1CAM / CD171). These were accompanied by strong negative coefficients for kinase and cell-cycle regulators (LYN, CDC25C), mitochondrial and translational machinery (TUFM), ubiquitin-associated regulation (UBAC1), and developmental pathway signalling (PTCH1), suggesting extensive rewiring of kinase signalling, checkpoint control, and cellular stress-response pathways in SYNE1-mutant tumours. In contrast, the methylation-defined model contributed only small-magnitude effects across a limited set of proteins (CDKN1B/p27, cleaved CASP7, ACSS1, PDGFRB, CD274/PD-L1, IDO1), consistent with its reduced discriminatory performance and indicating comparatively weak transmission of SYNE1 epigenetic variation to the proteomic layer. Consistent with these findings, transcription factor regulatory network analysis highlighted checkpoint and mitotic regulators including CHEK1, CDK1, and CCNB1 as central nodes within the SYNE1-associated regulatory programme, supporting a model of replication stress signalling and checkpoint fragility **(Figure 10)**. Collectively, these results define a SYNE1-associated proteomic programme characterised by PTEN-linked survival signalling, replication stress tolerance, and attenuated G2/M checkpoint control, with particularly strong and coherent proteomic encoding of mutation and expression states but limited signal for methylation-defined SYNE1 status. This hierarchy mirrors the broader theme of the study, in which structural and mechanotransductive gene alterations preferentially manifest at the functional proteomic level through stress-adaptive and signalling-rewired tumour states rather than through uniform epigenetic propagation. Consistent with this model, protein–protein interaction network analysis further revealed interconnected signalling modules linking growth-factor signalling, checkpoint control, and metabolic adaptation within the SYNE1-associated tumour state **(Figure 15)**. The STRING-derived network highlighted key interaction hubs including PDGFRB, BRAF, PTEN, CDK1, and CCNB1, integrating receptor tyrosine kinase signalling with cell-cycle checkpoint regulation and survival signalling pathways. These network relationships provide a mechanistic framework for understanding how structural alterations in SYNE1 propagate to downstream signalling modules within the tumour proteome.

**Table 4.** SYNE1-associated multi-omic feature coefficients across expression, methylation, and mutation models.

| # | feature | coef | abscoef | gene_symbol | model |
| --- | --- | --- | --- | --- | --- |
| 1 | AGID00061 | -0.650 | 0.650 | PTEN | Expression |
| 2 | AGID00390 | 0.601 | 0.601 | MSH2 | Expression |
| 3 | AGID00014 | 0.525 | 0.525 | BRAF | Expression |
| 4 | AGID00105 | 0.503 | 0.503 | P27 | Expression |
| 5 | AGID00408 | 0.418 | 0.418 | AceCS1 | Expression |
| 6 | AGID00075 | -0.368 | 0.368 | EPPK1 | Expression |
| 7 | AGID00293 | -0.366 | 0.366 | CDK1 | Expression |
| 8 | AGID00024 | -0.365 | 0.365 | CYCLINB1 | Expression |
| 9 | AGID00422 | 0.349 | 0.349 | Cdc6 | Expression |
| 10 | AGID00189 | -0.344 | 0.344 | CHK1 | Expression |
| 11 | AGID00105 | -0.121 | 0.121 | P27 | Methylation |
| 12 | AGID00015 | -0.0813 | 0.0813 | CASPASE7CLEAVEDD198 | Methylation |
| 13 | AGID00408 | 0.0484 | 0.0484 | AceCS1 | Methylation |
| 14 | AGID00385 | 0.0234 | 0.0234 | PDGFRB | Methylation |
| 15 | AGID00300 | -0.0102 | 0.0102 | PDL1 | Methylation |
| 16 | AGID00366 | -0.00279 | 0.00279 | IDO | Methylation |
| 17 | AGID00165 | 0.898 | 0.898 | TFRC | Mutation |
| 18 | AGID00439 | -0.825 | 0.825 | Lyn | Mutation |
| 19 | AGID00318 | 0.821 | 0.821 | Myt1 | Mutation |
| 20 | AGID00329 | -0.795 | 0.795 | cdc25C | Mutation |
| 21 | AGID00303 | 0.789 | 0.789 | CD171 | Mutation |
| 22 | AGID00214 | -0.738 | 0.738 | UBAC1 | Mutation |
| 23 | AGID00048 | 0.721 | 0.721 | NFKBP65 | Mutation |
| 24 | AGID00180 | 0.668 | 0.668 | BIM | Mutation |
| 25 | AGID00417 | -0.661 | 0.661 | TUFM | Mutation |
| 26 | AGID00274 | -0.659 | 0.659 | Patched | Mutation |

\*Comparable elastic-net coefficient tables summarising expression-, methylation-, and mutation-associated proteomic features for other recurrent mutation programmes analysed in this study (PIK3CA, TTN, MUC5B, and DST) are provided in the Supplementary Materials.

#### TTN multi-omic state inference reveals a stress-adapted, signalling-rewired proteomic programme

Elastic-net classifiers trained on RPPA features demonstrated robust proteome-level encoding of TTN molecular states across mutation, expression, and methylation layers **(Supplementary Figure S6)**. Discriminatory performance was near-perfect for TTN mutation status (AUC = 1.000, n = 172) and remained high for expression-defined (AUC = 0.952, n = 170) and methylation-defined states (AUC = 0.917, n = 170), indicating that TTN-associated variation is consistently captured at the proteomic level across orthogonal regulatory modalities despite its likely passenger status. Reactome pathway enrichment of model-derived RPPA features further clarified the signalling architecture underlying these TTN-associated states **(Figure 9B)**. Dominant pathways included TP53-dependent transcriptional regulation, PI3K–AKT signalling, and receptor-mediated kinase cascades (ERBB2, ALK), alongside FOXO-mediated transcription and PIP3-dependent control of AKT activation. The simultaneous enrichment of both activating and inhibitory PI3K/AKT modules indicates a proteomic landscape characterised by dynamic signalling flux, rather than simple pathway hyperactivation. Consistent with this architecture, the expression-derived classifier was driven by positive coefficients for signalling and translational regulators, including ERBB2, SRC, PAK1, EIF4E, and TP53, alongside negative contributions from checkpoint and replication-associated proteins such as CHEK2, AKT2, PCNA, and PAICS. This configuration supports a state of enhanced kinase signalling and translational output coupled with attenuated checkpoint enforcement. The methylation-defined model highlighted coordinated epigenetic modulation of lineage specification, redox buffering, and metabolic flexibility. Strong negative loading of CDKN1B (p27) was accompanied by positive coefficients for GATA3, PRDX1, G6PD, ACSS1, and TIGAR, indicating suppression of cell-cycle restraint alongside reinforcement of oxidative stress buffering and metabolic adaptation. In contrast, the mutation-defined model was dominated by large-magnitude coefficients implicating mitochondrial and metabolic regulators (TFAM, ACSS1), developmental signalling nodes (SMAD4, LKB1), and immune-associated kinases (ZAP70, IRS1), with concurrent negative contributions from adaptor and cytoskeletal regulators (FRS2-α, SYK, PEA15, EPPK1) **(Supplementary Table S8)**. Collectively, these multi-omic patterns converge on a proteomic state characterised by altered immune–metabolic signalling, translational activation, and cytoskeletal remodelling, rather than pathway-centric oncogene addiction. This supports a model in which TTN mutation status marks a stress-adapted, signalling-rewired tumour state, reflecting systems-level adaptation rather than direct oncogenic drive.

#### MUC5B multi-omic state inference defines a replication-stressed, transcriptionally active proteomic phenotype

Elastic-net classifiers trained on RPPA features demonstrated strong proteome-level encoding of MUC5B molecular states across mutation, expression, and methylation layers **(Supplementary Figure S6)**. Classification performance was complete for both MUC5B mutation status (AUC = 1.000, n = 172) and MUC5B expression-defined state (AUC = 1.000, n = 170), indicating highly concordant and robust proteomic signatures associated with these modalities. In contrast, predictive performance for the methylation-defined MUC5B state was more modest (AUC = 0.752, n = 172), suggesting that epigenetic variation at the MUC5B locus is only partially transmitted to downstream protein signalling. Reactome pathway enrichment of model-derived RPPA features further contextualised these proteomic programmes **(Figure 9C)**. The dominant enriched pathways highlighted VEGF–VEGFR2 signalling, receptor-mediated growth-factor signalling, and cellular responses to mechanical stimuli, alongside immune and adhesion-related pathways including Fcγ receptor–dependent phagocytosis, CD209 (DC-SIGN) signalling, and Basigin interactions. Additional enrichment of endothelial shear stress responses and VEGF signalling pathways indicates coordinated engagement of angiogenic and microenvironmental sensing programmes. Together, these pathways suggest that MUC5B-associated proteomic states integrate replication stress signalling with adaptive responses to hypoxia, vascular remodelling, and tumour–microenvironment interactions. Feature-level inspection of model coefficients revealed a biologically coherent stress-adapted programme dominated by replication and transcriptional stress responses **(Supplementary Table S9)**. The expression-based model was characterised by strong positive loading of ATR, alongside increased contribution from XBP1, consistent with activation of DNA damage sensing, replication stress tolerance, and unfolded protein response pathways. In parallel, negative coefficients were observed for multiple signalling, metabolic, and structural nodes, including AKT1S1 (PRAS40), IRS2, PTK2 (FAK), RELA (NF-κB p65), GLI1, PAK1, SLC16A3 (MCT4), and MFN1, indicating coordinated attenuation of growth-factor signalling, cytoskeletal dynamics, hypoxia-linked glycolytic flux, and mitochondrial fusion. Collectively, this pattern suggests a proteomic shift away from canonical oncogenic signalling toward survival under sustained transcriptional and proteotoxic stress. The mutation-defined model highlighted a distinct but complementary proteomic programme. Positive coefficients for ENO2, CHD1L, NDRG1, ACSS1, L1CAM, ESR1, ERCC4 (XPF), and CCNB1 implicate metabolic rewiring, chromatin remodelling, DNA repair capacity, hormone-linked signalling, and cell-cycle progression as core features of the MUC5B-mutant state. Concurrent negative loading of IGFBP2 and HK1 further supports altered growth-factor responsiveness and reprogrammed glucose utilisation in these tumours. In contrast, the methylation-defined model contributed only low-magnitude coefficients across a limited set of metabolic and junctional proteins (HK1, CTNNB1, GJA1, CD38, PLCG1), consistent with its reduced discriminatory power and weaker proteomic imprint. Taken together, the high predictive accuracy for mutation and expression states **(Supplementary Figure S6)** and the structured, modality-specific coefficient patterns **(Supplementary Table S9)**, together with pathway enrichment results **(Figure 8C)**, define a MUC5B-associated proteomic phenotype characterised by replication stress tolerance, enhanced transcriptional and unfolded protein response activity, metabolic adaptation, angiogenic signalling, and selective dampening of canonical growth-factor and cytoskeletal signalling pathways. This coherent stress-adapted tumour state provides a strong biological foundation for the replication- and transcription-targeted therapeutic vulnerabilities prioritised through downstream connectivity mapping. Transcription factor regulatory networks, together with pathway-level insights for mutation-defined multi-omic states (TTN, PIK3CA, SYNE1, DST, and MUC5B), were obtained via Enricher-KG (TRRUST 2019 library) and are illustrated in **Figure 10**, providing a consistent, network-based framework for systematically linking genes to upstream transcriptional regulators and functional pathways across all downstream analyses.

#### DST multi-omic state inference reveals a cytoskeleton-linked, growth factor– and immune-modulated proteomic programme

Elastic-net classifiers trained on RPPA proteomic features demonstrated strong but modality-dependent separability of DST-associated molecular states **(Supplementary Figure S6)**. Proteomic features predicted DST expression-defined states with near-perfect accuracy (AUC = 0.983, n = 170) and achieved complete discrimination for methylation-defined DST states (AUC = 1.000, n = 172), indicating robust propagation of transcriptional and epigenetic variation to downstream protein signalling. In contrast, classification of DST mutation status showed reduced but still robust performance (AUC = 0.820, n = 172), consistent with heterogeneous proteomic consequences of DST genomic alteration relative to its regulatory states. Reactome pathway enrichment of RPPA features associated with DST multi-omic states further clarified the signalling architecture underlying this programme **(Figure 9D)**. Enriched pathways highlighted cytoskeleton organisation, receptor tyrosine kinase signalling, and integrin-mediated adhesion pathways, alongside immune-modulatory and stress-response signalling modules. Prominent enrichment of EGFR- and KIT-associated receptor signalling, together with pathways related to cytoskeletal remodelling and extracellular matrix interaction, suggests coordinated coupling of structural epithelial integrity with growth-factor–driven signalling and microenvironmental sensing. Additional pathways involving immune signalling and metabolic regulation further indicate that DST-associated states integrate structural perturbation with adaptive signalling responses to tumour–microenvironment interactions. Feature-level analysis revealed that the expression model captured a coherent programme linking cell-cycle progression, DNA repair, cytoskeletal organisation, and receptor tyrosine kinase signalling **(Supplementary Table S10)**. Strong positive coefficients were observed for CCNE1 (Cyclin E1) and TRIP13, implicating G1/S progression and replication stress tolerance, alongside KIT, suggesting growth-factor–driven signalling activation. Negative loadings for MSH2, IRF3, TMEM173 (STING), and SFRP1 indicate attenuation of DNA mismatch repair, innate immune signalling, and WNT antagonism, consistent with immune-evasive and proliferative adaptation. Additional negative coefficients for TUBA/TUBD (α/δ-tubulin) and LASU1 further support perturbation of cytoskeletal integrity and ribosomal-associated processes, aligning with DST’s established structural role in epithelial cells. The methylation model exhibited the strongest and most biologically expansive signature, dominated by large-magnitude coefficients implicating angiogenesis, mechanotransduction, metabolic rewiring, and oncogenic signalling. Positive coefficients for IGFBP2, PECAM1 (CD31), TGM2 (transglutaminase), ASNS, AKT2, RPS6KA1 (p90RSK), and BRAF point to coordinated activation of growth-factor signalling, endothelial interaction, amino acid metabolism, and MAPK–AKT axis engagement. Concurrent negative loadings for ESR1, YAP1, and HSPA5 (BiP/GRP78) suggest selective suppression of hormone signalling, Hippo pathway output, and endoplasmic reticulum stress buffering. This profile indicates that epigenetic DST states are tightly coupled to vascular, metabolic, and mechanosensitive tumour adaptations. In contrast, the mutation model was characterised by smaller-magnitude coefficients, reflecting a more focal proteomic footprint. Positive contributions from CD274 (PD-L1), EGFR, SERPINE1 (PAI1), IRS1, and FASN suggest engagement of immune checkpoint signalling, growth-factor responsiveness, extracellular matrix remodelling, and lipid metabolism in DST-mutant tumours, while negative coefficients for FTO and IGFBP2 indicate selective modulation of RNA metabolism and growth-factor binding. The comparatively attenuated effect sizes are consistent with mutation status acting as a less deterministic driver of the DST-associated proteomic state relative to transcriptional or epigenetic regulation. Consistent with these pathway and proteomic patterns, transcription factor regulatory network analysis revealed that the DST programme integrates receptor signalling and chromatin regulation, including EGFR gain-of-function mutation and epigenetic modulation of ESR1 and HSPA5, linking receptor tyrosine kinase signalling with transcriptional and stress-response regulatory circuits **(Figure 10)**. Integration of these multi-omics features into a protein–protein interaction network further illustrates the modular signalling architecture of DST-associated states **(Figure 11).** Collectively, these results define a DST-associated proteomic programme characterised by disrupted cytoskeletal integrity coupled to compensatory activation of cell-cycle progression, receptor tyrosine kinase signalling, angiogenic interaction, metabolic adaptation, and immune modulation. The marked superiority of expression-and methylation-based classifiers over mutation-based models **(Supplementary Figure S6)** highlights regulatory rewiring as the dominant mechanism through which DST-associated states are functionally executed at the proteomic level. This integrated profile provides a strong mechanistic basis for prioritising DST-stratified therapeutic hypotheses targeting EGFR/RTK signalling, metabolic dependencies, and immune checkpoint engagement in cervical cancer.

**Figure 11.**
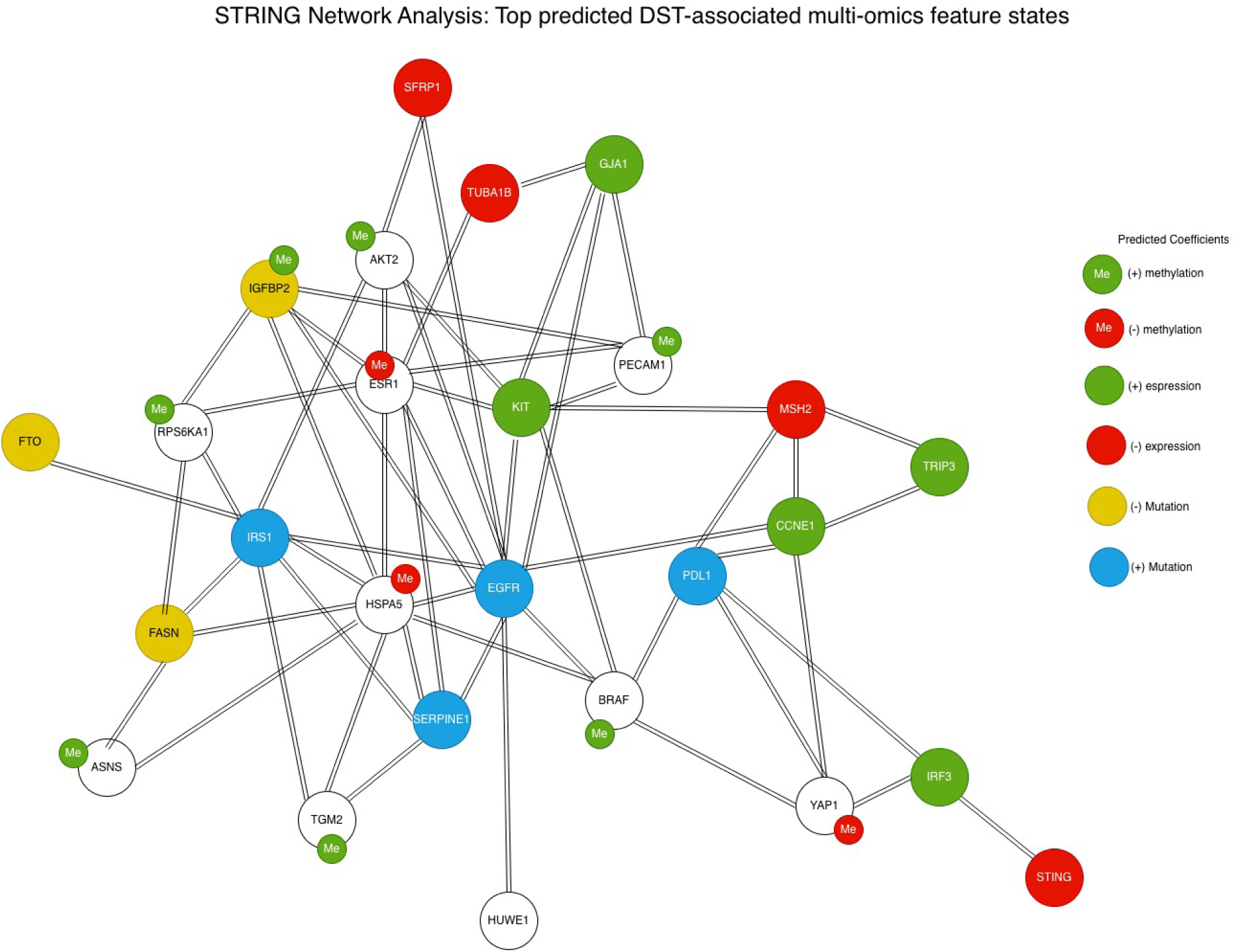
STRING protein–protein interaction network of DST-associated multi-omics features in cervical cancer. Protein–protein interaction network constructed using STRING for the top DST-associated multi-omics features identified by elastic-net modelling. Nodes represent proteins and edges indicate known or predicted protein–protein interactions. The network highlights key signalling hubs including EGFR, KIT, ESR1, CCNE1, and MSH2, linking receptor tyrosine kinase signalling, cell-cycle regulation, DNA repair, immune checkpoint pathways, and metabolic processes. The integration of transcriptional, epigenetic, and mutational annotations illustrates the complex regulatory environment associated with the DST tumour state and reveals multiple interconnected signalling modules within the protein interaction network. Node colours represent multi-omics annotations: green, positive expression; red, negative expression; green “Me”, positive methylation; red “Me”, negative methylation; yellow, negative mutation; and blue, positive mutation. “+” indicates positive coefficients and “−” indicates negative coefficients.

### L1000FWD Connectivity Mapping from RPPA-Derived Classifier Signatures

#### PIK3CA-associated connectivity mapping reveals vulnerabilities in replication stress, inflammatory signalling, developmental wiring, and metabolic control

To translate the PIK3CA-associated state into therapeutic hypotheses, we performed L1000FWD connectivity mapping using directionally consistent features derived from the elastic-net classifier. Although mutation and expression models were also evaluated (mutation AUC = 0.880; expression AUC = 0.825), the strongest discriminatory signal was observed for the methylation-defined state (AUC = 1.000), and the inferred programme is therefore best interpreted as a PIK3CA-linked epigenetic state. The PIK3CA methylation signature was characterised by positive weighting of tumour suppressor and chromatin/repair regulators (VHL, RB, SETD2) and inflammatory/STAT signalling (STAT3), together with suppression of stress-, translational-, and metabolic nodes (EIF4EBP1, MCT4) and genome maintenance factors (DDB1, NF2). In the mutational component, features such as DNA ligase IV and cell-cycle regulators further supported a replication- and repair-primed phenotype. Connectivity mapping identified multiple perturbagens with strong negative connectivity scores, predicted to reverse this inferred state, which converged on four mechanistically coherent axes: **(i)** replication stress and DNA damage vulnerability, **(ii)** inflammatory signalling suppression, **(iii)** developmental pathway dependence, and **(iv)** metabolic and oxidative stress induction. Because all identified compounds shared identical FDR-adjusted q-values (Q = 0.296), these associations should be interpreted as hypothesis-generating rather than statistically definitive. The top-ranked reversing compounds for the PIK3CA methylation–associated proteomic signature are summarised in **Table 5**. A leading reversing hit was SN-38 (BRD-A36630025), the active metabolite of irinotecan and a potent topoisomerase I inhibitor. Importantly, cervical cancer–specific in vitro evidence shows that SN-38 suppresses proliferation of HeLa and SiHa cells in a time- and dose-dependent manner and induces apoptosis through an Akt–p53/p21 axis, with downregulation of phosphorylated Akt and induction of p53 and p21; forced Akt activation partially rescues SN-38–induced apoptosis, supporting a causal role for Akt suppression in cervical cancer models (Liu et al., 2009). Targeted delivery strategies, including EGF-functionalised PLGA nanoparticles co-loaded with SN-38, further reinforce cervical cancer relevance by enhancing uptake and apoptosis induction in HeLa and CaSki cells (Zheng et al., 2023). Within our framework, these findings are mechanistically consistent with a PIK3CA-linked epigenetic state that appears replication- and repair-engaged, supporting sensitisation to agents that exacerbate replication-associated DNA damage while collapsing PI3K–Akt survival signalling. Valdecoxib (BRD-K12994359), a selective COX-2 inhibitor, also emerged as a strongly antagonistic perturbagen. While cardiovascular safety considerations limit clinical enthusiasm for coxibs, they remain of mechanistic interest in oncology. Beyond canonical COX-2 inhibition, valdecoxib has been shown to alter membrane lipid order and fluidity in cancer cells independently of COX-2 expression, suggesting a biophysical mechanism capable of modulating membrane-linked inflammatory and survival signalling aligned with STAT3/NF-κB activity (İnan Genç et al., 2017; El-Malah et al., 2022). Erismodegib (NVP-LDE225; BRD-K19796430), a Smoothened antagonist targeting Hedgehog signalling, was also identified among the reversing compounds. Hedgehog pathway inhibition has demonstrated synergy with PI3K/mTOR and receptor tyrosine kinase inhibitors in solid tumour models, supporting developmental and stemness wiring as a tractable vulnerability in PIK3CA-associated states (Pan et al., 2010; D’Amato et al., 2014).

**Table 5.**
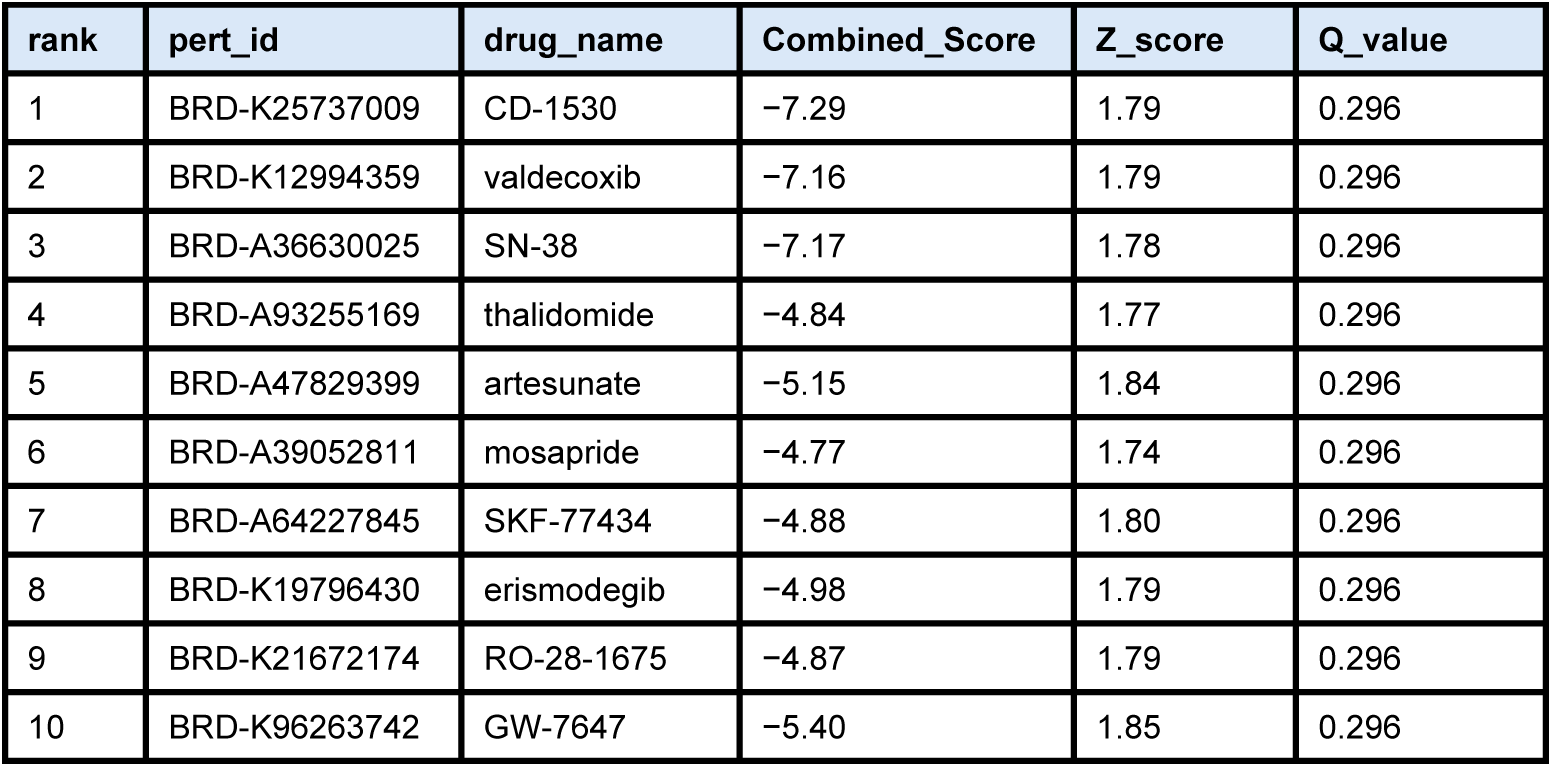
Top candidate compounds predicted to reverse the PIK3CA methylation–associated proteomic signature.

Artesunate (BRD-A47829399) showed robust antagonistic connectivity with direct cervical cancer relevance. In SiHa cervical cancer cells, artesunate inhibits growth and induces apoptosis in a dose- and time-dependent manner, accompanied by increased ROS and Ca²⁺, reduced mitochondrial membrane potential, and a shift in Bcl-2 family expression toward apoptosis (BIM/Bok/Bax/Bak ↑; Bcl-2/Bcl-xL/Mcl-1 ↓) (Sadana et al., 2025). In addition, translational evidence motivates HPV-linked relevance: topical/intravaginal artesunate is being investigated as an adjuvant strategy to improve HPV clearance in settings with high persistent HPV risk (including women living with HIV) (Zhang et al., 2024). Given that the PIK3CA signature includes metabolic/transport modulation (e.g., MCT4 and hexokinase directionality), these findings are consistent with metabolic fragility that can be exposed by oxidative and mitochondrial stressors. Additional mechanistically informative hits include thalidomide (BRD-A93255169), which is mechanistically tied to cereblon (CRBN) binding (Lopez-Girona et al., 2012). Importantly, cervical cancer–specific evidence indicates that thalidomide can synergise with cisplatin in HeLa and SiHa cells and suppress PI3K/AKT and JAK1/STAT3 signalling, providing a plausible link between immunomodulatory biology and the inflammatory/PI3K-aligned wiring inferred here (Liu et al., 2022). GW-7647 (BRD-K96263742), a potent PPARα agonist, points to lipid-inflammatory transcriptional control as a modulatory axis; while the strongest mechanistic literature is broader than cervical cancer, PPAR signalling is widely implicated in cancer metabolism and survival, supporting its inclusion as a pathway-level hypothesis (Di Leo et al., 2018; Asgharzadeh et al., 2024). RO-28-1675 (BRD-K2167721774), a glucokinase activator, highlights the possibility that glucose utilisation/flux control intersects with the PIK3CA state in perturbational space, given its known ability to shift glucose sensing/metabolism and alter systemic glucose handling *in vivo* (Iynedjian, 2008; Oh et al., 2014). Mosapride (BRD-A39052811), a 5-HT4 receptor agonist, appears as a reversing hit; while not a standard oncology agent, serotonergic signalling has recognised relevance to cancer biology, and 5-HT4-directed modulation has demonstrated anti-angiogenic effects in endothelial models, consistent with GPCR-linked transcriptional rewiring as a plausible reversal lever (Nishikawa et al., 2010; Thampongsa et al., 2025). SK-77434 (BRD-A64227845) (dopamine D1 receptor partial agonist) similarly nominates GPCR/second-messenger modulation as a perturbational route, supported by broader oncology-oriented reviews of dopamine receptor biology (Sobczuk et al., 2020; Wong et al., 1992). Finally, CD-1530 (BRD-K25737009), a selective RARγ agonist, supports retinoid-linked differentiation/transcriptional reprogramming as another plausible axis for state reversal (Powała et al., 2024). Overall, PIK3CA methylation-anchored connectivity mapping supports a coherent framework in which the PIK3CA-associated epigenetic state may be reversible by: replication stress/DNA damage induction with cervical cancer–validated apoptosis signalling effects (SN-38); inflammatory prostaglandin signalling suppression with additional lipid-membrane biophysical effects (valdecoxib); developmental pathway inhibition (erismodegib/SMO blockade); and oxidative/metabolic stress induction with direct cervical cancer evidence (artesunate), with additional support for immune-modulatory and lipid/glucose-transcriptional rewiring (thalidomide; GW-7647; RO-28-1675). Top PIK3CA-associated features identified by elastic-net modelling were submitted to STRING, generating networks in which nodes represent proteins and edges indicate documented or predicted interactions. Node annotations reflect multi-omics states (expression, methylation, and mutation), and predicted sites of perturbation by top L1000FWD reversing compounds were mapped onto the relevant functional branches of the network. This network highlights programme-specific signalling hubs, including replication stress/DNA repair (SN-38), inflammatory-survival (valdecoxib, thalidomide), developmental (erismodegib), and metabolic-oxidative (artesunate, GW-7647, RO-28-1675) modules, with additional transcriptional-differentiation modulation by CD-1530 **(Figure 12)**.

**Figure 12.**
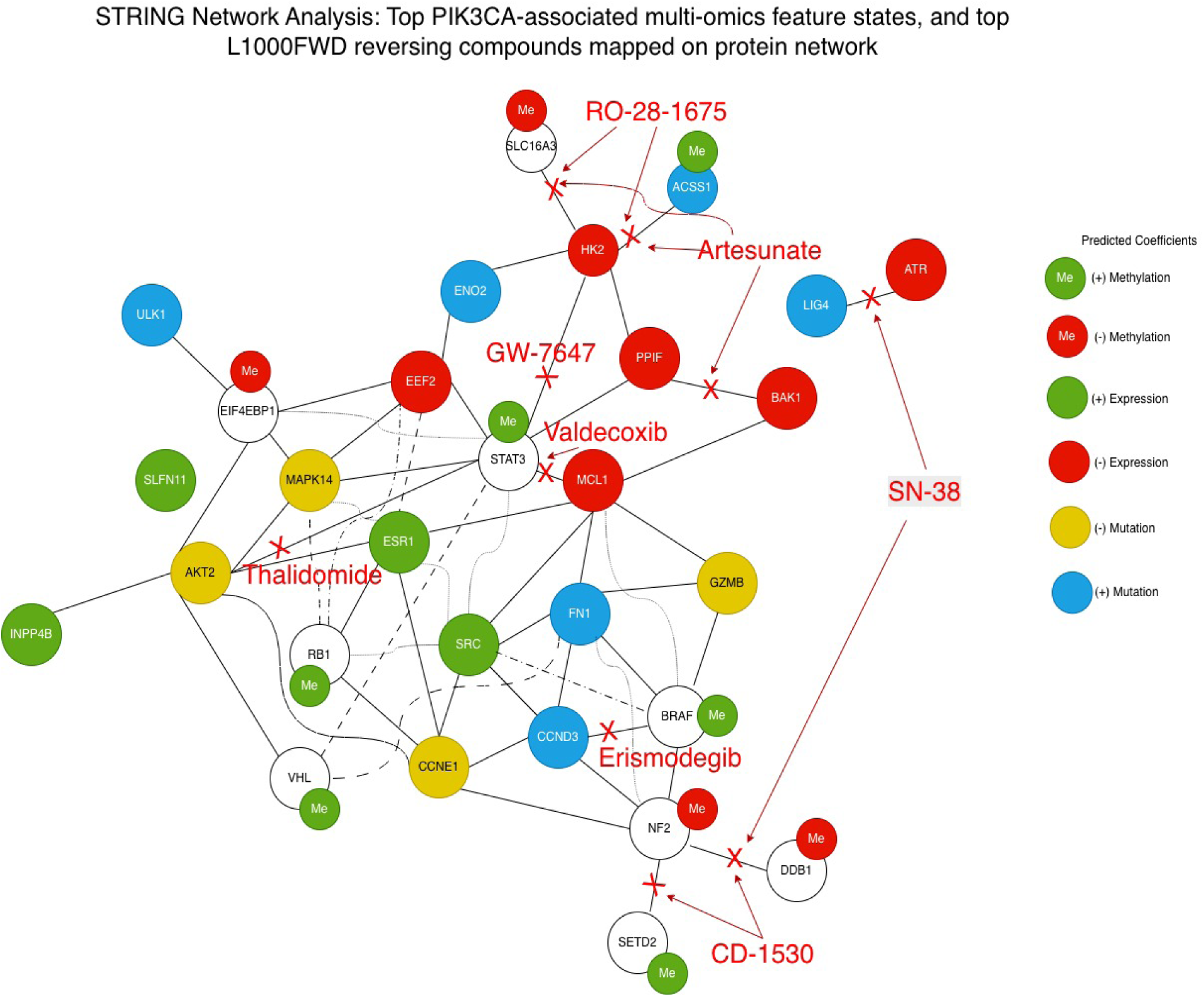
STRING protein–protein interaction network of PIK3CA-associated multi-omics features with predicted sites of action for reversing compounds. Protein–protein interaction network constructed using STRING for the top PIK3CA-associated multi-omics features, with top L1000FWD reversing compounds mapped onto the relevant functional branches of the network. Nodes represent proteins and edges indicate known or predicted protein–protein interactions. SN-38 is mapped to the replication stress/DNA repair branch (ATR/LIG4, DDB1/NF2), valdecoxib and thalidomide to the STAT3/AKT2 inflammatory-survival axis, erismodegib to the BRAF/CCND3 developmental module, and artesunate, GW-7647, and RO-28-1675 to the HK2/ACSS1/SLC16A3 metabolic-oxidative branch. CD-1530 is positioned near the NF2–SETD2/DDB1 transcriptional-differentiation region, consistent with retinoid-mediated state rewiring. Together, the network illustrates how predicted perturbagens converge on replication stress tolerance, inflammatory signalling, developmental wiring, and metabolic control within the PIK3CA methylation-associated proteomic state. Node colours represent multi-omics annotations: green, positive expression; red, negative expression; green “Me”, positive methylation; red “Me”, negative methylation; yellow, negative mutation; and blue, positive mutation. “+” indicates positive coefficients and “−” indicates negative coefficients. Red labels, arrows, and crosses indicate predicted pathway-level sites of perturbation by reversing compounds.

#### TNN-associated connectivity mapping implicates secretory-proteostasis stress, Wnt developmental wiring, immune-inflammatory control, and p53-axis fragility

To translate the TTN-associated multi-omic programme into therapeutic hypotheses, we performed L1000FWD connectivity mapping using directionally consistent proteomic features derived from integrated elastic-net models (Mutation AUC = 1.000; Expression AUC = 0.952; Methylation AUC = 0.917; **Supplementary Figure S7**). These features correspond to the modality-specific coefficient structure defining the TTN-associated proteomic state and were interpreted in the context of Reactome pathway enrichment highlighting TP53-regulated transcription, PI3K–AKT signalling cascades, receptor-mediated kinase signalling (including ERBB2 and ALK pathways), and FOXO-associated transcriptional programmes **(Figure 9B)**. Consistent with these pathway-level findings, transcription factor regulatory network analysis identified a TTN-associated regulatory module centred on TP53 and replication stress signalling **(Figure 10)**. Together, these upstream analyses define a TTN-associated tumour state characterised by signalling plasticity, checkpoint fragility, metabolic adaptation, and stress-buffering capacity, providing the biological framework for downstream connectivity mapping. The inferred TTN state was characterised by coordinated signalling and stress-response features spanning growth-factor and kinase circuitry (ERBB2, SRC, PAK1, IRS1, with AKT2 suppressed), cell-cycle and checkpoint wiring (CHEK2, PCNA, CDKN1B/p27), p53 pathway engagement (TP53 upweighted), and metabolic/redox buffering (G6PD, PRDX1, TFAM, ACSS1). This pattern is consistent with a tumour state capable of tolerating oncogenic signalling and proliferative stress through reliance on proteostasis capacity, stress checkpoints, and immunomodulatory signalling—dependencies reflected in the top reversing perturbagens identified by connectivity mapping. Connectivity analysis identified multiple perturbagens with strong negative connectivity and modest-to-borderline multiple-testing-adjusted significance (Q = 0.048–0.095), converging on four mechanistic vulnerability axes **(Table 6)**. The strongest reversing perturbagen was brefeldin A (BRD-A31107743), a fungal metabolite that disrupts ER-to-Golgi anterograde trafficking, rapidly collapsing Golgi architecture and rerouting trans-Golgi and endosomal membrane dynamics, thereby imposing acute secretory and organelle stress (Wood et al., 1991). This mechanism is particularly relevant in cervical cancer: brefeldin A derivatives have been engineered to improve selectivity and exhibit potent antiproliferative activity in HeLa cells with minimal toxicity to non-malignant cells, inducing G1 arrest and mitochondrial-dependent apoptosis (Wang et al., 2023). Within the TTN programme—characterised by stress buffering and metabolic–mitochondrial adaptation—trafficking and proteostasis disruption is consistent with the hypothesis that TTN-associated states are vulnerable to intracellular processing overload rather than single oncogenic node inhibition. XAV939 (BRD-K12762134) emerged as another prominent reversing compound with direct cervical cancer support. In HeLa models exposed to high-LET carbon ion irradiation, inhibition of Wnt/β-catenin signalling by XAV939 significantly sensitised cells, increasing apoptosis, enhancing γ-H2AX DNA damage foci, and strengthening checkpoint disruption while suppressing β-catenin pathway effectors including Wnt3a, Wnt5a, cyclin D1, and c-Myc (Wang et al., 2023). This finding aligns with the TTN programme as a signalling-adapted state in which transcriptional regulators intersect with developmental programmes, making Wnt pathway suppression a coherent strategy to destabilise TTN-associated signalling architecture. Several additional perturbagens reinforce a model of checkpoint and stress-response dependence. Serdemetan (BRD-K60219430; JNJ-26854165), originally developed as an MDM2 antagonist, has been shown to reduce clonogenic survival and act as a radiosensitiser in tumour models through induction of G2/M arrest and increased TP53–p21 signalling (Chargari et al., 2011). Although not cervical cancer–specific in the cited studies, this mechanism aligns with the TTN regulatory module centred on TP53 and replication stress **(Figure 8),** suggesting that forced engagement of stress-checkpoint signalling may destabilise TTN-associated tumour states. Similarly, radicicol (BRD-K33551950), an HSP90 inhibitor, supports vulnerability to proteostasis disruption. HSP90 inhibition destabilises numerous kinase and transcription-factor client proteins, promoting mitochondrial depolarisation and caspase activation under cytotoxic stress (Wu et al., 2013). In the TTN context, where signalling plasticity and buffering capacity appear central, collapse of chaperone-supported proteome stability provides a mechanistically consistent route for reversing the TTN-associated state. A second cluster of perturbagens points to stress–immune and redox regulatory axes. Cyclosporin A (BRD-K13533483), although classically immunosuppressive, has demonstrated cervical cancer cell-line activity in combination settings: co-treatment with arsenic trioxide induces synergistic cytotoxicity in CaSki and SiHa cells through mitochondrial membrane depolarisation, increased ROS production, and autophagy activation (Kooshan et al., 2025). This is compatible with the TTN programme’s strong metabolic and mitochondrial features and suggests that mitochondrial–ROS–autophagy stress coupling may destabilise TTN-associated tumour homeostasis. Deracoxib (BRD-K68558722), a COX-2 selective inhibitor, nominates prostaglandin-mediated inflammatory signalling as another modulatory lever. Although the cited evidence is primarily veterinary, the result reinforces the broader concept that inflammatory signalling frequently intersects with HPV-driven oncogenic circuitry in cervical cancer and may therefore provide an indirect route to reversing TTN-associated transcriptional states. JAK3 inhibitor VI (BRD-K04546108) further implicates cytokine signalling control, as JAK–STAT pathways are repeatedly implicated in cervical cancer progression and HPV E6/E7–driven signalling networks (Gutiérrez-Hoya & Soto-Cruz, 2020). Kinase-library screening also indicates that JAK3 inhibitor VI can inhibit EGFR T790M in resistant contexts (Nishiya et al., 2015), suggesting that this compound may exert broader kinase-targeting effects consistent with suppression of cytokine and growth-factor signalling programmes in the L1000 perturbational landscape.

**Table 6.** Connectivity mapping–derived candidate compounds predicted to reverse gene-specific proteomic tumour states in cervical cancer. . Top-ranked perturbagens identified by L1000FWD connectivity mapping using RPPA-derived elastic-net classifier signatures for recurrently altered genes (SYNE1, DST, MUC5B, and TTN) in the TCGA-CESC cohort. Compounds are ranked by Combined Score, with negative scores indicating predicted reversal of the inferred proteomic state. Z-scores quantify the strength of anti-similarity between the compound-induced transcriptional signature and the tumour-state signature. Nominal p-values and FDR-adjusted q-values are reported to provide statistical context; given the conservative correction applied across perturbagens, q-values are interpreted as prioritisation metrics rather than definitive significance thresholds. Together, these results nominate mechanistically coherent candidate compounds targeting stress-adaptation, signalling, metabolic, and survival pathways associated with distinct gene-defined proteomic programmes.

| rank | pert_id | drug_name | Combined_Score | Z_score | P_value | Q_value |
| --- | --- | --- | --- | --- | --- | --- |
| Top candidate compounds predicted to reverse the SYNE1-associated proteomic signature |  |  |  |  |  |  |
| 1 | BRD-K13049116 | BMS-754807 | -7.82 | 1.82 | 0.0000514 | 0.584 |
| 2 | BRD-K54256913 | MK-1775 | -7.51 | 1.78 | 0.0000603 | 0.584 |
| 3 | BRD-K51967704 | BIIB021 | -4.73 | 1.70 | 0.00165 | 0.584 |
| 4 | BRD-K39120595 | bithionol | -5.45 | 1.86 | 0.00117 | 0.584 |
| 5 | BRD-K82941592 | rosuvastatin | -4.80 | 1.76 | 0.00191 | 0.584 |
| 6 | BRD-U29336476 | XMD-16144 | -4.93 | 1.73 | 0.00140 | 0.584 |
| 7 | BRD-K42918627 | GSK-2126458 | -4.33 | 1.59 | 0.00186 | 0.584 |
| 8 | BRD-K08316444 | rotenone | -4.79 | 1.73 | 0.00169 | 0.584 |
| 9 | BRD-A75409952 | wortmannin | -4.71 | 1.65 | 0.00142 | 0.584 |
| 10 | BRD-K37865504 | LY-2183240 | -4.74 | 1.73 | 0.00180 | 0.584 |
| Top candidate compounds predicted to reverse the DST-associated proteomic signature |  |  |  |  |  |  |
| 1 | BRD-K20742498 | RS-39604 | -11.1 | 1.82 | 0.000000833 | 0.00518 |
| 2 | BRD-K13514097 | everolimus | -10.3 | 1.71 | 0.000000869 | 0.00518 |
| 3 | BRD-K71103788 | duloxetine | -7.97 | 1.74 | 0.0000269 | 0.0112 |
| 4 | BRD-K60623809 | SU-11652 | -8.52 | 1.83 | 0.0000220 | 0.0112 |
| 5 | BRD-K82036761 | sertraline | -8.59 | 1.86 | 0.0000246 | 0.0112 |
| 6 | BRD-A52650764 | ingenol | -8.37 | 1.81 | 0.0000235 | 0.0112 |
| 7 | BRD-K66296774 | fluvastatin | -7.74 | 1.70 | 0.0000272 | 0.0112 |
| 8 | BRD-K82983861 | GW-0742 | -8.21 | 1.80 | 0.0000274 | 0.0112 |
| 9 | BRD-K56301217 | ABT-737 | -7.80 | 1.77 | 0.0000403 | 0.0113 |
| 10 | BRD-K61463582 | BRD-K61463582 | -7.43 | 1.68 | 0.0000384 | 0.0112 |
| Top candidate compounds predicted to reverse the MUC5B-associated proteomic signature |  |  |  |  |  |  |
| 1 | BRD-K40329609 | BRD-K40329609 | -8.44 | 1.75 | 0.0000146 | 0.0192 |
| 2 | BRD-K53780220 | BRD-K53780220 | -8.57 | 1.75 | 0.0000125 | 0.0192 |
| 3 | BRD-K97309399 | thiothixene | -9.58 | 1.88 | 0.00000824 | 0.0192 |
| 4 | BRD-A82371568 | clofarabine | -9.58 | 1.90 | 0.00000914 | 0.0192 |
| 5 | BRD-K04210847 | tamoxifen | -9.53 | 1.87 | 0.00000793 | 0.0192 |
| 6 | BRD-A94413429 | NTNCB | -9.59 | 1.91 | 0.00000922 | 0.0192 |
| 7 | BRD-A73909368 | dactinomycin | -8.41 | 1.70 | 0.0000114 | 0.0192 |
| 8 | BRD-K53423944 | BRD-K53423944 | -5.80 | 1.74 | 0.000461 | 0.0365 |
| 9 | BRD-K93176058 | AC-55649 | -5.95 | 1.78 | 0.000446 | 0.0365 |
| 10 | BRD-K59488055 | BRD-K59488055 | -5.60 | 1.70 | 0.000503 | 0.0365 |
| Top candidate compounds predicted to reverse the TTN-associated proteomic signature |  |  |  |  |  |  |
| 1 | BRD-A31107743 | brefeldin-A | -8.56 | 1.76 | 1.35e-5 | 0.0482 |
| 2 | BRD-K12762134 | XAV-939 | -6.87 | 1.83 | 1.75e-4 | 0.0793 |
| 3 | BRD-K57631554 | aminolevulinic-acid | -6.60 | 1.71 | 1.39e-4 | 0.0793 |
| 4 | BRD-K13533483 | cyclosporin-A | -6.67 | 1.81 | 2.11e-4 | 0.0793 |
| 5 | BRD-A73741725 | exemestane | -6.77 | 1.83 | 1.98e-4 | 0.0793 |
| 6 | BRD-K33551950 | radicicol | -5.87 | 1.68 | 3.20e-4 | 0.0951 |
| 7 | BRD-K60219430 | serdemetan | -7.16 | 1.84 | 1.30e-4 | 0.0793 |
| 8 | BRD-K68558722 | deracoxib | -6.50 | 1.79 | 2.35e-4 | 0.0793 |
| 9 | BRD-K04546108 | JAK3-inhibitor-VI | -6.66 | 1.83 | 2.33e-4 | 0.0793 |
| 10 | BRD-K65955264 | BRD-K65955264 | -6.47 | 1.75 | 1.96e-4 | 0.0793 |

Additional hits are best interpreted as hypothesis-generating. Exemestane (BRD-A73741725), an aromatase inhibitor widely used in breast cancer endocrine therapy, may reflect perturbational signatures pointing to hormone-responsive transcriptional circuitry intersecting with TTN-associated regulatory networks, although cervical cancer-specific mechanisms remain to be validated (Robinson, 2009; Dumas et al., 2025). In contrast, 5-aminolevulinic acid (5-ALA) photodynamic therapy shows direct cervical cancer relevance: topical ALA-PDT promotes HPV clearance and lesion regression in cervical LSIL cohorts, and mechanistic studies in SiHa cells demonstrate suppression of invasion and migration through modulation of a miR-152-3p/JAK1/STAT1 signalling axis (Li et al., 2024; Wang et al., 2024). This observation further supports oxidative stress and cytokine signalling modulation as plausible reversal mechanisms consistent with TTN-associated stress signalling. Collectively, TTN connectivity mapping supports a biologically coherent framework in which TTN-associated cervical cancer states are preferentially reversible by perturbations that (i) collapse trafficking and proteostasis capacity (brefeldin A; radicicol), (ii) suppress developmental signalling circuits (XAV939-mediated Wnt inhibition), (iii) engage stress–checkpoint responses through TP53-axis activation (serdemetan), and (iv) modulate cytokine, inflammatory, and oxidative stress signalling (JAK3 inhibitor VI, cyclosporin A, 5-ALA). These results indicate that the TTN programme is less consistent with single oncogenic pathway dependence and instead reflects reliance on distributed homeostatic buffering networks, making proteostasis disruption and stress-axis engagement particularly promising therapeutic strategies for experimental validation. Network topology analysis further illustrated how predicted reversing perturbagens converge on key regulatory hubs within the TTN-associated proteomic programme **(Figure 13)**. Compounds targeting proteostasis and trafficking (brefeldin A, radicicol) localise near the SRC–ERBB2 signalling module, developmental pathway suppression (XAV939) aligns with the SMAD4/GATA3 transcriptional branch, TP53-axis activation through serdemetan centres on the TP53 checkpoint hub, while oxidative and mitochondrial stress–inducing interventions such as aminolevulinic acid photodynamic therapy and cyclosporin A align with the metabolic–redox module containing G6PD, PRDX1, and TFAM. Together, this network integration reinforces the interpretation that TTN-associated cervical cancer states exhibit vulnerabilities across interconnected stress-buffering pathways rather than a single dominant oncogenic dependency.

**Figure 13.**
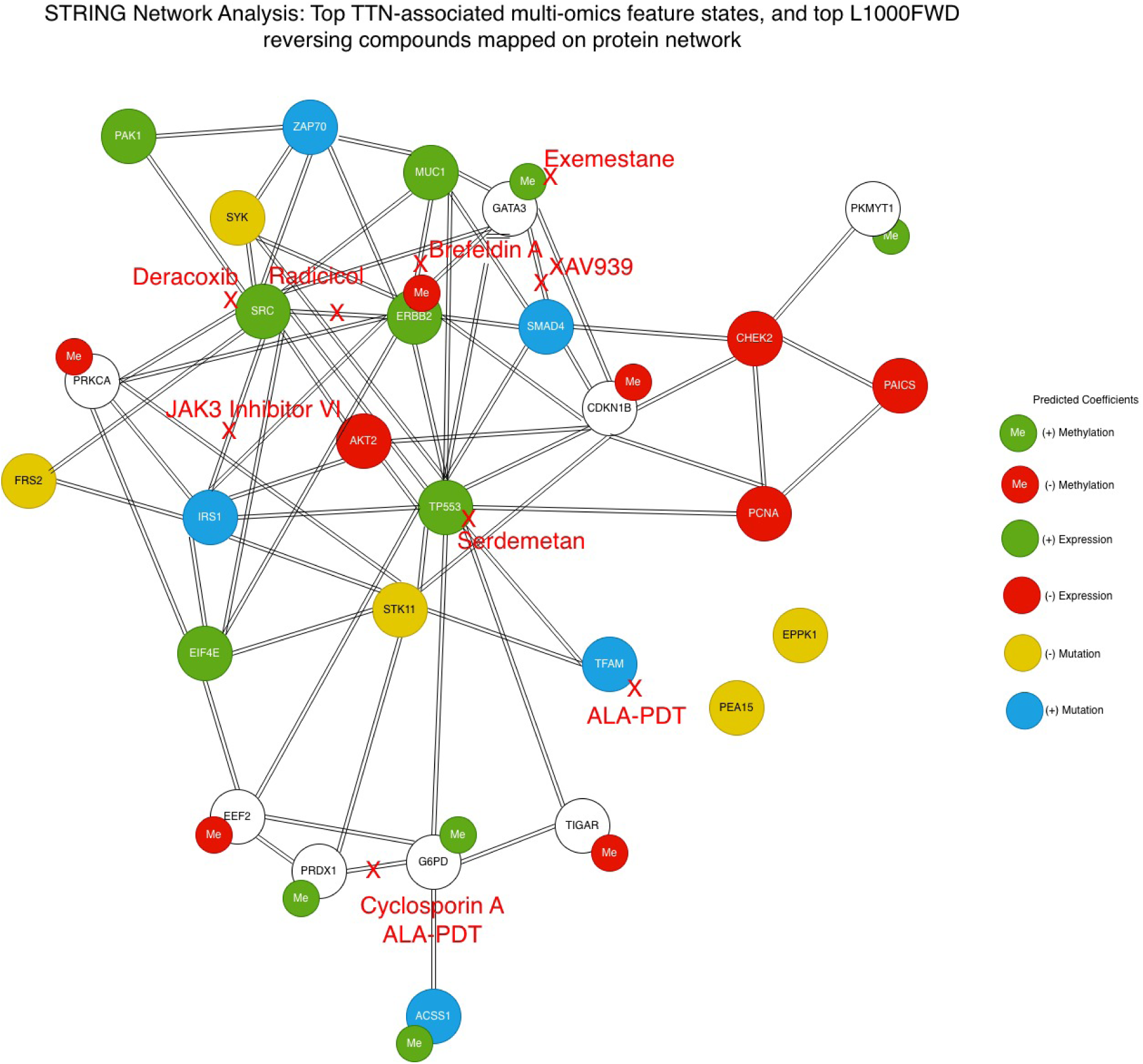
STRING protein interaction network of TTN-associated multi-omics features with predicted reversing perturbagens mapped to mechanistic nodes. A protein–protein interaction network was constructed using STRING to contextualise the top TTN-associated molecular features identified from integrated multi-omics elastic-net modelling. Nodes represent genes from the predictive feature set and are colour-coded according to the direction of model coefficients: green indicates increased expression or methylation, red indicates decreased expression or methylation, yellow denotes negatively associated mutations, and blue denotes positively associated mutations. Edges represent experimentally supported or predicted protein–protein interactions. Predicted therapeutic perturbagens identified by L1000FWD connectivity mapping are overlaid on the network (red labels) and positioned near the most plausible mechanistic targets or pathway modules within the network. These include trafficking/proteostasis stress modulators (brefeldin A, radicicol), developmental signalling suppression (XAV939), stress–checkpoint engagement through the p53 axis (serdemetan), inflammatory or cytokine signalling modulation (deracoxib, JAK3 inhibitor VI), and oxidative or mitochondrial stress inducers (aminolevulinic acid photodynamic therapy and cyclosporin A). Mapping of compounds onto the interaction topology highlights convergence on several functional hubs, including RTK/SRC signalling, p53-mediated stress checkpoints, transcriptional developmental signalling, and metabolic–redox buffering pathways, supporting the hypothesis that TTN-associated tumour states rely on stress-adaptation networks rather than a single oncogenic driver.

#### MUC5B-associated connectivity mapping highlights replication-stress tolerance, adhesive-inflammatory signalling, and endocrine-linked transcriptional control

To translate the MUC5B-associated multi-omics programme into therapeutic hypotheses, we performed L1000FWD connectivity mapping using directionally consistent features derived from the integrated elastic-net models (Mutation AUC = 1.00; Expression AUC = 1.00; Methylation AUC = 0.752), which demonstrated strong proteome-level encoding of mutation and expression states but weaker methylation transmission **(Supplementary Figure S6).** The features used for connectivity analysis were derived from the modality-specific elastic-net coefficient structure defining the MUC5B proteomic programme **(Supplementary Table S9)** and were interpreted in the context of pathway enrichment analyses highlighting VEGF–VEGFR2 signalling, growth-factor signalling, and microenvironmental response pathways **(Figure 9C)**. In addition, transcription-factor regulatory network analysis identified RELA-mediated inflammatory transcriptional activity as a central regulatory node within the MUC5B module **(Figure 10)**. Together, these upstream analyses define a replication-stressed, transcriptionally active tumour state characterised by inflammatory signalling and adaptive microenvironmental responses, providing the biological framework for the downstream connectivity mapping results. The inferred MUC5B state was characterised by coordinated weighting of replication stress and DNA damage signalling (ATR, CHD1L, XPF, Cyclin B1), adhesion–motility and growth-factor circuitry (FAK, PAK1, IRS2), and inflammatory transcriptional control (NFκB/p65), alongside developmental pathway signalling (GLI1) and metabolic transport or rewiring (MCT4, Hexokinase-I). Notably, the presence of ERα among mutation-weighted features further implicates hormone-responsive transcriptional programmes as a defining regulatory axis of this subset. Connectivity mapping identified multiple perturbagens with strong negative connectivity and significant multiple-testing support (Q ≈ 0.019–0.036), nominating convergent vulnerabilities in global transcriptional output, DNA replication processes, and endocrine-linked survival signalling **(Table 6)**. A prominent reversing compound was clofarabine (BRD-A82371568), a purine nucleoside analogue with established cytotoxic activity in hematologic malignancies and emerging evidence of broader antitumour mechanisms. Beyond direct inhibition of DNA synthesis, recent mechanistic work in solid-tumour models shows that clofarabine can trigger a p53–STING interaction, activating a non-canonical STING–NFκB signalling pathway that transcriptionally induces targets including CCL5, CXCL10, HLA genes, and BAX. This response couples apoptosis with GSDME-mediated pyroptosis and promotes immunogenic cell death, ultimately enhancing CD8⁺ T-cell cytotoxic activity (Wu et al., 2025). These mechanisms align closely with the MUC5B tumour programme described in this study: a state enriched for replication and repair pressure would be expected to collapse following nucleoside-analogue–induced replication disruption. Importantly, activation of the STING–NFκB axis provides an additional immunogenic route by which tumour-state reversal may occur, particularly relevant in HPV-associated cancers where innate immune signalling is frequently rewired. Dactinomycin (actinomycin D; BRD-A73909368) also emerged among the strongest antagonistic perturbagens and has direct relevance to cervical cancer biology. In SiHa cervical cancer cells, actinomycin D is a well-established inducer of apoptosis, producing measurable early and late apoptotic biomarkers including Annexin V positivity, mitochondrial membrane potential loss, caspase-3/7 activation, and DNA damage (Punchoo et al., 2021). More broadly, dactinomycin has been recognised as a potent inducer of immunogenic cell death (ICD). Comparative analyses of ICD-inducing agents suggest that inhibition of RNA synthesis—often accompanied by secondary inhibition of translation—represents a common initiating event leading to pre-mortem stress signalling capable of supporting immune-dependent anticancer activity in vivo (Humeau et al., 2020). Together, these findings support the interpretation that the MUC5B-associated tumour state may rely on high transcriptional throughput and stress-adaptive programmes, rendering global transcriptional blockade an effective strategy for collapsing network-level homeostasis rather than targeting individual signalling nodes. Tamoxifen (BRD-K04210847), a selective estrogen receptor modulator, also appeared among the top reversing perturbagens. The most defensible interpretation of this signal is not that tamoxifen represents a direct cervical cancer therapeutic; rather, it suggests the presence of a hormone-responsive transcriptional component intersecting with tumour regulatory circuitry. Supporting this interpretation, colposcopy and biopsy data from HPV-positive individuals indicate that tamoxifen use was not associated with increased abnormal cervical histopathology in a breast cancer cohort, supporting its gynaecologic safety profile in screened populations (Cetin et al., 2024). Accordingly, tamoxifen is best interpreted here as a perturbational indicator of ER-linked transcriptional wiring rather than evidence of direct cervical anticancer efficacy.

Thiothixene (BRD-K97309399), a dopamine receptor antagonist, produced a strong reversing signature and suggests a distinct immune-microenvironment mechanism. Recent work shows that thiothixene stimulates efferocytosis, the macrophage-mediated clearance of apoptotic cells, and promotes sustained efferocytic activity through upregulation of a retinol-binding protein receptor signalling axis (Stra6L → arginase 1), whereas dopamine inhibits this process (Kojima et al., 2025). Although not a conventional oncology agent, this mechanism provides a plausible cancer-relevant interpretation for the reversal signal: modulation of tumour–myeloid clearance pathways could influence immune tone and the downstream consequences of tumour cell death, particularly when combined with perturbagens capable of inducing immunogenic death programmes such as clofarabine or dactinomycin. AC-55649 (BRD-K93176058) is best described cautiously as a retinoic acid receptor (RAR) isoform-selective retinoid-class compound. Retinoid signalling broadly regulates growth arrest, differentiation, and apoptosis, and modern RAR agonist development increasingly emphasises isoform selectivity and improved pharmacological properties (Lund et al., 2005; Borthwick et al., 2020). Within the present framework, this signal suggests that differentiation-state rewiring may counter the stress-adapted MUC5B tumour programme; however, this interpretation should be considered hypothesis-generating unless validated experimentally in cervical cancer models. Several additional high-ranking perturbagens remain poorly annotated in current pharmacological resources (e.g., BRD-A94413429 labelled “NTNCB,” BRD-K40329609, BRD-K53780220, BRD-K53423944, and BRD-K59488055). Rather than speculative interpretation, these compounds are best positioned as empirically observed reversing perturbational profiles that warrant further target deconvolution and annotation expansion through resources such as LINCS, CTRP, or PubChem to identify the molecular nodes responsible for reversal. Taken together, the MUC5B connectivity results prioritise three mechanistically coherent vulnerability axes: (i) replication stress disruption with immunogenic amplification (clofarabine; apoptosis/pyroptosis/ICD via p53–STING–NFκB signalling and CD8⁺ T-cell engagement), (ii) global transcriptional suppression with immunogenic cell death potential (dactinomycin/actinomycin D; transcription inhibition as a central ICD mechanism), and (iii) microenvironmental modulation influencing tumour clearance and immune tone (thiothixene-driven efferocytosis). Retinoid/RAR signals (AC-55649) and endocrine-linked transcriptional signals (tamoxifen) are best interpreted as state-rewiring hypotheses requiring targeted validation in MUC5B-stratified cervical cancer models. Top MUC5B-associated features identified by elastic-net modelling were subsequently submitted to STRING to construct a protein–protein interaction network in which nodes represent proteins and edges indicate documented or predicted interactions. Node annotations reflect multi-omic states (expression, methylation, and mutation), and predicted sites of perturbation by top L1000FWD reversing compounds were mapped onto relevant network branches. The resulting interaction landscape highlights programme-specific signalling hubs linking replication stress and DNA repair (clofarabine), transcriptional regulation (dactinomycin, tamoxifen), and developmental signalling pathways (AC-55649), providing a network-based framework integrating multi-omic inference with functional protein interactions and mechanistically guided pharmacological hypotheses across the MUC5B programme **(Figure 14)**.

**Figure 14.**
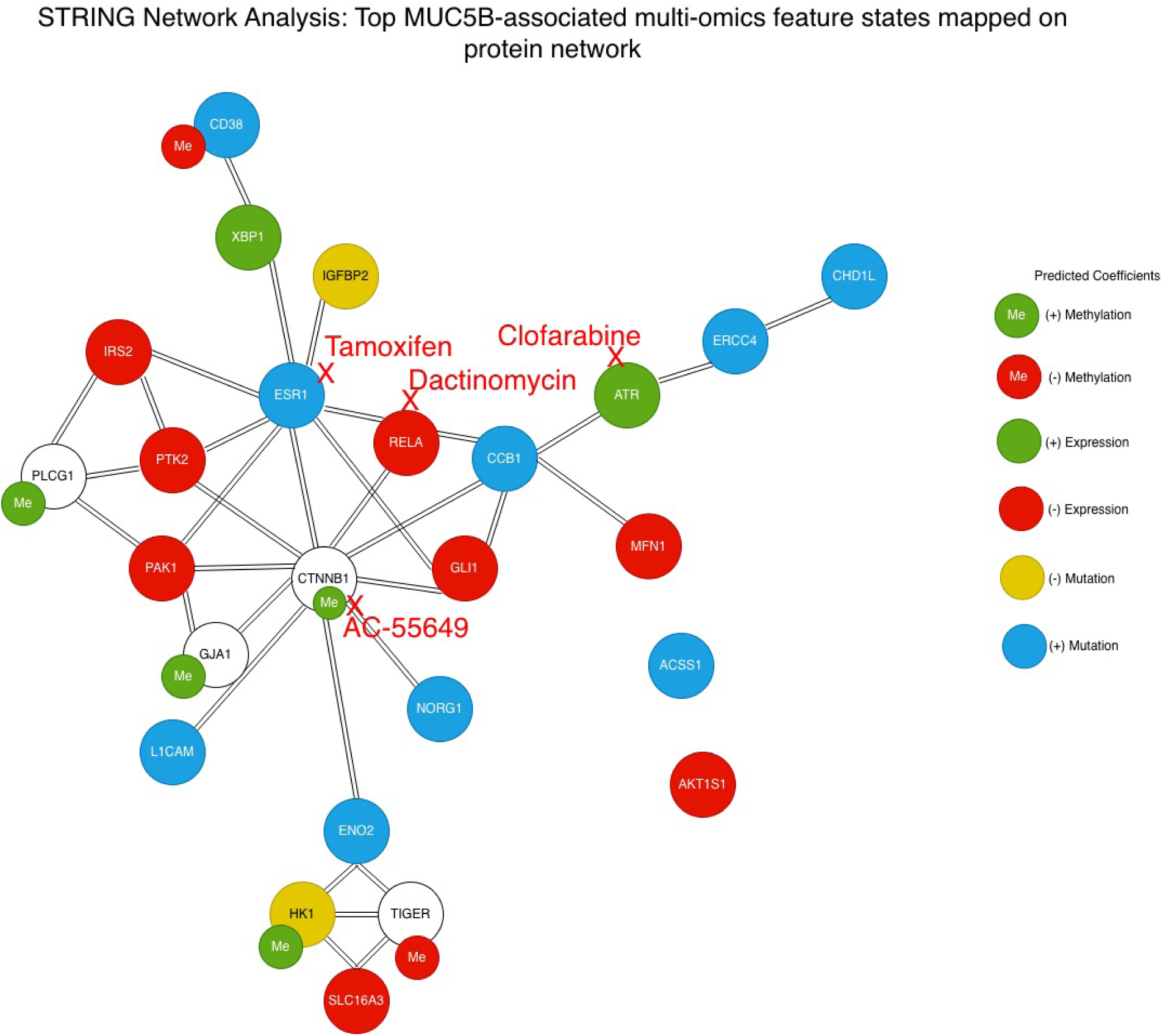
STRING network showing predicted drug action on the MUC5B-associated multi-omics programme in cervical cancer. Protein–protein interaction network constructed from the top MUC5B-associated multi-omics features using STRING. Nodes represent proteins and edges represent known or predicted protein–protein interactions. Clofarabine is mapped to the replication stress and DNA repair axis (ATR–ERCC4–CHD1L), consistent with its nucleoside analogue mechanism disrupting DNA synthesis and activating p53–STING–NFκB signalling. Dactinomycin (actinomycin D) targets the transcriptional module centred on ESR1 and RELA, reflecting inhibition of RNA synthesis and induction of immunogenic cell death pathways. Tamoxifen is positioned at the ESR1 node, highlighting the endocrine-linked transcriptional component of the MUC5B programme. AC-55649 maps to the developmental signalling branch involving CTNNB1 and GLI1, suggesting a potential differentiation-based mechanism for network reversal. Together, the network illustrates how predicted perturbagens converge on replication stress, transcriptional regulation, and developmental signalling pathways within the MUC5B-associated tumour state. Node colours represent multi-omics annotations: green, positive expression; red, negative expression; green “Me”, positive methylation; red “Me”, negative methylation; yellow, negative mutation; and blue, positive mutation. “+” indicates positive coefficients and “−” indicates negative coefficients. Red labels, arrows, and crosses indicate predicted pathway-level sites of perturbation by reversing compounds.

#### SYNE1-associated connectivity mapping implicates growth-factor/PI3K rewiring, checkpoint fragility, and proteostasis dependence

To translate the SYNE1-associated multi-omics programme into therapeutic hypotheses, we performed L1000FWD connectivity mapping using directionally consistent features derived from integrated elastic-net models (Mutation AUC = 1.00; Expression AUC = 0.999; Methylation AUC = 0.736), which demonstrated strong proteomic discrimination of SYNE1 mutation and expression states but weaker methylation encoding **(Supplementary Figure S6)**. The underlying signalling architecture and pathway context for these features were previously characterised through Reactome enrichment and transcription-factor regulatory analysis **(Figure 9A**; **Figure 10**; **Table 4)**. The inferred SYNE1 state combined: (i) oncogenic growth-factor signalling (BRAF, PDGFRB, NFκB/p65) with PTEN suppression, (ii) DNA maintenance/replication wiring (MSH2, Cdc6), and (iii) a pattern consistent with checkpoint attenuation (CDK1, Cyclin B1, CHK1, CDC25C), as defined by elastic-net feature loading patterns and pathway enrichment analyses. Together, these features suggest a tumour state supported by survival signalling and replication-stress tolerance, but potentially vulnerable to forced checkpoint failure and multi-node destabilisation of signalling networks. In contrast to the DST, TTN, MUC5B and PIK3CA (methylation) runs, the top SYNE1 reversing signatures showed modest multiple-testing support (Q ≈ 0.584 across the top hits), so the results are best framed as mechanism-guided hypothesis generation rather than statistically prioritised drug claims **(Table 6)**. Even so, the leading perturbagens are mechanistically coherent with the inferred biology and converge on three actionable axes: growth-factor and PI3K-pathway dependence; checkpoint fragility and replication-stress exploitation; and proteostasis-mediated signalling destabilisation. The strongest reversing compound was BMS-754807 (BRD-K13049116), a potent dual IGF-1R/insulin receptor (IR) kinase inhibitor that targets an upstream growth-factor axis feeding directly into PI3K–AKT and MAPK signalling. While the foundational pharmacology and antitumour activity of BMS-754807 were established across multiple epithelial and mesenchymal models, subsequent mechanistic work has shown that IGF pathway inhibition can induce replication stress (via RRM2/dNTP depletion) and creates synthetic-lethal vulnerability with checkpoint blockade (e.g., CHK1 or WEE1 co-inhibition), providing a biologically coherent rationale for combining IGF-axis suppression with cell-cycle stressors in solid tumours (Carboni et al., 2009; Wu et al., 2021). Within the cohort, this aligns tightly with the inferred SYNE1 proteomic programme—notably PTEN↓ with BRAF↑/PDGFRB↑/ NFκB↑—a configuration in which sustained upstream receptor input can maintain survival signalling and transcriptional output, consistent with the growth-factor signalling enrichment identified in the SYNE1 pathway analysis. Consistent with this dependency on PI3K-pathway wiring, multiple additional SYNE1 hits implicated direct PI3K/mTOR suppression. GSK-2126458 (dactolisib) is a highly potent dual PI3Kα/mTOR inhibitor with broad antitumour activity in vivo, supporting pathway-level reversal of PI3K-driven survival particularly when PTEN restraint is reduced (Knight et al., 2010; Huang et al., 2023). Wortmannin, a canonical PI3K inhibitor (with broader PI3K-family activity in some contexts), further reinforces PI3K-pathway reversal as a consistent theme; in cervical cancer–relevant cellular contexts, wortmannin-related natural products have also been shown to attenuate NF-κB signalling and induce oxidative stress–linked apoptosis in HeLa cervical cells, supporting the plausibility of PI3K/NF-κB-coupled vulnerability axes in HPV-driven disease (Acuña et al., 2013). A second prominent reversing perturbagen was MK-1775/adavosertib/AZD1775 (BRD-K54256913), a first-in-class WEE1 inhibitor. WEE1 restrains CDK1/2 to enforce the G2/M checkpoint; inhibition can force unscheduled mitotic entry and lethal replication catastrophe, especially in tumours already operating under replication stress. This mechanism is directly concordant with the SYNE1 state’s coordinated suppression of core G2/M regulators (CDK1↓, Cyclin B1↓, CDC25C↓, CHK1↓), consistent with reliance on residual checkpoint buffering as suggested by the transcription-factor regulatory network analysis highlighting CHEK1, CDK1, and CCNB1 as central nodes in the SYNE1 regulatory programme. Importantly, beyond single-agent rationale, early clinical experience in gynecologic chemoradiation (including predominantly cervical cancer patients) supports feasibility signals while highlighting overlapping hematologic and gastrointestinal toxicities that must be managed when pairing WEE1 blockade with DNA-damaging therapy (Do et al., 2015; Gonzalez-Ochoa et al., 2023).

BIIB021 (BRD-K51967704), an orally bioavailable HSP90 inhibitor, was also among the top antagonistic hits and is particularly relevant to cervical cancer biology: BIIB021 has demonstrated nanomolar cytotoxicity in HeLa cells, with biochemical evidence of HSP90 inhibition and induction of intrinsic apoptosis (BCL-2↓ with BAX/CYTO-c/CASP3↑), supporting a plausible “network-collapse” strategy for SYNE1-linked states exhibiting multi-pathway activation (Güven & Özgür, 2023). This interpretation is consistent with the interconnected signalling architecture observed in the SYNE1 protein–protein interaction network, where growth-factor signalling, checkpoint regulation, and metabolic pathways converge **(Figure 15)**. Several additional hits are best framed as state stressors that may expose metabolic or signalling fragility rather than as immediate cervical-cancer drug claims. Rotenone (BRD-K08316444), a mitochondrial complex I inhibitor, has been shown to suppress cancer cell motility/EMT and downregulate AKT/mTOR phosphorylation, with reversal of IGF-1–driven signalling in cancer models—mechanistically consistent with targeting PI3K–AKT–mTOR–linked survival states, though careful toxicity considerations apply (Heinz et al., 2017; Xiao et al., 2020). Rosuvastatin (BRD-K82941592) nominates the mevalonate/cholesterol axis as a modulatory lever for membrane-associated signalling; however, population-level evidence for statins in cervical cancer prevention is inconsistent, so this should be treated as a hypothesis about pathway modulation rather than disease-specific efficacy (Bajaj et al., 2025; Löfling et al., 2025). LY-2183240 (endocannabinoid uptake/FAAH-pathway inhibitor) and bithionol are supported primarily by non-cervical cancer evidence and should be interpreted as exploratory repurposing candidates within perturbational space rather than cervical-validated agents (Marzęda et al., 2024; Załuska-Ogryzek et al., 2025; Ayyagari & Brard, 2014; Ayyagari et al., 2016). XMD-16144 (BRD-U29336476) remains under-annotated and is best prioritised for target deconvolution before mechanistic interpretation (Liu et al., 2025). Overall, SYNE1 connectivity mapping supports a coherent, cervical cancer–relevant model in which SYNE1-associated tumours may be preferentially reversible by: (i) blocking upstream growth-factor input and PI3K–mTOR survival signalling (BMS-754807, dactolisib, wortmannin); (ii) exploiting checkpoint dependence with forced mitotic entry (adavosertib); and (iii) destabilising oncogenic signalling networks via proteostasis disruption (BIIB021). Given the modest multiple-testing support for SYNE1 (Q ≈ 0.584), these agents are best presented as mechanism-guided candidates for prioritised validation in SYNE1-stratified cervical cancer models rather than as statistically definitive drug nominations **(Table 6)**. Top SYNE1-associated features identified by elastic-net modelling were submitted to STRING, generating networks in which nodes represent proteins and edges indicate documented or predicted interactions. Node annotations reflect multi-omics states (expression, methylation, and mutation), and predicted sites of perturbation by top L1000FWD reversing compounds were mapped onto relevant network branches. The resulting interaction map highlights programme-specific signalling hubs linking growth-factor/PI3K signalling, checkpoint and replication stress modules, and additional metabolic and inflammatory pathways within the SYNE1-associated tumour state **(Figure 15)**.

**Figure 15.**
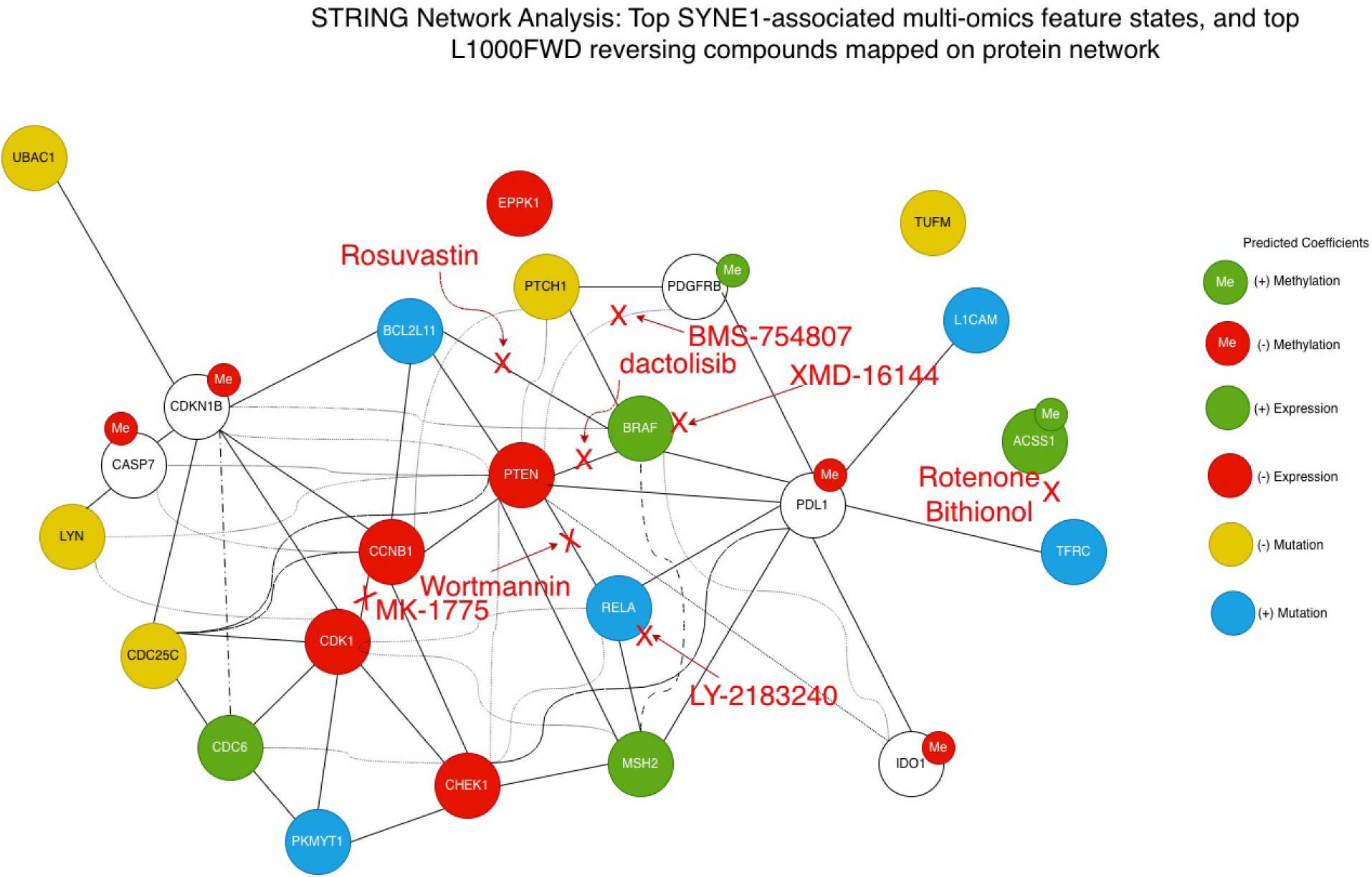
STRING protein–protein interaction network of SYNE1-associated multi-omics features with predicted sites of action for reversing compounds. Protein–protein interaction network constructed using STRING for the top SYNE1-associated multi-omics features identified by elastic-net modelling. Nodes represent proteins and edges indicate known or predicted protein–protein interactions. BMS-754807 and dactolisib target upstream growth-factor and PI3K signalling, converging on the PDGFRB–BRAF–PTEN axis. MK-1775 (adavosertib) targets the CDK1–Cyclin B1 checkpoint module, exploiting checkpoint fragility and replication stress. Additional perturbagens including wortmannin, rosuvastatin, rotenone, bithionol, LY-2183240, and XMD-16144 highlight potential vulnerabilities in PI3K signalling, metabolic pathways, inflammatory signalling, and membrane-associated signalling processes. Together, the network illustrates how predicted therapeutic perturbations converge on the interconnected signalling modules defining the SYNE1 tumour state in cervical cancer. Node colours represent multi-omics annotations: green, positive expression; red, negative expression; green “Me”, positive methylation; red “Me”, negative methylation; yellow, negative mutation; and blue, positive mutation. “+” indicates positive coefficients and “−” indicates negative coefficients. Red arrows and crosses indicate predicted sites where top L1000FWD reversing compounds may perturb the network.

#### DST-associated connectivity mapping implicates mTOR/RTK signalling, apoptotic priming, and metabolic-immune rewiring

To translate the DST-associated multi-omic programme into therapeutic hypotheses, we performed L1000FWD connectivity mapping using directionally consistent proteomic features derived from the integrated elastic-net models (Mutation AUC = 0.820; Expression AUC = 0.983; Methylation AUC = 1.000; **Supplementary Figure S6)**. These features correspond to the modality-specific coefficient structure defining the DST-associated proteomic state **(Supplementary Table S10)** and were interpreted in the context of pathway enrichment analyses highlighting cytoskeletal organisation, receptor tyrosine kinase signalling, integrin-mediated adhesion, and immune-modulatory pathways **(Figure 9D)**. In addition, transcription factor regulatory network analysis identified regulatory coupling between receptor signalling and chromatin-associated stress-response pathways, including EGFR gain-of-function signalling and epigenetic modulation of ESR1 and HSPA5 **(Figure 10)**. Together, these upstream analyses define a DST-associated tumour state characterised by cytoskeletal perturbation, receptor tyrosine kinase–driven signalling, angiogenic interaction, metabolic rewiring, and immune modulation, providing the biological framework for downstream connectivity mapping. The inferred DST state was dominated by growth-factor and MAPK/PI3K signalling (EGFR, BRAF, AKT2, IRS1, p90RSK), proliferative and mitotic regulators (Cyclin E1, TRIP13), angiogenesis and extracellular remodelling cues (CD31, PAI1), and immune evasion features (PD-L1), together with suppression of innate immune and DNA surveillance modules (STING, IRF3, MSH2). Connectivity mapping identified multiple perturbagens with strong negative connectivity and statistically significant FDR-adjusted values (Q = 0.005–0.011), nominating convergent vulnerabilities in RTK–mTOR signalling control, apoptosis regulation, and lipid–inflammatory homeostasis **(Table 6)**. Among the most strongly reversing and mechanistically interpretable perturbagens was everolimus (BRD-K13514097), an allosteric mTOR inhibitor that suppresses mTORC1 signalling through FKBP12 binding. In cervical cancer, aberrant activation of the PI3K–AKT–mTOR axis is a recurrent feature driven both by genomic alterations and by HPV oncoprotein activity. Preclinical and early-phase clinical studies have demonstrated that everolimus inhibits cervical cancer cell proliferation, impairs tumour-associated angiogenesis, and sensitises tumour cells and vasculature to cisplatin and radiotherapy, partly through modulation of HPV E7–dependent signalling and mTOR-driven survival pathways (de Melo et al., 2016). Given the prominence of AKT2- and IRS1-linked survival signalling within the DST-associated proteomic programme, mTOR blockade provides a direct and biologically coherent pathway-level strategy to oppose this state. A second high-confidence hit, SU-11652 (BRD-K60623809), is a multitarget receptor tyrosine kinase inhibitor with documented activity against FLT3 and broader RTK networks including VEGFR, PDGFR, and KIT. Although originally characterised in hematologic malignancies, SU-11652 has been shown to exert potent cytotoxic effects in cervical carcinoma cells, including HeLa models, through disruption of RTK-driven survival signalling, induction of apoptosis, and lysosomal destabilisation mechanisms independent of classical apoptosis pathways (Guo et al., 2012; Ellegaard et al., 2013). This aligns closely with the DST signature’s strong dependence on RTK signalling (EGFR, KIT) and angiogenic features (CD31, PAI1), supporting RTK inhibition as a rational vulnerability in DST-stratified cervical tumours.

DST connectivity mapping also highlighted liabilities consistent with apoptotic buffering and stress tolerance. ABT-737 (BRD-K56301217), a BH3 mimetic that antagonises pro-survival BCL-2 family proteins, emerged as a robust reversing perturbagen. In cervical cancer, BCL-2 expression has been associated with disease stage and treatment resistance, and preclinical studies have demonstrated that ABT-737 synergises with irradiation to induce mitochondrial depolarisation, reactive oxygen species accumulation, and caspase-dependent apoptosis in cervical cancer cell lines (Shen et al., 2019). These findings support the hypothesis that DST-associated tumours rely on anti-apoptotic buffering to sustain proliferative and checkpoint stress. In parallel, the presence of fluvastatin (BRD-K66296774) implicates dependence on the mevalonate and cholesterol biosynthesis pathway, which supports membrane organisation, receptor signalling, and oncogenic fitness. Statins have been shown to induce p21 expression, DNA damage responses, and growth inhibition in cervical cancer cells through both p53-dependent and p53-independent mechanisms, and to modulate sensitivity to combination therapies (Lin et al., 2019). This suggests that metabolic pathway interference may destabilise RTK- and PI3K-driven signalling scaffolds in DST-defined states. Additional antagonistic compounds reinforce a model in which DST tumours integrate oncogenic signalling with immune and metabolic regulation. GW-0742 (BRD-K82983861), a selective PPARβ/δ agonist, points to lipid and inflammatory transcriptional control as a modulatory axis capable of reversing the DST programme. Emerging evidence indicates that PPAR signalling intersects with tumour metabolism, immune evasion, and survival pathways relevant to gynaecologic malignancies (Gogola-Mruk et al., 2024). Meanwhile, RS-39604 (BRD-K20742498), a selective 5-HT4 receptor antagonist, nominates serotonergic GPCR signalling as an additional regulatory layer. Serotonin signalling has increasingly been implicated in tumour growth, immune modulation, and angiogenesis, and serotonergic-targeted agents have been proposed as repurposable candidates in cancer contexts, including cervical carcinoma (Hegde et al., 1995; Chen et al., 2024). Finally, ingenol (BRD-A52650764), a PKC modulator, supports sensitivity to perturbation of PKC-dependent stress and signalling nodes that interface with MAPK and NF-κB circuitry—pathways known to be active in HPV-driven cervical cancer and associated with immune evasion and inflammatory signalling. Collectively, DST connectivity mapping prioritises actionable therapeutic strategies centred on RTK–mTOR pathway inhibition (everolimus; SU-11652), apoptotic priming (ABT-737), and metabolic–inflammatory transcriptional rewiring (fluvastatin; GW-0742), with additional support for GPCR- and PKC-linked network perturbation (RS-39604; ingenol). Together, these hypotheses define a coherent, cervical cancer–relevant framework for experimental validation in DST-stratified models, particularly in tumours characterised by EGFR/BRAF/AKT2 activation and PD-L1–associated immune evasion. To further contextualise these therapeutic hypotheses within a systems framework, the top DST-associated proteomic features identified by elastic-net modelling were submitted to STRING to generate protein–protein interaction networks. In these networks, nodes represent proteins and edges indicate documented or predicted functional interactions. Node annotations reflect the contributing multi-omic states (expression, methylation, and mutation), and predicted sites of perturbation by top L1000FWD reversing compounds were mapped onto relevant network branches. The resulting network highlights programme-specific signalling hubs, including growth-factor/PI3K signalling (everolimus, SU-11652, fluvastatin), metabolic modules (GW-0742), stress-response pathways (duloxetine, sertraline), innate immune signalling (ingenol), and anti-apoptotic survival networks (ABT-737). This network-based framework integrates multi-omic features with functional interaction architecture and mechanistically guided pharmacological hypotheses across the DST programme **(Figure 16)**.

**Figure 16.**
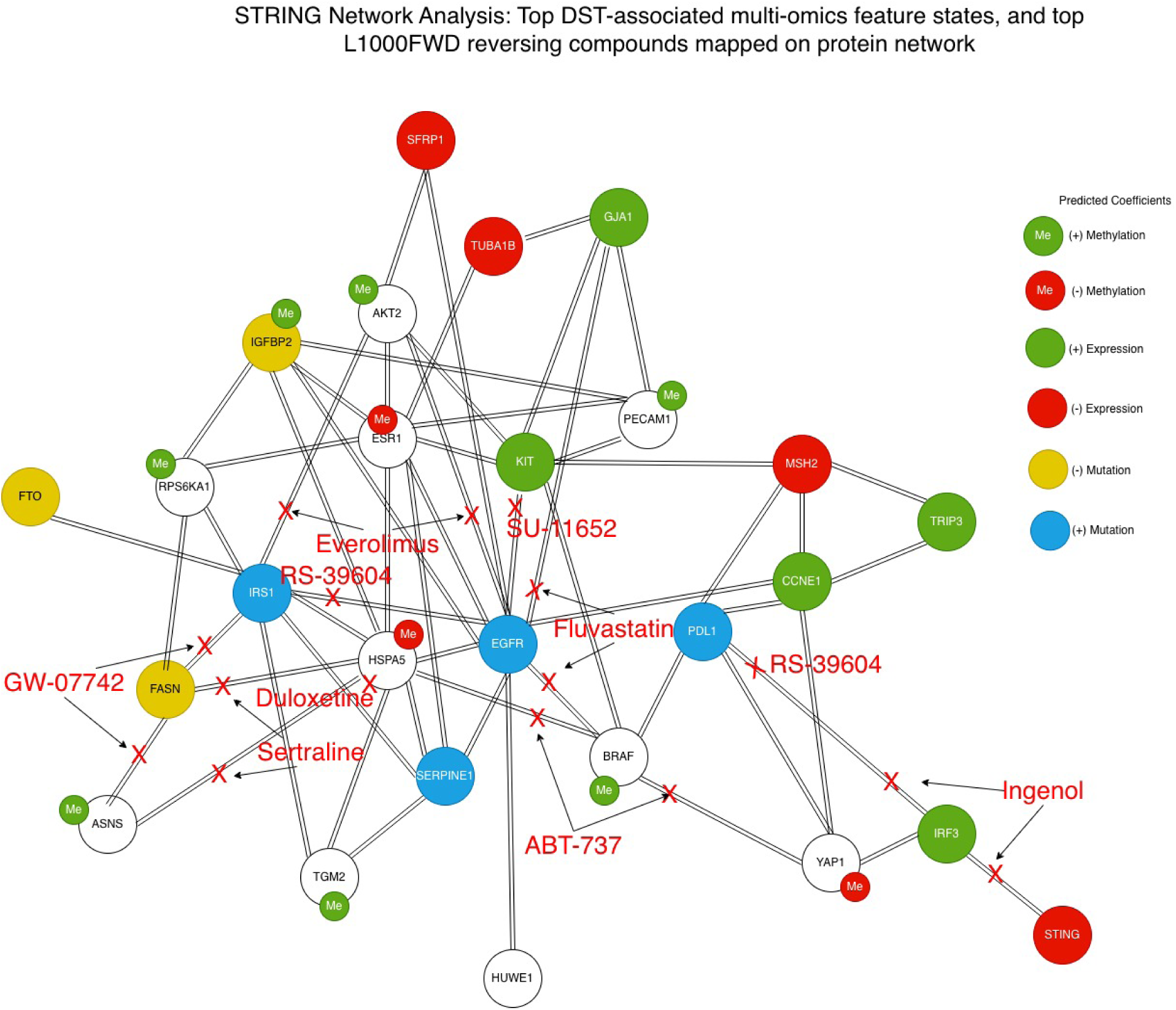
STRING protein–protein interaction network of DST-associated multi-omics features with predicted sites of action for reversing compounds. Protein–protein interaction network constructed using STRING for the top DST-associated multi-omics features identified by elastic-net modelling. Nodes represent proteins and edges represent known or predicted protein–protein interactions. Everolimus targets the IRS1–AKT2 growth signalling branch, SU-11652 and fluvastatin map to the KIT/EGFR receptor signalling axis, GW-0742 perturbs the FASN–ASNS metabolic module, and duloxetine and sertraline affect the HSPA5-associated stress response network. Ingenol is positioned on the IRF3–STING innate immune signalling axis, while ABT-737 targets the central anti-apoptotic survival network. Together, the network illustrates how predicted perturbagens converge on growth signalling, metabolic regulation, immune signalling, and apoptosis pathways within the DST-associated tumour state. Node colours represent multi-omics annotations: green, positive expression; red, negative expression; green “Me”, positive methylation; red “Me”, negative methylation; yellow, negative mutation; and blue, positive mutation. “+” indicates positive coefficients and “−” indicates negative coefficients. Red labels, arrows, and crosses indicate predicted pathway-level sites of perturbation by reversing compounds.

## Discussion

### Clinical Implications

Clinically, integrating RPPA-derived tumour-state classifiers with L1000FWD connectivity mapping provides a pragmatic route to functional stratification in TCGA-CESC. Rather than treating somatic mutation calls as direct therapeutic surrogates, recurrent alterations are translated into proteome-defined tumour “states” that can be interrogated in perturbational space to nominate mechanistically coherent, state-reversing candidates for prioritised validation **(Table 7)**. This strategy is particularly relevant in cervical cancer, where oncogenic HPV biology, PI3K-pathway activation, inflammatory signalling, and stress-adaptation circuitry frequently intersect, complicating single-marker treatment assignment. For the PIK3CA methylation-defined state (AUC = 1.00), the strongest reversal signals converge on replication-associated DNA damage induction (SN-38; irinotecan-class topoisomerase I inhibition) and inflammatory/prostaglandin-axis suppression (valdecoxib; COX-2 inhibition), with complementary vulnerability routes involving developmental pathway inhibition (erismodegib; Hedgehog blockade) and oxidative or mitochondrial stress induction (artesunate). Given modest multiple-testing support (Q ≈ 0.296), these findings should be interpreted as hypothesis-led repurposing signals rather than treatment recommendations. Nevertheless, the nominated mechanisms align with cervical cancer–relevant experimental evidence: SN-38 induces apoptosis in HeLa and SiHa cells via an Akt–p53/p21 axis with partial rescue by forced Akt activation, consistent with selective vulnerability of PI3K-linked states to combined replication stress and survival pathway collapse (Liu et al., 2009), and targeted SN-38 delivery enhances uptake and apoptosis in HeLa and CaSki models (Zheng et al., 2023). Valdecoxib’s relevance can be cautiously justified through both broader oncologic repurposing discussions and COX-independent effects on membrane lipid order that may modulate STAT3/NF-κB-aligned inflammatory survival signalling (İnan Genç et al., 2017; El-Malah et al., 2022). Developmental wiring vulnerability is supported by the pharmacology of SMO antagonism (Pan et al., 2010) and evidence that Hedgehog blockade can cooperate with pathway inhibitors in solid tumours (D’Amato et al., 2014), while artesunate shows direct cervical cancer evidence for ROS induction, mitochondrial depolarisation, and pro-apoptotic Bcl-2 family shifts (Sadana et al., 2025), with additional HPV-relevant translational motivation (Zhang et al., 2024). Across other programmes, **Table 7** highlights a consistent “stress-collapsing” logic, albeit with gene-specific entry points. TTN-associated states (Q ≈ 0.048–0.095) preferentially nominate secretory trafficking and proteostasis disruption (brefeldin A; radicicol/HSP90 inhibition), developmental wiring suppression (XAV-939; Wnt/β-catenin inhibition), and checkpoint or stress-axis enforcement (serdemetan; MDM2–p53 modulation). Mechanistically, brefeldin A induces acute ER–Golgi collapse and organelle stress (Wood et al., 1991), with cervical cancer–relevant derivatives showing antiproliferative activity and mitochondrial apoptosis signatures in HeLa models (Wang et al., 2023). Wnt pathway inhibition aligns with radiosensitisation and enhanced DNA damage signalling in HeLa cells (Wang et al., 2023), while stress-axis engagement is supported by p53/p21-linked radiosensitiser behaviour for serdemetan (Chargari et al., 2011). Complementary evidence further supports oxidative and mitochondrial stress coupling as a viable lever in cervical cancer contexts (Kooshan et al., 2025) and underscores the relevance of JAK/STAT signalling to HPV-associated disease biology (Gutiérrez-Hoya & Soto-Cruz, 2020). MUC5B-associated states (Q ≈ 0.019–0.036) concentrate on replication stress disruption and global transcriptional output suppression, with additional signals implicating endocrine-linked transcriptional control and microenvironmental modulation. Clofarabine is notable for coupling replication collapse to innate immune activation via p53–STING–NF-κB signalling, promoting immunogenic cell death and CD8⁺ T-cell engagement (Wu et al., 2025). Transcriptional blockade via dactinomycin is supported by direct apoptosis induction in SiHa cells (Punchoo et al., 2021) and broader immunogenic cell-death literature identifying RNA synthesis inhibition as a common initiating event (Humeau et al., 2020). Tamoxifen is best interpreted here as a marker of endocrine transcriptional rewiring rather than a direct cervical cancer efficacy claim, while thiothixene suggests immune-tone modulation via macrophage efferocytosis (Kojima et al., 2025). SYNE1-associated states (Q ≈ 0.584) remain statistically exploratory but converge on a clinically intelligible triad of growth-factor/PI3K dependence, checkpoint fragility, and proteostasis-based network destabilisation. The leading hit BMS-754807 (dual IGF-1R/IR inhibition) aligns with foundational pharmacology and antitumour activity (Carboni et al., 2009) and with mechanistic work linking IGF pathway inhibition to replication stress and checkpoint-linked synthetic lethality (Wu et al., 2021). Complementary support comes from dual PI3K/mTOR inhibition (Knight et al., 2010; Huang et al., 2023), WEE1-mediated checkpoint abrogation in gynaecologic contexts (Do et al., 2015; Gonzalez-Ochoa et al., 2023), and HSP90 inhibition–driven network collapse with HeLa-specific apoptosis evidence (Güven & Özgür, 2023). Collectively, SYNE1 is best positioned as a mechanism-driven validation programme rather than a statistically prioritised one. Finally, DST represents the most statistically robust connectivity programme (Q ≈ 0.005–0.011), making it the most defensible entry point for early translational testing within this framework. DST reversal candidates converge on RTK–mTOR suppression (everolimus; SU-11652), apoptotic priming (ABT-737), and metabolic or inflammatory modulation. Cervical cancer relevance is supported by evidence that mTOR inhibition suppresses proliferation and sensitises tumours to cisplatin or radiotherapy (de Melo et al., 2016), RTK inhibition induces cytotoxicity and lysosomal destabilisation in cervical carcinoma models (Guo et al., 2012; Ellegaard et al., 2013), and BH3 mimetics enhance radiosensitivity via mitochondrial apoptosis (Shen et al., 2019). Statin-linked metabolic scaffolding modulation further supports DNA damage and growth-inhibitory responses in cervical cancer cells (Lin et al., 2019). In sum, the clinical implication of the RPPA→state→connectivity workflow is a tiered translational roadmap **(Table 7)**: (i) prioritise statistically supported programmes (DST, MUC5B, TTN) for first-pass functional validation; (ii) treat lower-support states (PIK3CA methylation, SYNE1) as biologically coherent, mechanism-led hypothesis generators; and (iii) operationalise each state using matched pharmacodynamic readouts aligned to its inferred liabilities, enabling rational, state-stratified therapeutic discovery in cervical cancer.

**Table 7.**
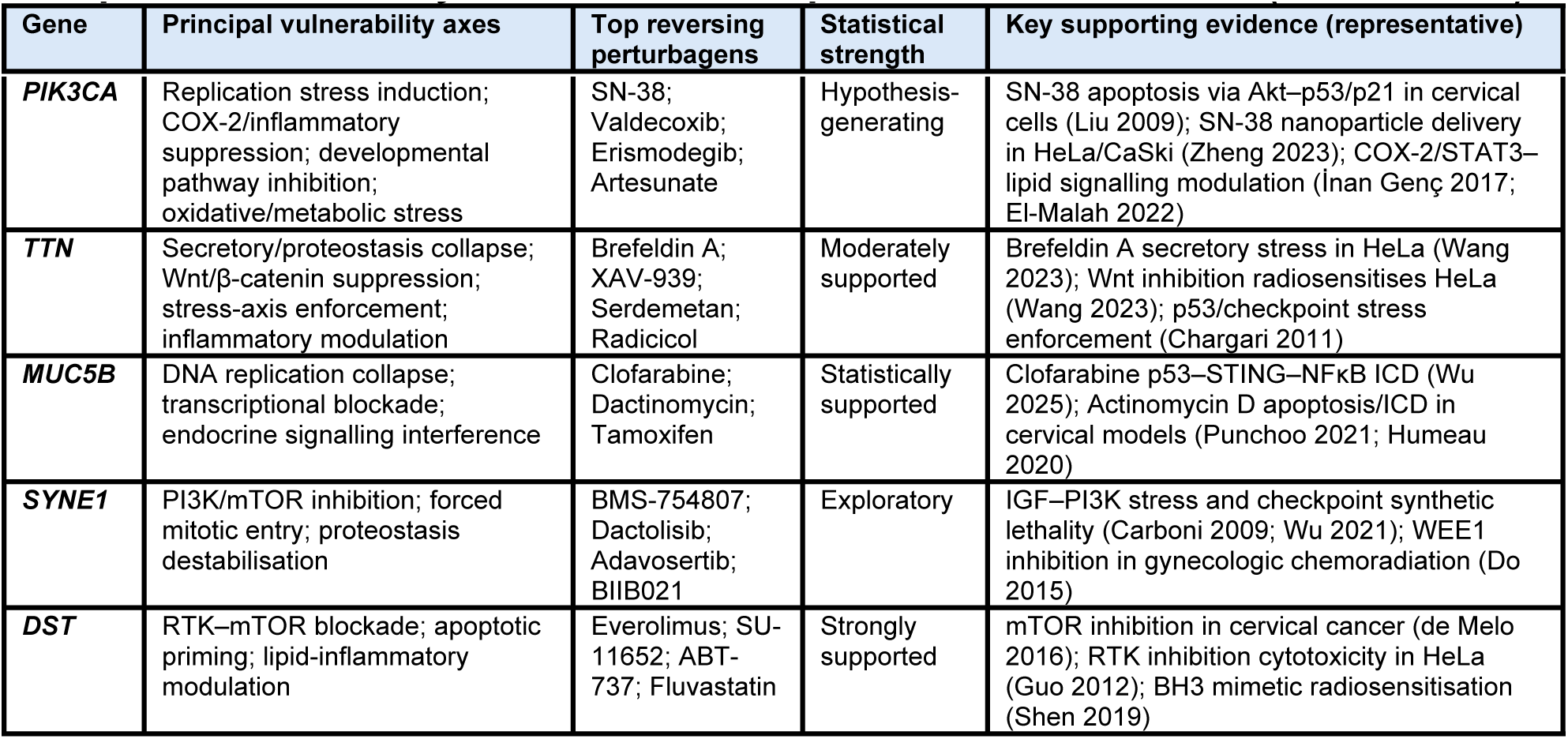
Proteome-anchored connectivity mapping defines gene-specific therapeutic vulnerability axes in cervical squamous cell carcinoma (TCGA-CESC)

### Prospective Synergy Hypotheses for Cervical Cancer Emerging from Connectivity-Defined Proteomic Vulnerabilities

Connectivity-defined reversal signatures nominate tumour-state liabilities that are unlikely to be optimally exploited by single agents. Instead, the most coherent translational next step is to formulate prospective, mechanism-anchored synergy hypotheses that (i) align with the dominant vulnerability axes encoded within each gene-defined programme and (ii) can be tested using tractable pharmacodynamic readouts in cervical cancer models. Importantly, these hypotheses are grounded not only in the statistical connectivity mapping results but also in the regulatory architectures inferred from transcription factor network analysis. As illustrated in **Figure 17**, each mutation-defined tumour state is organised around a distinct transcriptional backbone that coordinates stress responses, proliferative signalling, and survival buffering. For example, TTN-associated states centre on TP53-anchored stress adaptation and proteostasis control; PIK3CA states are structured around RB1–E2F replication and transcriptional regulation; SYNE1 states display MYC/E2F-driven proliferation with attenuation of key checkpoint regulators; DST programmes are dominated by EGFR-linked RTK signalling coupled to survival and apoptotic buffering; and MUC5B states integrate MYCN-dependent transcriptional throughput with RELA/NF-κB inflammatory signalling. These regulatory frameworks provide mechanistic context for the connectivity-derived perturbagens positioned within each network, highlighting where pharmacologic perturbation is predicted to collapse key survival circuits. Importantly, the structure of these networks also reveals complementary intervention points—such as replication stress, survival signalling, proteostasis control, mitochondrial apoptosis, and immune activation—that naturally lend themselves to combination strategies targeting orthogonal stress-buffering systems. Accordingly, the sections below outline a series of prospective synergy hypotheses that arise from these integrated multi-omic and regulatory analyses. Where statistical support is modest (e.g., PIK3CA Q ≈ 0.296; SYNE1 Q ≈ 0.584), the combinations should be interpreted as hypothesis-generating prioritisation candidates rather than efficacy claims, providing a structured framework for pursuing experimental validation in cervical cancer models.

**Figure 17.**
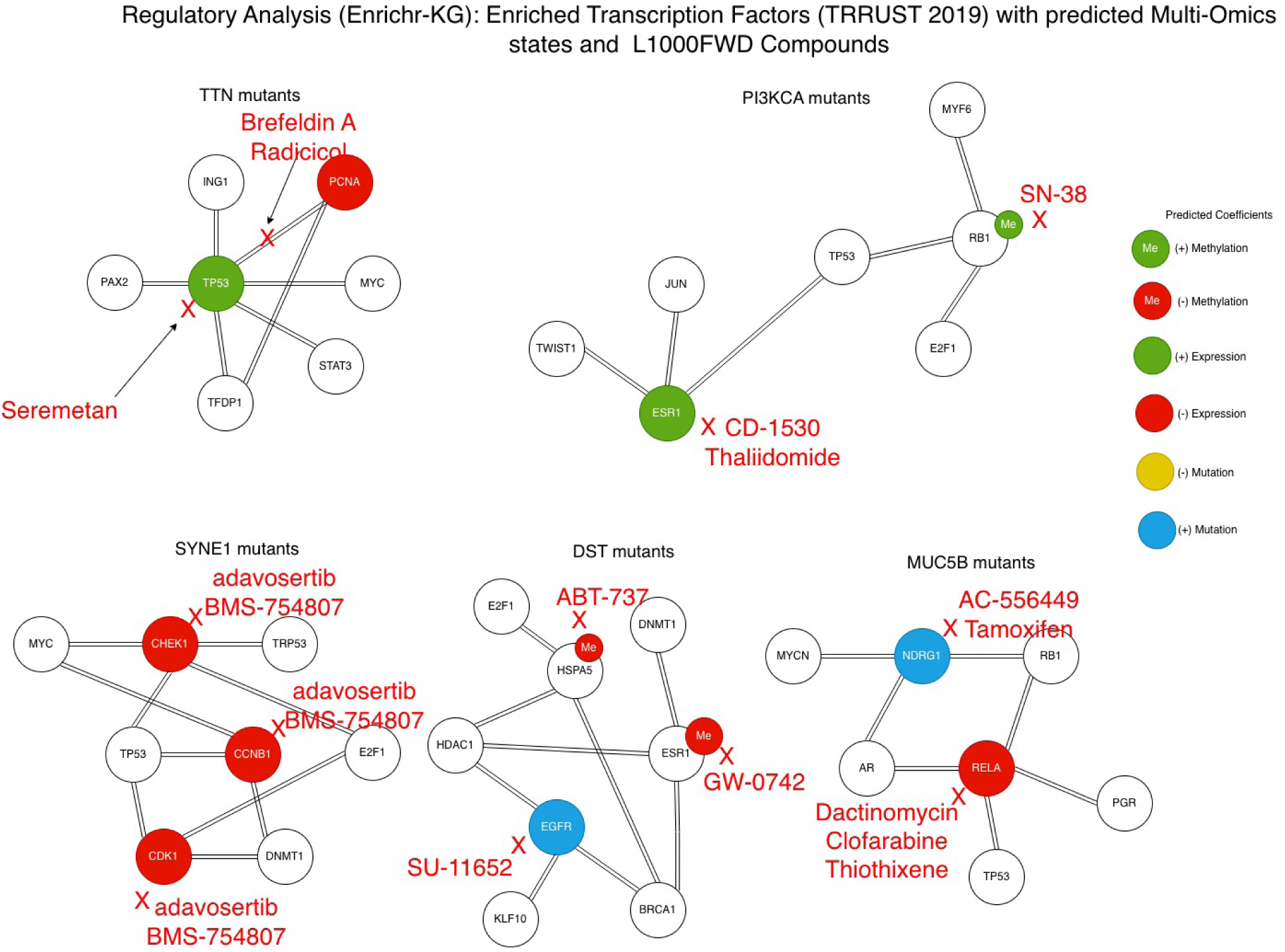
Transcription factor regulatory networks for gene-defined cervical cancer states with predicted reversing compounds. Regulatory network analysis was performed using Enrichr-KG transcription factor enrichment, integrating multi-omic gene states with L1000FWD connectivity-derived perturbagens. Each panel depicts enriched transcription factors and interacting regulatory nodes for tumours stratified by mutations in TTN, PIK3CA, SYNE1, DST, and MUC5B. Nodes represent transcription factors or interacting regulatory proteins, while edges indicate known regulatory or functional interactions. The resulting networks highlight distinct regulatory backbones across gene-defined tumour states, including TP53-centred stress buffering in TTN tumours, ESR1/RB-linked transcriptional control in PIK3CA states, checkpoint-fragile MYC–E2F cell-cycle regulation in SYNE1 tumours, EGFR-driven RTK signalling in DST tumours, and NF-κB-dominated inflammatory transcription in MUC5B tumours, providing mechanistic context for the connectivity-derived therapeutic hypotheses. Node colours represent multi-omics annotations: green, positive expression; red, negative expression; green “Me”, positive methylation; red “Me”, negative methylation; yellow, negative mutation; and blue, positive mutation. “+” indicates positive coefficients and “−” indicates negative coefficients. Red labels, arrows, and crosses indicate predicted pathway-level sites of perturbation by reversing compounds.

#### Replication Stress Coupled to PI3K–AKT–mTOR Survival Collapse

The regulatory architectures inferred from the transcription factor networks **(Figure 17)** suggest a mechanistic convergence between replication control and survival signalling pathways across mutation-defined molecular states. In the PIK3CA-associated state, the network is organised around an RB1–E2F regulatory axis, a canonical driver of DNA replication and S-phase progression. In contrast, the SYNE1-associated state is characterised by suppression of key checkpoint regulators—including CHEK1, CDK1, and CCNB1—within a broader MYC/E2F proliferative framework. Together, these regulatory structures indicate a tumour context in which replication programmes remain active while checkpoint buffering is weakened, creating a cellular environment that is highly dependent on compensatory survival signalling. Within this framework, a central therapeutic hypothesis emerges: topoisomerase-I–induced replication stress may become lethal when PI3K–AKT–mTOR survival signalling is simultaneously suppressed. In the PIK3CA state, the presence of active replication machinery combined with PI3K pathway dependence suggests that pharmacological disruption of survival signalling would compromise the tumour’s ability to tolerate replication-associated DNA damage. Similarly, the SYNE1 checkpoint-attenuated state may further exacerbate replication vulnerability, as defective checkpoint enforcement limits the ability of tumour cells to stabilise replication forks and coordinate DNA repair. Support for this mechanistic model is provided by multiple cross-tumour studies investigating combinations of topoisomerase-I inhibitors and PI3K–mTOR pathway inhibitors. In colorectal cancer models, it has been demonstrated that the mTOR inhibitor everolimus synergises with irinotecan/SN-38, producing enhanced anti-proliferative activity in vitro and superior tumour growth inhibition in vivo under pharmacokinetically optimised dosing regimens (Bradshaw-Pierce et al., 2013). Notably, the strongest responses were observed in PIK3CA- and BRAF-mutant tumours, where metabolomic profiling revealed coordinated suppression of glycolysis by everolimus together with lipid accumulation induced by irinotecan, indicating simultaneous disruption of metabolic survival pathways and replication-associated cytotoxic stress. Further evidence comes from studies using catalytic mTOR inhibitors that suppress both mTORC1 and mTORC2 activity. It has been reported that combining low-dose irinotecan with dual mTOR inhibitors such as AZD2014 or AZD8055 produced markedly greater suppression of tumour growth, migration, invasion, and metastatic dissemination than either treatment alone in colon cancer models, including orthotopic and patient-derived xenografts (Reita et al., 2019). While single-agent irinotecan or mTOR inhibition showed limited impact on metastatic behaviour, the combination nearly abolished metastatic spread, highlighting how collapse of the PI3K/AKT/mTOR/HIF-1α axis fundamentally alters tumour tolerance to replication stress. Evidence specific to cervical cancer further supports this mechanistic interaction. SN-38, the active metabolite of irinotecan, suppresses proliferation and induces apoptosis in HeLa and SiHa cervical cancer cells through an Akt–p53/p21 regulatory axis, with partial rescue of cytotoxicity observed following forced Akt activation (Liu et al., 2009). These findings directly demonstrate that PI3K–Akt signalling modulates cellular sensitivity to replication-associated DNA damage in cervical cancer systems. Additional work using targeted SN-38 delivery strategies has further confirmed the feasibility of exploiting topoisomerase-I–induced stress in cervical cancer models (Zheng et al., 2023). Taken together, these observations support a unified mechanistic model in which replication damage and PI3K–AKT–mTOR survival signalling act as complementary stress-buffering systems. Disruption of both processes simultaneously—by imposing replication stress with topoisomerase-I inhibition while pharmacologically collapsing PI3K–mTOR signalling—produces supra-additive antitumour effects. The regulatory networks identified in **Figure 17** provide a molecular context for this interaction: replication programmes driven by RB1–E2F activity operate alongside weakened checkpoint control but sustained survival signalling, creating a state in which tumour cells are particularly vulnerable to combined replication stress and PI3K pathway inhibition. Pharmacodynamic validation of this hypothesis would involve monitoring markers of replication stress (γH2AX, phosphorylated RPA, replication fork instability), suppression of PI3K pathway output (pAKT, pS6), activation of DNA damage response pathways (p53/p21 induction), and execution of apoptotic programmes (cleaved caspase-3). Such markers provide a mechanistically grounded framework for evaluating SN-38–based combination therapies in PIK3CA- and SYNE1-stratified cervical cancer models.

#### Replication Stress Amplified by Oxidative and Mitochondrial Destabilisation

The transcription factor regulatory networks shown in **Figure 17** further suggest that replication-associated vulnerabilities may converge with oxidative and mitochondrial stress pathways across several mutation-defined molecular states. In particular, the TTN-associated network is organised around stress-adaptive regulators including TP53, MYC, and STAT3, which are central mediators of cellular responses to metabolic imbalance, oxidative stress, and proteotoxic pressure. In contrast, the PIK3CA and MUC5B networks link replication-associated transcriptional programmes—driven by RB1–E2F and MYCN regulatory axes—to inflammatory and metabolic signalling nodes. Together, these regulatory architectures indicate a tumour context in which active replication programmes depend on coordinated redox control, mitochondrial function, and proteostasis buffering to tolerate intrinsic stress. Within this framework, a complementary therapeutic hypothesis emerges: replication-associated DNA damage becomes more lethal when combined with oxidative or mitochondrial stress that erodes the tumour’s intrinsic stress-buffering capacity. Connectivity mapping analyses repeatedly nominate compounds that either induce replication stress (e.g., the SN-38/topoisomerase-I axis) or impose mitochondrial and reactive oxygen species (ROS) stress, suggesting that these perturbational signatures converge on vulnerabilities in redox regulation and checkpoint tolerance. Mechanistic evidence strongly supports this interaction; it has been demonstrated that artemisinin and artesunic acid derivatives can directly engage topoisomerase I, particularly when engineered as hybrid molecules incorporating camptothecin or SN-38 pharmacophores (Botta et al., 2020). These compounds inhibited human topoisomerase-I activity in vitro and exhibited superior anticancer activity compared with camptothecin or paclitaxel alone. Structural modelling further revealed that the sesquiterpene endoperoxide scaffold localises near the camptothecin binding cleft, providing a mechanistic explanation for how replication-associated DNA damage can be coupled to endoperoxide-driven radical generation. This dual mechanism supports the hypothesis that combining SN-38 with artemisinin-class agents could simultaneously intensify replication stress while overwhelming tumour redox defences. The biological plausibility of this model is reinforced by extensive evidence linking mitochondrial ROS generation, endoplasmic reticulum stress, and redox imbalance to cellular responses to chemotherapeutic injury. A study reported that cyclosporin A (CsA) induces pronounced ER stress, mitochondrial ROS production, lipid peroxidation, and redox imbalance, activating signalling pathways including MAPK, NF-κB, TGF-β, and Nrf2 (Wu et al., 2018). Importantly, CsA-induced oxidative stress can also trigger compensatory autophagy, highlighting how ROS–autophagy coupling functions as a dynamic stress-buffering axis that partially mitigates cytotoxic damage. In the context of TTN-associated molecular states, where redox regulation, proteostasis, and mitochondrial adaptation appear prominent, this biology provides a mechanistic framework for how oxidative stressors may synergise with replication stress: by pushing tumour cells beyond their capacity to activate protective autophagy and checkpoint responses, replication damage becomes irrecoverable rather than tolerated. Evidence in cervical cancer systems further supports this interaction. Artesunate has been shown to induce apoptosis in SiHa cervical cancer cells through increased ROS production and Ca²⁺ influx, mitochondrial membrane depolarisation, and a pro-apoptotic shift in Bcl-2 family proteins (Sadana et al., 2025). These mitochondrial and oxidative effects align with metabolic and transport features inferred in PIK3CA-associated states and with the replication-stress tolerance programmes observed in MUC5B networks, suggesting that mitochondrial destabilisation may act as a critical second axis of vulnerability alongside replication damage. Collectively, these observations support a unified model in which replication-associated DNA damage is potentiated by oxidative and mitochondrial stress that disables redox buffering, autophagy-mediated adaptation, and checkpoint recovery mechanisms. The regulatory architectures observed in **Figure 17** reinforce this concept by showing that tumour survival depends on coordinated stress-response networks integrating replication control, metabolic adaptation, and redox homeostasis. Disrupting these buffering systems simultaneously—by imposing replication stress through topoisomerase-I inhibition while inducing oxidative or mitochondrial destabilisation—may therefore produce supra-additive cytotoxic effects. Pharmacodynamic validation of this hypothesis would involve monitoring ROS flux (e.g., DCFDA assays), mitochondrial membrane potential (JC-1 or TMRM), lipid peroxidation, mitochondrial apoptosis markers (BAX translocation and cytochrome-c release), and γH2AX accumulation as a marker of replication-associated DNA damage.

#### Proteostasis and Trafficking Collapse Synergises with Checkpoint and p53-Axis Enforcement

A further coherent synergy hypothesis emerging from TTN- and SYNE1-associated programmes is that acute disruption of proteostasis or secretory trafficking will synergise with enforced checkpoint stress, particularly via the p53 axis, because these tumour states appear sustained by homeostatic buffering rather than dependence on a single oncogenic node. Connectivity mapping nominates both ER–Golgi trafficking disruption (brefeldin A–class compounds) and chaperone-client destabilisation (HSP90 inhibition) as effective routes to collapse these buffered states. This hypothesis is strongly supported by mechanistic evidence linking ER stress to p53-dependent apoptosis. Lin et al. demonstrated that classical ER stress inducers, including brefeldin A, induce robust p53 expression, nuclear localisation, and Ser15 phosphorylation in both MCF-7 and HeLa cells, and that this induction is mediated through NF-κB activation. Importantly, p53 induction was functionally required for brefeldin A–induced apoptosis, as p53 knockdown significantly attenuated cell death. These data establish a direct mechanistic link between trafficking-induced ER stress, NF-κB–dependent p53 activation, and apoptotic execution, providing a strong biological rationale for pairing trafficking collapse with explicit p53-axis enforcement in TTN/SYNE1 states (Lin et al., 2012). Within this framework, enforcing checkpoint stress through pharmacologic p53-axis manipulation further strengthens the collapse of homeostatic buffering. Agents such as serdemetan, developed to disrupt MDM2–p53 interactions, exemplify this strategy by amplifying p53-mediated stress responses and radiosensitisation in solid tumours (Chargari et al., 2011). When combined with ER stressors like brefeldin A, such approaches are predicted to drive tumours past their stress tolerance threshold by simultaneously increasing unfolded protein load and lowering the apoptotic decision point. This logic is particularly relevant to TTN-associated states, which show reliance on stress-buffering, proteostasis, and redox control to sustain proliferative signalling under adverse conditions.

An orthogonal but convergent route to proteostasis collapse is HSP90 inhibition, which destabilises a broad spectrum of oncogenic client proteins, including RTKs, AKT, and cell-cycle regulators. Recent comprehensive analyses highlight that cancer cells exhibit a unique dependence on HSP90, and that HSP90 inhibitors consistently show superior efficacy in combination regimens rather than as single agents, due to synergistic destabilisation of survival networks (Rastogi et al., 2024). Clinical trial experience across tumour types further supports this view: while single-agent HSP90 inhibitors have shown limited activity due to toxicity or resistance, combination strategies—particularly with stress-inducing or checkpoint-targeting therapies—have produced improved outcomes and clearer pharmacodynamic target engagement. These observations align closely with the TTN and SYNE1 connectivity signatures, which point to multi-client signalling dependence rather than single-pathway addiction, making them ideal candidates for network-level destabilisation. Crucially, this strategy is directly feasible in cervical cancer models. Brefeldin A imposes ER–Golgi collapse (Wood et al., 1991), and engineered derivatives show antiproliferative activity with mitochondrial apoptosis signatures in HeLa cells (M. Wang et al., 2023). Likewise, the HSP90 inhibitor BIIB021 exhibits nanomolar cytotoxicity in HeLa with clear evidence of intrinsic apoptosis induction and client protein destabilisation (Güven & Özgür, 2023). Together with the demonstrated requirement for p53 activation in ER stress–induced apoptosis (Lin et al., 2012) and the strong clinical rationale for combination-based HSP90 inhibition (Rastogi et al., 2024), these findings support a unified model in which proteostasis or trafficking collapse is rendered lethal when paired with checkpoint or p53-axis enforcement. Pharmacodynamic validation of this hypothesis would focus on ER stress and unfolded protein response markers (CHOP/DDIT3, ATF4), NF-κB and p53 activation status, depletion of HSP90 client proteins (RTKs, AKT), cell-cycle catastrophe indicators, and downstream apoptotic execution.

The transcription factor networks in **Figure 17** show that TTN tumours are organised around MYC/STAT3-driven stress-adaptive programmes anchored by TP53, while the SYNE1 state combines active MYC–E2F proliferation with reduced expression of key checkpoint regulators (CHEK1, CDK1, CCNB1). These regulatory architectures indicate reliance on proteostasis and homeostatic buffering rather than single-pathway addiction, providing a mechanistic basis for the predicted synergy between proteostasis or trafficking collapse and pharmacologic enforcement of the p53 stress axis.

#### Developmental Pathway Blockade as a Stress Amplifier for Cytotoxic and Radiobiological Injury

A further, strong mechanistic synergy hypothesis emerging from the connectivity-defined vulnerabilities involves developmental pathway blockade as a stress amplifier, particularly in TTN-associated Wnt-dependent states, PIK3CA-associated Hedgehog-linked states, and DST programmes with active RTK–mTOR signalling. The central premise is that developmental signalling pathways such as Wnt/β-catenin and Hedgehog function as stress-adaptive transcriptional backbones that sustain tumour-state resilience under cytotoxic or radiobiological insult; their inhibition is therefore predicted to cooperate with DNA damage–inducing modalities by disabling compensatory survival, EMT, and stemness programmes. This logic is supported by compelling combination evidence from non-cervical solid tumours. In triple-negative breast cancer, Shetti et al. demonstrated that Wnt pathway inhibition with XAV939 synergises with low-dose paclitaxel, achieving apoptosis and tumour regression comparable to high-dose chemotherapy while suppressing β-catenin signalling, EMT markers, and angiogenic programmes both in vitro and in vivo. Mechanistically, the combination reduced Bcl-2, increased caspase-3 and PARP cleavage, suppressed β-catenin and downstream EMT regulators, and restored E-cadherin expression, indicating that Wnt blockade sensitises tumour cells to cytotoxic stress by dismantling EMT and survival circuitry (Shetti et al., 2015). Although this study focused on breast cancer, the mechanistic principles are directly relevant to TTN-associated cervical cancer states, which show dependence on stress-buffered signalling and developmental wiring. Parallel support for this concept comes from Hedgehog pathway targeting in stemness-driven malignancies. Sharma et al. showed that combined inhibition of PI3K/Akt/mTOR and Hedgehog signalling using an SMO inhibitor (NVP-LDE225) together with PI3K/mTOR inhibition produced superior suppression of tumour growth and cancer stem cell properties in pancreatic cancer models. This combination cooperatively suppressed GLI transcriptional activity, reduced pluripotency-maintaining factors (Nanog, Oct4, Sox2, c-Myc), inhibited EMT regulators (Snail, Slug, Zeb1), and impaired spheroid formation and tumour growth in vivo (Sharma et al., 2015). These findings provide a mechanistic template for PIK3CA-associated cervical cancer states, where Hedgehog-linked developmental wiring may cooperate with PI3K–mTOR survival signalling to sustain tumour fitness under stress. In this context, simultaneous disruption of Hedgehog/SMO signalling (e.g., erismodegib/LDE225-class inhibitors) and PI3K–mTOR pathways is predicted to erode transcriptional plasticity and stress tolerance, rendering tumours more susceptible to cytotoxic or radiobiological insult, consistent with cooperative effects observed across solid-tumour models (Pan et al., 2010; D’Amato et al., 2014). Importantly, this synergy hypothesis is directly anchored to cervical cancer biology by experimental evidence showing that Wnt/β-catenin inhibition with XAV939 radiosensitises HeLa cells, increasing apoptosis and γH2AX foci while suppressing β-catenin effectors and cell-cycle regulators (Wang et al., 2023). Together, these data support a unified model in which developmental pathway inhibitors act as stress multipliers rather than stand-alone cytotoxics, disabling EMT, stemness, and transcriptional adaptability and thereby amplifying the efficacy of DNA damage–inducing therapies. Pharmacodynamic validation of this hypothesis would focus on suppression of developmental outputs (β-catenin nuclear localisation; AXIN2, c-Myc, cyclin D1; GLI1/GLI2 transcription), coupled with enhanced DNA damage signalling (γH2AX), apoptosis induction, and reduced clonogenic survival under stress. The transcription factor networks in **Figure 17** reveal that TTN tumours are organised around MYC- and STAT3-driven transcriptional programmes compatible with Wnt-dependent signalling, while PIK3CA and DST states integrate developmental wiring with PI3K/RTK survival pathways. These regulatory architectures indicate that developmental signalling contributes to tumour stress tolerance and transcriptional plasticity, providing a mechanistic basis for the predicted synergy between developmental pathway blockade and cytotoxic or radiobiological therapies.

#### Immunogenic Cell Death Induction Coupled to Microenvironmental Clearance Modulation

A distinct and conceptually important synergy hypothesis emerging from the MUC5B-associated programme centres on the coupling of immunogenic cell death (ICD) induction with microenvironmental clearance modulation. The inferred MUC5B state is characterised by replication-stress tolerance and high transcriptional output, suggesting that tumour survival depends not only on DNA repair capacity but also on sustained transcription–translation throughput. In this setting, agents that disrupt transcription and provoke pre-mortem stress signalling are predicted to induce ICD, but their full antitumour impact may depend on efficient processing and clearance of dying cells. This hypothesis is strongly supported by mechanistic work demonstrating that inhibition of RNA synthesis is a common initiating event across pharmacological ICD inducers. Using an AI-guided screen of ∼50,000 compounds, Humeau et al. identified dactinomycin (actinomycin D) as a potent ICD inducer that mediates immune-dependent tumour control in vivo, and showed that transcriptional inhibition—with secondary translation shutdown and eIF2α phosphorylation—represents a shared mechanistic leitmotif across diverse ICD stimulators (Humeau et al., 2020). This directly supports the appearance of dactinomycin as a top reversing perturbagen in the MUC5B connectivity results and provides a unifying explanation for why transcriptional blockade emerges as a vulnerability in this state. Importantly, dactinomycin also has demonstrated direct cervical cancer relevance as an apoptosis inducer in SiHa cells (Punchoo et al., 2021), anchoring this ICD logic within disease-relevant experimental systems. Within the same framework, clofarabine extends this hypothesis by coupling replication stress and transcriptional disruption to innate immune activation. Clofarabine has been shown to induce a p53–STING interaction and activate non-canonical STING–NFκB signalling, transcriptionally upregulating immune-relevant genes while engaging apoptosis and pyroptosis and enhancing CD8⁺ T-cell cytotoxicity (Wu et al., 2025). Together with the transcription-centric ICD mechanism defined by Humeau et al., this supports a model in which replication/transcription collapse in MUC5B-positive tumours can generate immunogenic death signals, but the durability of the antitumour response will depend on how effectively dying cells are processed within the tumour microenvironment. In this context, the emergence of thiothixene as a strong reversing hit provides a mechanistically complementary axis. Kojima et al. demonstrated that thiothixene potently stimulates macrophage efferocytosis, enhancing the continual clearance of apoptotic and lipid-laden cells by both mouse and human macrophages. Mechanistically, thiothixene relieves dopaminergic suppression of efferocytosis and promotes a Stra6L–arginase 1 axis that sustains phagocytic capacity (Kojima et al., 2025). Although thiothixene is not a conventional oncology agent, this biology provides a plausible route by which ICD-inducing therapies could be functionally amplified: by improving the clearance and processing of dying tumour cells, thiothixene-like proefferocytic modulation may shape downstream immune tone, antigen presentation, and resolution of tumour debris, thereby increasing the likelihood that cytotoxic stress translates into durable immune engagement rather than immunologically silent cell death. Taken together, these findings support a two-step synergy model for the MUC5B programme: (i) induction of immunogenic tumour cell death via replication stress and transcriptional shutdown (clofarabine; dactinomycin/actinomycin D), consistent with ICD mechanisms driven by RNA synthesis inhibition (Humeau et al., 2020), followed by (ii) microenvironmental clearance modulation through enhanced efferocytosis (thiothixene), as demonstrated by dopamine-regulated macrophage phagocytic control (Kojima et al., 2025). Pharmacodynamic validation of this hypothesis would focus on ICD hallmarks (calreticulin exposure, HMGB1 release), STING/NFκB-dependent transcriptional outputs (e.g., CXCL10/CCL5-like signatures), and functional phagocytosis/efferocytosis assays in tumour–macrophage co-culture systems, providing a coherent and testable framework for MUC5B-stratified cervical cancer models. Consistent with this hypothesis, the transcription factor network for the MUC5B-associated state in **Figure 17** is centred on MYCN-driven transcriptional output and RELA/NF-κB–mediated inflammatory signalling, indicating a tumour programme characterised by high transcriptional throughput and replication stress tolerance. These regulatory features provide a mechanistic basis for the predicted synergy between transcriptional inhibitors that induce immunogenic cell death and agents that modulate microenvironmental clearance processes.

#### RTK–mTOR Suppression Combined with Mitochondrial Apoptotic Priming

A particularly strong and mechanistically grounded synergy hypothesis emerges from the DST-associated state (Q ≈ 0.005–0.011), which is characterised by coordinated RTK–AKT–mTOR pathway activation alongside apoptotic buffering. In this context, simultaneous RTK–mTOR blockade and mitochondrial apoptotic priming is predicted to produce supra-additive cytotoxicity by collapsing survival signalling while lowering the apoptotic threshold. This logic is strongly supported by cross-cancer mechanistic evidence demonstrating that mTOR inhibition can directly sensitise tumours to BH3 mimetics. In triple-negative breast cancer, Li et al. showed that combining BH3 mimetics with mTOR inhibitors resulted in marked apoptosis in vitro and superior tumour regression in xenograft models compared with either agent alone. Mechanistically, mTOR inhibition suppressed the anti-apoptotic protein MCL-1 while inducing FOXO3a-dependent upregulation of PUMA, thereby facilitating release and activation of core mitochondrial apoptotic effectors (BIM, BAX, BAK) and sensitising cells to BH3-mimetic activity (Li et al., 2018). This mechanism provides a compelling biological rationale for the DST programme, which displays both strong mTOR pathway engagement and apoptotic buffering at the proteomic level. Within cervical cancer–relevant contexts, this synergy logic is further reinforced by direct experimental evidence. Everolimus, an mTOR inhibitor, has demonstrated activity in cervical cancer models through suppression of PI3K–AKT–mTOR signalling and sensitisation to standard therapies (de Melo et al., 2016). Complementarily, the BH3 mimetic ABT-737 radiosensitises cervical cancer cell lines by inducing mitochondrial depolarisation, reactive oxygen species accumulation, and caspase-dependent apoptosis (Shen et al., 2019), consistent with relief of apoptotic buffering. Upstream survival input can also be targeted at the receptor level: SU-11652, a multitarget RTK inhibitor, induces cytotoxicity in cervical carcinoma cells, including HeLa, through disruption of RTK-driven survival signalling and lysosomal destabilisation (Guo et al., 2012; Ellegaard et al., 2013). Taken together, the convergence of (i) statistically robust DST connectivity signals, (ii) cervical cancer–specific evidence for mTOR inhibition and BH3 mimetic efficacy, and (iii) cross-tumour mechanistic validation that mTOR inhibition downregulates MCL-1 while upregulating PUMA (Li et al., 2018), strongly supports RTK–mTOR blockade plus mitochondrial apoptotic priming as a high-priority, biologically coherent combination strategy for DST-positive cervical cancer states. Pharmacodynamic validation would centre on suppression of mTOR outputs (pS6, p4EBP1), BH3 profiling and mitochondrial priming shifts, caspase activation, and clonogenic survival under stress. From **Figure 17**, the DST regulatory network links predicted RTK inhibition (SU-11652) and BH3-mimetic activity (ABT-737) to key nodes within a survival-signalling transcriptional architecture centred on EGFR and stress-adaptation regulators, supporting a model in which RTK–mTOR pathway suppression collapses survival signalling while BH3 mimetics lower the mitochondrial apoptotic threshold, producing synergistic tumour cell killing.

### Future Work: Development of an In Silico, State-Stratified Synergy Discovery and Validation Pipeline

An important next step emerging from this work is the development of a state-stratified, in silico synergy discovery and validation pipeline tailored to cervical cancer, designed to rigorously prioritise the mechanistically grounded combination hypotheses proposed here. While classical synergy metrics such as Bliss independence are widely used for initial screening, they lack statistical rigor and frequently yield false-positive interaction calls because response variability and dose-dependent uncertainty are not modelled (Zhao et al., 2014). To address this limitation, future analyses will incorporate response-surface–based synergy models that enable formal statistical testing of interaction effects and identification of significant synergy regions across full dose–response matrices (Zhao et al., 2014). In parallel, Loewe additivity–based frameworks will be implemented using explicit null reference formulations, which have been shown to reduce bias and mean squared error relative to traditional isobole-based approaches, particularly in settings where the Loewe Additivity Consistency Condition is violated, as is common in high-throughput combination screens (Yadav et al., 2015; Lederer et al., 2018). Complementing these approaches, the zero interaction potency (ZIP) model will be used to quantify changes in drug potency in combination relative to single agents, enabling construction of interaction landscapes that capture synergistic and antagonistic dose regions with low false-positive rates (Yadav et al., 2015). Given that comparative evaluations of large oncology drug screens demonstrate substantial discordance between Bliss, Loewe, highest single agent, and ZIP models depending on dose range, maximal response, and EC50 differences (Vlot et al., 2019), the proposed pipeline will prioritise combinations that show concordant synergy signals across multiple models and dose regions, rather than relying on a single metric. Critically, synergy assessment will be performed within proteomic state–defined tumour subsets, allowing quantitative interaction landscapes to be interpreted in direct alignment with the connectivity-defined vulnerabilities (e.g., RTK–mTOR–active DST states or transcriptionally driven MUC5B states), thereby enabling biologically informed prioritisation of combinations for targeted experimental validation and translational follow-up in cervical cancer.

## Conclusion

This study demonstrated that the functional consequences of recurrent somatic alterations in cervical cancer are not reliably captured at the level of mutation frequency or steady-state mRNA abundance, but are more coherently represented at the level of proteomic signalling states. By integrating genomic, epigenomic, transcriptomic, and RPPA-based proteomic data within a unified gene-centric framework, we were able to show that diverse upstream perturbations converge onto distinct and classifiable tumour states that can be robustly inferred using elastic-net modelling. A central finding was the systematic dissociation between molecular layers. While PIK3CA mutation is associated with broad and structured transcriptional and epigenetic reprogramming, other recurrent alterations—including TTN, SYNE1, and DST—are largely transcriptionally silent yet give rise to distinct proteomic signatures. These results support a network propagation model in which tumour phenotypes are encoded through multi-layer integration and emerge most clearly at the level of signalling proteins and pathway activity. This highlights the limitation of transcriptome-centric or mutation-centric stratification strategies and underscores the value of proteome-anchored approaches for functional interpretation. Importantly, not all recurrent mutations exert equivalent functional impact in cervical cancer. TTN displays minimal epigenetic, transcriptomic, and proteomic effects consistent with passenger behaviour, whereas DST and MUC5B define coherent proteomic programmes involving growth-factor signalling, immune modulation, metabolism, and chromatin regulation. These distinctions provide a rational basis for prioritising mutation-associated programmes for downstream investigation. By linking proteomic tumour states to perturbational signatures through connectivity mapping, this framework further enables systematic generation of therapeutic hypotheses for cervical cancer. The observed convergence on stress-buffering and survival systems—including proteostasis, checkpoint control, metabolic adaptation, and inflammatory signalling—suggests that cervical tumours may be more effectively targeted through disruption of adaptive network dependencies rather than single pathway inhibition. However, these predictions should be interpreted as hypothesis-generating and require experimental validation. Overall, this work provides a generalisable strategy for translating multi-omics heterogeneity into mechanistically interpretable and pharmacologically actionable tumour states. Extending this framework to additional recurrent alterations, tumour types, and higher-resolution data modalities (e.g. single-cell and spatial profiling) may further refine state-based cancer classification and support the development of more precise, context-dependent therapeutic interventions.

## Data Availability

All datasets analysed in this study are publicly available through internationally recognised, version-controlled repositories providing open-access genomic and proteomic data. No controlled-access permissions were required for data retrieval or processing. All resources used are reproducible and fully documented in their respective databases.

### TCGA Multi-Omics Data

Multi-omics data for cervical squamous cell carcinoma and endocervical adenocarcinoma (TCGA-CESC), including RNA-seq, DNA methylation (Illumina HumanMethylation450), RPPA proteomics, somatic mutation (MAF) files, and associated clinical metadata, were obtained from the Genomic Data Commons (GDC) portal of The Cancer Genome Atlas (https://portal.gdc.cancer.gov/). All datasets are publicly available and were accessed under open-access conditions.

### Pathway Enrichment Analysis (Reactome-based)

Pathway enrichment analysis was performed using the Reactome pathway database implemented through the Reactome package in R to identify overrepresented biological pathways within user-defined gene sets. Reactome provides a curated and peer-reviewed repository of human signalling and biochemical pathways, enabling functional interpretation of molecular features in a biologically coherent context. Enrichment results were summarised and visualised as bar plots, with pathways ranked according to statistical significance, allowing straightforward identification of dominant signalling and regulatory processes associated with the multi-omics states.

### Protein–Protein Interaction Networks (STRING)

Protein–protein interaction networks were obtained from the STRING (Search Tool for the Retrieval of Interacting Genes/Proteins) database (https://string-db.org/), a comprehensive resource integrating known and predicted physical as well as functional associations between proteins across diverse organisms. STRING aggregates interaction evidence from high-throughput experiments, curated databases, text mining of the literature, co-expression analyses, and evolutionary genomic context, providing confidence-scored associations for large-scale interaction mapping and network analysis in systems biology and functional genomics.

### Transcription Factor Enrichment (TRRUST 2019)

Transcription factor target enrichment analysis was performed using the TRRUST 2019 database via the Enrichr platform (https://maayanlab.cloud/Enrichr/), which contains curated transcriptional regulatory networks derived from literature. This resource enables systematic identification of transcription factors whose known targets are overrepresented in a given gene set, supporting mechanistic interpretation of connectivity or perturbation signatures.

### Perturbational Transcriptional Signatures (L1000FWD)

Perturbational transcriptional signatures used for connectivity mapping were obtained from the LINCS L1000 Fireworks Display (L1000FWD) platform (https://maayanlab.cloud/L1000FWD/), which provides open-access perturbational gene expression profiles generated across multiple human cancer cell lines.

### Drug Annotation and Response Metadata (CTRP v2)

Drug annotation metadata used to contextualise and interpret L1000 perturbagens were derived from the Cancer Therapeutics Response Portal (CTRP v2), an open-access resource providing compound-level annotations, protein target information, and cancer cell line response data.

### Code and Reproducibility

All computational analyses were conducted using R (v4.3 or later) under a reproducible and open-source environment. Scripts used for data integration, preprocessing, model training, and visualisation are available in the associated Tutorial Handbook, which is available upon request.

## Author Contributions

Saltiel Hamese completed this research and wrote the research paper together with the contributions from all the authors as follows: Dr. Mutsa Takundwa, Prof. Earl Prinsloo and Dr. Deepak B. Thimiri Govinda Raj. All the authors have read and approved the manuscript.

## Acknowledgements

This work is funded by the South African National Research Foundation (NRF).

---

**Supplementary Figure S0.**
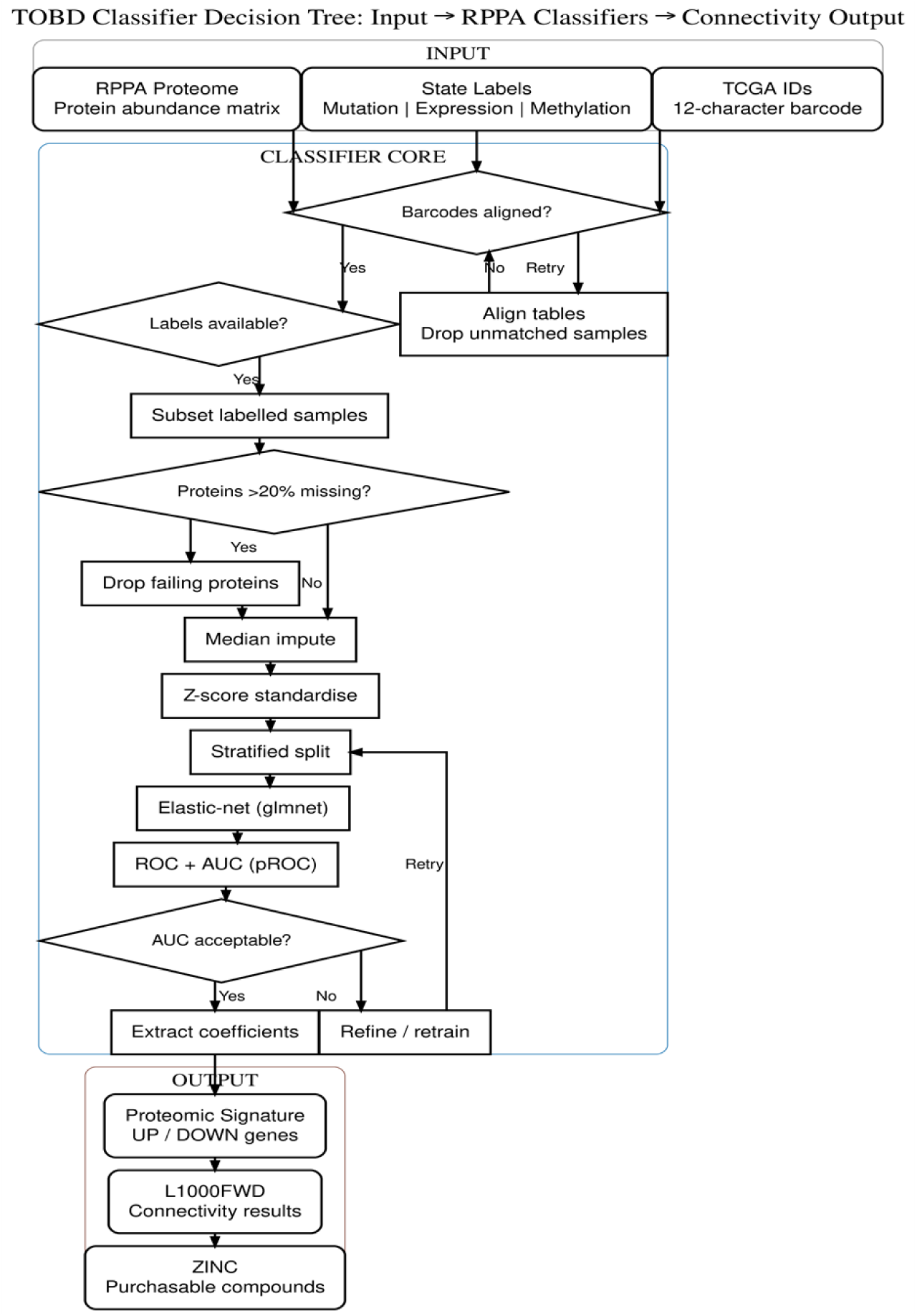
Elastic-net classifier decision tree for proteome-driven functional tumour-state projection. This schematic illustrates the elastic-net classifier workflow used to infer gene-specific molecular activation states directly from RPPA proteomic data across multiple recurrently mutated genes. RPPA protein abundance matrices serve as the sole feature space, while class labels are independently derived from upstream genomic (mutation), transcriptomic (expression), or epigenomic (DNA methylation) layers. Samples are first harmonised using 12-character TCGA barcodes, and only tumours with non-missing state labels are retained. Proteins with >20% missing values are excluded, remaining missing values are median-imputed, and all features are z-score standardised. The resulting samples × proteins matrix is split using stratified sampling and modelled using elastic-net regression (glmnet), with optimisation of the mixing parameter (α) and regularisation strength (λ). Predictive performance is evaluated using receiver operating characteristic (ROC) curves and area under the curve (AUC). Upon satisfactory performance, non-zero coefficients are extracted and mapped to canonical gene symbols to define sparse, directionally signed proteomic signatures. These signatures represent the functional execution of genomic, transcriptomic, or epigenomic tumour states at the protein level and are subsequently used as inputs for L1000FWD connectivity mapping and therapeutic hypothesis generation

**Supplementary Figure S3.**
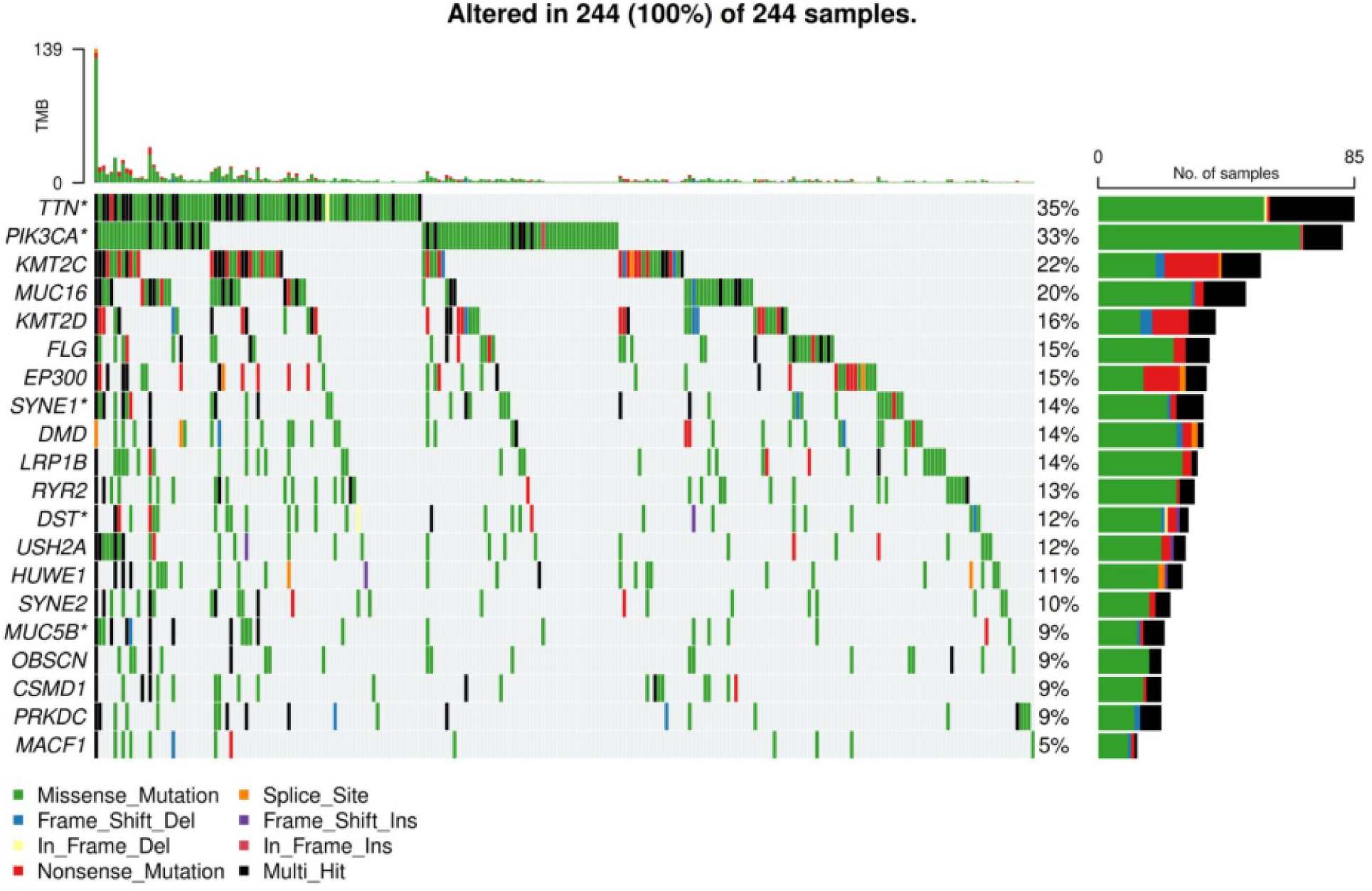
Somatic mutation landscape of methylation-covered genes in TCGA-CESC. Oncoplot showing somatic alterations across 244 TCGA cervical cancer tumours with matched DNA methylation data, restricted to the top 20 most frequently mutated genes with methylation probe coverage. Each column represents an individual tumour and each row represents a gene; mutation types are colour-coded by variant classification. The upper barplot indicates tumour mutation burden per sample, while the right-hand barplot summarises the proportion of tumours harbouring mutations in each gene. Genes marked with an asterisk (*) ***(TTN, PIK3CA, SYNE1, DST, and MUC5B)*** denote genes containing at least one significantly differentially methylated CpG, highlighting loci with concurrent genetic and epigenetic alteration.

**Supplementary Figure S6.**
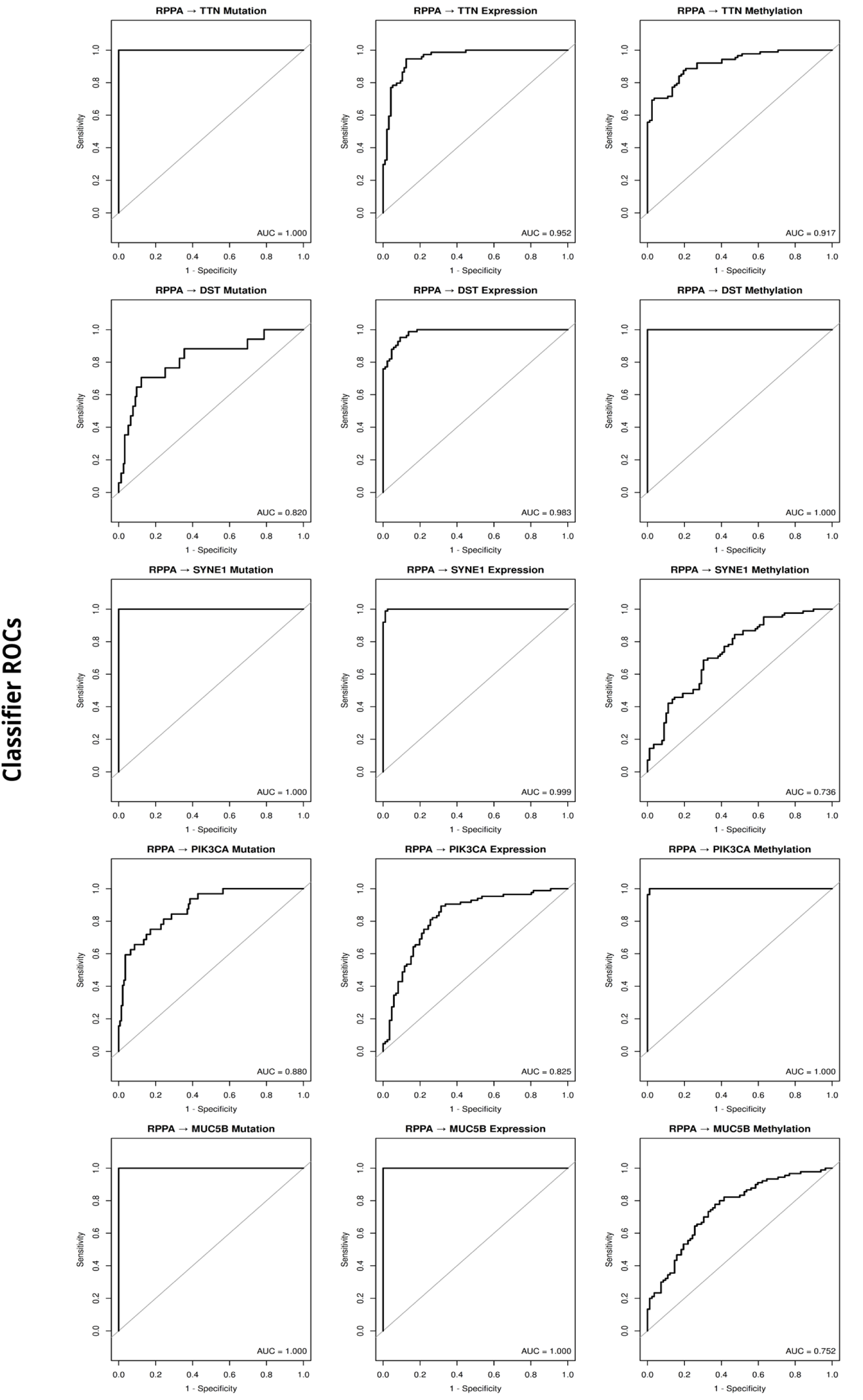
RPPA-based prediction of recurrently mutated gene multi-omics states. Receiver operating characteristic (ROC) curves summarise elastic-net classifier performance using RPPA proteomic features to predict molecular states across genomic, transcriptomic, and epigenomic layers for TTN, MUC5B, DST, SYNE1, and PIK3CA. For TTN, mutation status is perfectly predicted (AUC = 1.000), with high accuracy for expression (AUC = 0.952) and methylation (AUC = 0.917). MUC5B shows perfect discrimination for mutation (AUC = 1.000) and expression (AUC = 1.000) states, while methylation is moderately predictive (AUC = 0.752). DST classifiers achieve excellent accuracy for expression (AUC = 0.983) and methylation (AUC = 1.000), with reduced but robust prediction of mutation (AUC = 0.820). SYNE1 exhibits near-perfect classification of mutation (AUC = 1.000) and expression (AUC = 0.999) states, while methylation is moderately predictive (AUC = 0.736). PIK3CA shows moderate-to-high discrimination for mutation (AUC = 0.880) and robust prediction of expression states (AUC = 0.825), with near-perfect performance for methylation-defined states (AUC = 1.000). These results indicate that RPPA proteomic profiles reliably capture downstream effects of genomic and transcriptomic alterations, with variable propagation of epigenetic changes to protein-level signalling. Grey diagonals indicate no-discrimination baselines. Equivalent RPPA-based classifier performance plots for additional recurrently mutated genes are provided in the Supplementary Material.

**Supplementary Table S1.** Unified gene-centric multi-omics annotation dictionary for TCGA-CESC. This table defines the master gene-level annotation framework used for multi-omics integration across transcriptomic, epigenomic, proteomic, and genomic mutation layers. Each row corresponds to a single gene represented by a stable Ensembl identifier and HGNC-approved gene symbol, and is annotated with genomic coordinates, gene biotype, and explicit indicators of data availability across RNA-Seq, DNA methylation (Illumina HM450), RPPA proteomics, and somatic mutation data. Quantitative summary metrics include the number of mapped CpG probes per gene, RPPA peptide and antibody coverage, total mutation records, number of mutated samples, and the fraction of non-synonymous and high-impact mutations. This dictionary serves as the structural backbone for deterministic multi-omics harmonisation, ensuring consistent gene-level alignment, explicit encoding of modality coverage, and interpretable downstream modelling of molecular tumour states.

| Feature | Description |
| --- | --- |
| Ensembl ID | Ensembl gene identifier used as the stable genomic reference. |
| Gene Symbol | HGNC-approved gene symbol used for cross-modal mapping. |
| Gene Type | Gene biotype (e.g., protein_coding, lncRNA, pseudogene). |
| Chromosome (Chr) | Chromosomal location of the gene. |
| Start | Genomic start coordinate (GRCh38). |
| End | Genomic end coordinate (GRCh38). |
| Strand | Transcriptional strand (+ / -). |
| RNA-Seq Available (has_rnaseq) | Indicates availability of transcriptomic expression data for the gene (TRUE for all entries). |
| Methylation Available (has_methylation) | Indicates whether the gene has at least one mapped CpG probe from the HM450 array. |
| Number of CpG Probes (n_meth_probes) | Total number of CpG probes mapped to the gene, representing epigenomic coverage depth. |
| RPPA Available (has_rppa) | Indicates whether the gene is represented in the RPPA proteomic panel. |
| Number of RPPA Targets (n_rppa_targets) | Number of unique peptide targets mapping to the gene. |
| Number of RPPA Antibodies (n_rppa_agid) | Number of distinct RPPA antibodies measuring the gene product. |
| Mutation Records (mut_n_records) | Total number of somatic mutation calls observed for the gene. |
| Mutated Samples (mut_n_samples) | Number of unique tumour samples carrying at least one mutation in the gene. |
| Fraction Non-synonymous (mut_frac_nonsyn) | Proportion of mutations that are non-synonymous, reflecting functional alteration burden. |
| Fraction High-Impact (mut_frac_high_impact) | Proportion of mutations classified as high-impact (nonsense, frameshift, splice-site, etc.). |

**Supplementary Table S2.**
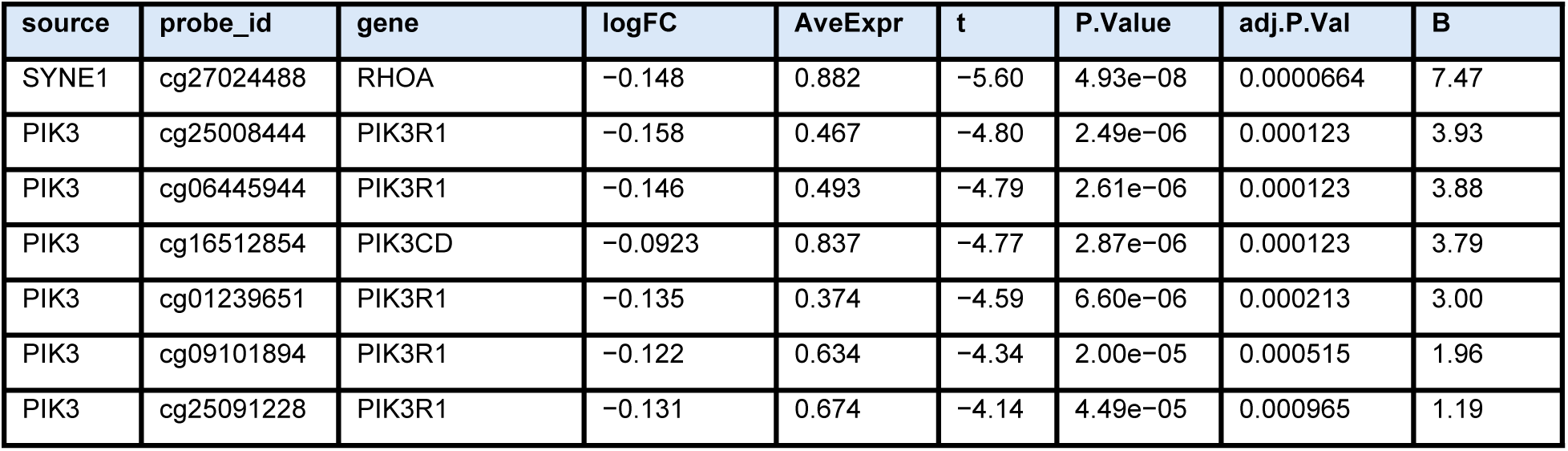
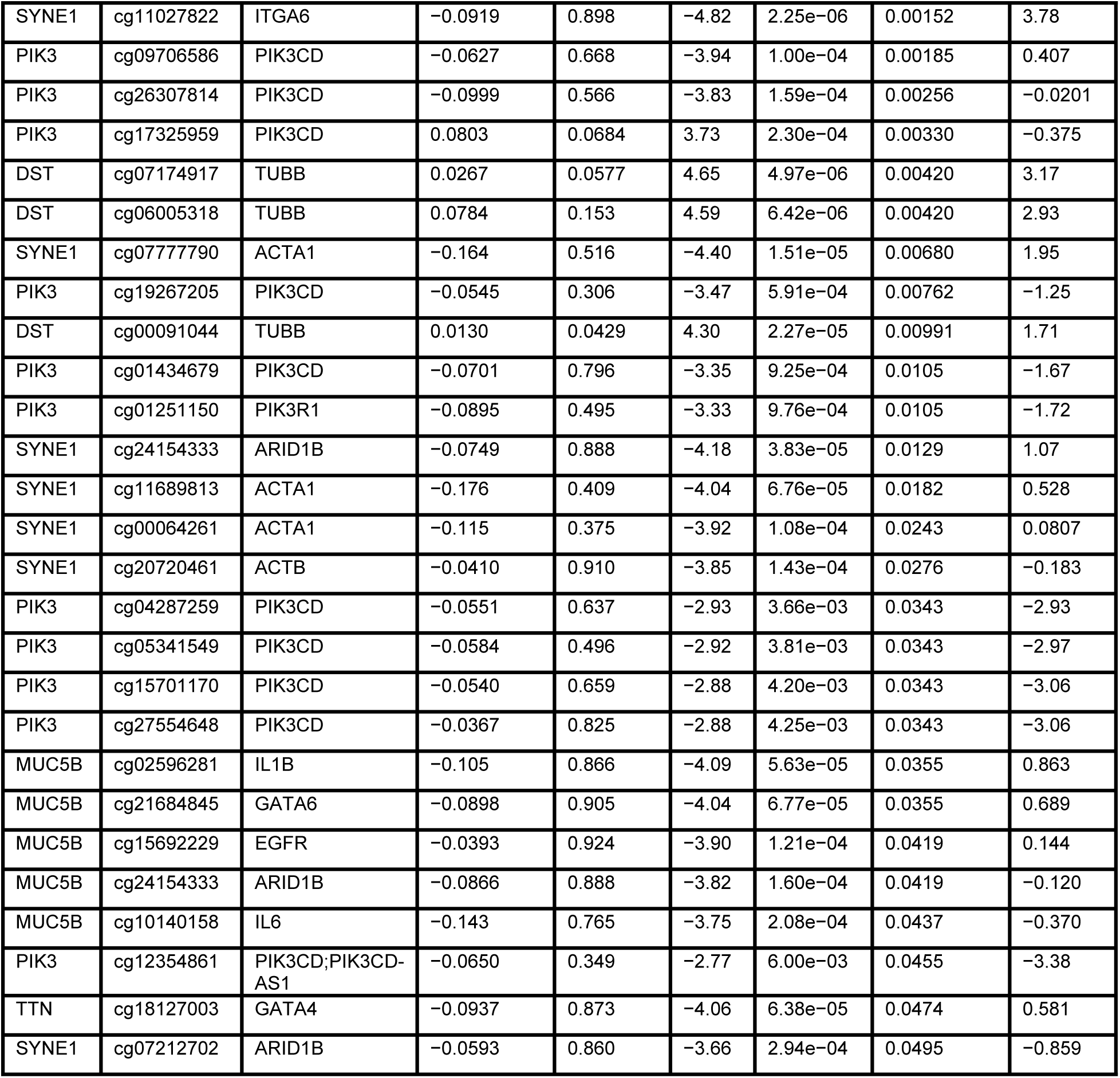
Significant mutation-associated differentially methylated CpG sites across recurrently mutated genes in TCGA-CESC.

**Supplementary Table S4.**
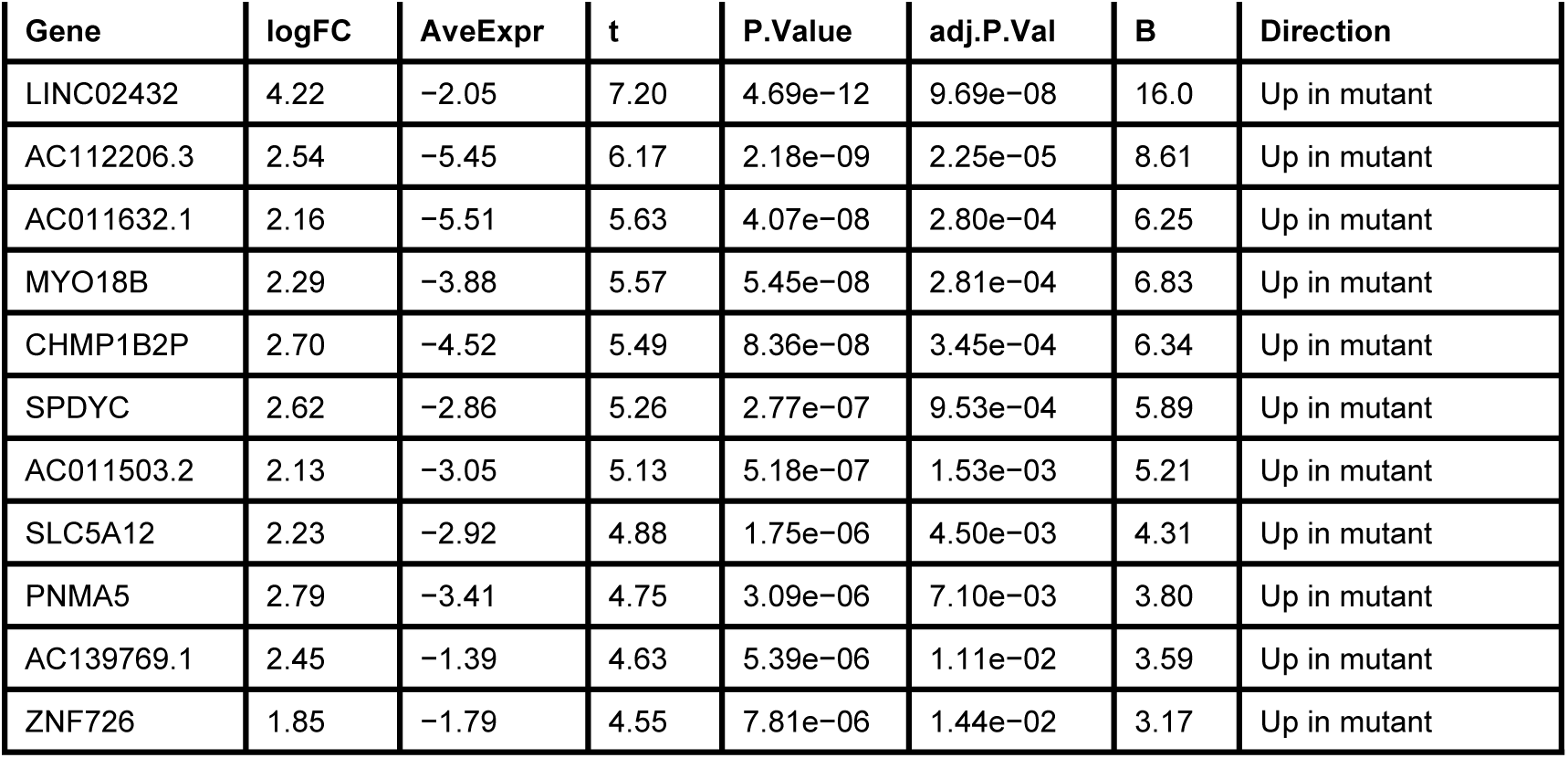
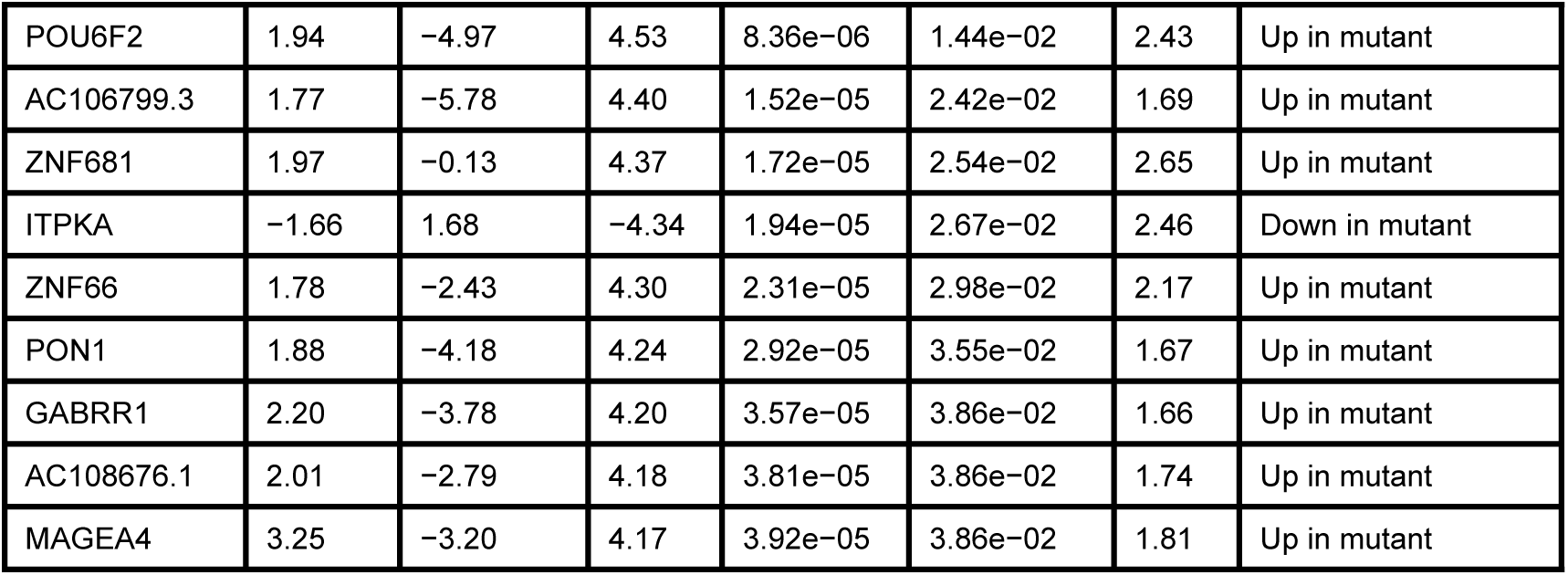
Differential gene expression associated with MUC5B mutation status in cervical cancer. Genome-wide RNA-seq differential expression analysis comparing MUC5B-mutant (n = 21) and wild-type (n = 283) tumours in the TCGA-CESC cohort. Shown are the top significant genes (FDR < 0.05), ranked by adjusted p-value. Positive logFC values indicate higher expression in MUC5B-mutant tumours.

**Supplementary Table S5.** Differential expression of PI3K pathway regulators associated with PIK3CA mutation status in TCGA-CESC. Regulator-restricted RNA-seq differential expression analysis comparing PIK3CA-mutant (n = 79) and wild-type (n = 225) tumours in the TCGA-CESC cohort. Analysis was limited to a curated set of six PI3K pathway regulatory genes. Shown are all genes meeting the significance threshold (FDR < 0.05). Positive logFC values indicate higher expression in PIK3CA-mutant tumours.

| Gene | logFC | AveExpr | t | P.Value | adj.P.Val | B | Direction |
| --- | --- | --- | --- | --- | --- | --- | --- |
| PIK3R1 | 0.454 | 17.8 | 3.67 | 2.85e-04 | 0.00171 | 0.100 | Up in mutant |
| PIK3CD | -0.432 | 17.0 | -3.20 | 0.00153 | 0.00460 | -1.29 | Down in mutant |
| PIK3C3 | -0.193 | 17.6 | -2.93 | 0.00364 | 0.00727 | -2.14 | Down in mutant |

**Supplementary Table S7.** RPPA feature coefficients contributing to PIK3CA-associated proteomic state classification across molecular modalities. Elastic-net–derived RPPA features contributing to expression-, methylation-, and mutation-anchored classifiers for PIK3CA molecular states. Coefficients represent signed feature weights; absolute coefficients indicate relative contribution magnitude within each model.

| Feature ID | Gene Symbol | Coefficient | Absolute Coefficient | Model |
| --- | --- | --- | --- | --- |
| AGID00135 | EEF2 | -0.218 | 0.218 | Expression |
| AGID00066 | SRC | 0.132 | 0.132 | Expression |
| AGID02153 | INPP4B | 0.102 | 0.102 | Expression |
| AGID00255 | SLFN11 | 0.0749 | 0.0749 | Expression |
| AGID00442 | Hexokinase-II | -0.0676 | 0.0676 | Expression |
| AGID00009 | BAK | -0.0567 | 0.0567 | Expression |
| AGID00315 | ATR | -0.0481 | 0.0481 | Expression |
| AGID00209 | Cyclophilin-F | -0.0287 | 0.0287 | Expression |
| AGID00335 | ERALPHA | 0.0244 | 0.0244 | Expression |
| AGID00198 | MCL-1 | -0.0202 | 0.0202 | Expression |
| AGID00421 | VHL | 0.591 | 0.591 | Methylation |
| AGID00060 | RB | 0.534 | 0.534 | Methylation |
| AGID00185 | STAT3 | 0.467 | 0.467 | Methylation |
| AGID00343 | DDB1 | -0.409 | 0.409 | Methylation |
| AGID00408 | ACSS1 | 0.409 | 0.409 | Methylation |
| AGID00134 | NF2 | -0.366 | 0.366 | Methylation |
| AGID00014 | BRAF | 0.362 | 0.362 | Methylation |
| AGID00443 | EIF4EBP1 | -0.336 | 0.336 | Methylation |
| AGID00288 | MCT4 | -0.336 | 0.336 | Methylation |
| AGID00294 | SETD2 | 0.320 | 0.320 | Methylation |
| AGID00031 | Fibronectin | 0.200 | 0.200 | Mutation |
| AGID00319 | Granzyme-B | -0.181 | 0.181 | Mutation |
| AGID00395 | Enolase-2 | 0.161 | 0.161 | Mutation |
| AGID00437 | Cyclin E1 | -0.159 | 0.159 | Mutation |
| AGID00316 | AKT2 | -0.157 | 0.157 | Mutation |
| AGID00408 | ACSS1 | 0.147 | 0.147 | Mutation |
| AGID00284 | ULK1 | 0.140 | 0.140 | Mutation |
| AGID00098 | p38 | -0.135 | 0.135 | Mutation |
| AGID00025 | Cyclin D3 | 0.122 | 0.122 | Mutation |
| AGID00352 | DNA Ligase IV | 0.119 | 0.119 | Mutation |

**Supplementary Table S8.** TTN-associated multi-omics feature coefficients across expression, methylation, and mutation models.

| # | feature | coef | abscoef | gene_symbol | model |
| --- | --- | --- | --- | --- | --- |
| 1 | AGID00235 | 0.270 | 0.270 | EMA | Expression |
| 2 | AGID00050 | 0.248 | 0.248 | P53 | Expression |
| 3 | AGID00131 | 0.242 | 0.242 | HER2 | Expression |
| 4 | AGID00323 | 0.217 | 0.217 | PAK1 | Expression |
| 5 | AGID00018 | -0.216 | 0.216 | CHK2 | Expression |
| 6 | AGID00316 | -0.215 | 0.215 | Akt2 | Expression |
| 7 | AGID00066 | 0.199 | 0.199 | SRC | Expression |
| 8 | AGID00078 | 0.196 | 0.196 | EIF4E | Expression |
| 9 | AGID00224 | -0.188 | 0.188 | PAICS | Expression |
| 10 | AGID00248 | -0.175 | 0.175 | PCNA | Expression |
| 11 | AGID00105 | -0.410 | 0.410 | P27 | Methylation |
| 12 | AGID00086 | 0.240 | 0.240 | GATA3 | Methylation |
| 13 | AGID00492 | 0.216 | 0.216 | PRDX1 | Methylation |
| 14 | AGID00460 | -0.212 | 0.212 | PKCALPHA | Methylation |
| 15 | AGID00318 | 0.158 | 0.158 | Myt1 | Methylation |
| 16 | AGID00142 | -0.158 | 0.158 | HER2 | Methylation |
| 17 | AGID00135 | -0.147 | 0.147 | EEF2 | Methylation |
| 18 | AGID00408 | 0.140 | 0.140 | AceCS1 | Methylation |
| 19 | AGID00149 | -0.118 | 0.118 | TIGAR | Methylation |
| 20 | AGID02141 | 0.113 | 0.113 | G6PD | Methylation |
| 21 | AGID00408 | 1.05 | 1.05 | AceCS1 | Mutation |
| 22 | AGID02160 | 1.02 | 1.02 | SMAD4 | Mutation |
| 23 | AGID02167 | -0.864 | 0.864 | FRS2-alpha | Mutation |
| 24 | AGID00228 | 0.812 | 0.812 | TFAM | Mutation |
| 25 | AGID00326 | 0.799 | 0.799 | ZAP-70 | Mutation |
| 26 | AGID00529 | 0.728 | 0.728 | LKB1 | Mutation |
| 27 | AGID00075 | -0.667 | 0.667 | EPPK1 | Mutation |
| 28 | AGID00128 | -0.661 | 0.661 | SYK | Mutation |
| 29 | AGID00092 | 0.641 | 0.641 | IRS1 | Mutation |
| 30 | AGID00125 | -0.636 | 0.636 | PEA15 | Mutation |

**Supplementary Table S9.** MUC5B-associated multi-omics feature coefficients across expression, methylation, and mutation models.

| # | feature | coef | abscoef | gene_symbol | model |
| --- | --- | --- | --- | --- | --- |
| 1 | AGID00315 | 0.694 | 0.694 | ATR | Expression |
| 2 | AGID00081 | -0.583 | 0.583 | PRAS40 | Expression |
| 3 | AGID00040 | -0.572 | 0.572 | IRS2 | Expression |
| 4 | AGID02151 | -0.571 | 0.571 | FAK | Expression |
| 5 | AGID00048 | -0.522 | 0.522 | NFKBP65 | Expression |
| 6 | AGID00270 | -0.506 | 0.506 | Gli1 | Expression |
| 7 | AGID00133 | 0.487 | 0.487 | XBP1 | Expression |
| 8 | AGID00288 | -0.460 | 0.460 | MCT4 | Expression |
| 9 | AGID00323 | -0.427 | 0.427 | PAK1 | Expression |
| 10 | AGID00321 | -0.422 | 0.422 | Mitofusin-1 | Expression |
| 11 | AGID00397 | 0.143 | 0.143 | Hexokinase-I | Methylation |
| 12 | AGID00179 | 0.0485 | 0.0485 | b-Catenin | Methylation |
| 13 | AGID00149 | -0.0333 | 0.0333 | TIGAR | Methylation |
| 14 | AGID00267 | 0.0157 | 0.0157 | Connexin-43 | Methylation |
| 15 | AGID00369 | -0.0120 | 0.0120 | CD38 | Methylation |
| 16 | AGID00512 | 0.00441 | 0.00441 | PLC-gamma1 | Methylation |
| 17 | AGID00395 | 0.454 | 0.454 | Enolase-2 | Mutation |
| 18 | AGID00447 | 0.384 | 0.384 | CHD1L | Mutation |
| 19 | AGID00156 | 0.352 | 0.352 | NDRG1 | Mutation |
| 20 | AGID00408 | 0.342 | 0.342 | AceCS1 | Mutation |
| 21 | AGID00038 | -0.302 | 0.302 | IGFBP2 | Mutation |
| 22 | AGID00303 | 0.291 | 0.291 | CD171 | Mutation |
| 23 | AGID00397 | -0.278 | 0.278 | Hexokinase-I | Mutation |
| 24 | AGID00029 | 0.275 | 0.275 | ERALPHA | Mutation |
| 25 | AGID00349 | 0.274 | 0.274 | XPF | Mutation |
| 26 | AGID00024 | 0.259 | 0.259 | CYCLINB1 | Mutation |

**Supplementary Table S10.** DST-associated multi-omics feature coefficients across expression, methylation, and mutation models.

| # | feature | coef | abscoef | gene_symbol | model |
| --- | --- | --- | --- | --- | --- |
| 1 | AGID00437 | 0.556 | 0.556 | CYCLINE1 | Expression |
| 2 | AGID00290 | -0.314 | 0.314 | Lasu1 | Expression |
| 3 | AGID00390 | -0.283 | 0.283 | MSH2 | Expression |
| 4 | AGID00426 | 0.248 | 0.248 | TRIP13 | Expression |
| 5 | AGID02179 | -0.237 | 0.237 | SFRP1 | Expression |
| 6 | AGID00267 | 0.226 | 0.226 | Connexin-43 | Expression |
| 7 | AGID00519 | -0.214 | 0.214 | IRF-3 | Expression |
| 8 | AGID00247 | -0.199 | 0.199 | D-a-Tubulin | Expression |
| 9 | AGID00020 | 0.197 | 0.197 | CKIT | Expression |
| 10 | AGID00357 | -0.195 | 0.195 | STING | Expression |
| 11 | AGID00038 | 0.980 | 0.980 | IGFBP2 | Methylation |
| 12 | AGID00264 | 0.817 | 0.817 | ASNS | Methylation |
| 13 | AGID00335 | -0.755 | 0.755 | ERALPHA | Methylation |
| 14 | AGID00182 | 0.727 | 0.727 | p90RSK | Methylation |
| 15 | AGID00088 | -0.669 | 0.669 | YAP | Methylation |
| 16 | AGID00017 | 0.660 | 0.660 | CD31 | Methylation |
| 17 | AGID00109 | 0.652 | 0.652 | TRANSGLUTAMINASE | Methylation |
| 18 | AGID00316 | 0.588 | 0.588 | Akt2 | Methylation |
| 19 | AGID00221 | -0.581 | 0.581 | BiP-GRP78 | Methylation |
| 20 | AGID00014 | 0.560 | 0.560 | BRAF | Methylation |
| 21 | AGID00171 | -0.107 | 0.107 | FASN | Mutation |
| 22 | AGID00300 | 0.102 | 0.102 | PDL1 | Mutation |
| 23 | AGID00026 | 0.0832 | 0.0832 | EGFR | Mutation |
| 24 | AGID02213 | -0.0564 | 0.0564 | FTO | Mutation |
| 25 | AGID00053 | 0.0492 | 0.0492 | PAI1 | Mutation |
| 26 | AGID00092 | 0.0432 | 0.0432 | IRS1 | Mutation |
| 27 | AGID00038 | -0.0152 | 0.0152 | IGFBP2 | Mutation |

